# Constructing a high-density linkage map to infer the genomic landscape of recombination rate variation in European Aspen *(Populus tremula)*

**DOI:** 10.1101/664037

**Authors:** Rami-Petteri Apuli, Carolina Bernhardsson, Bastian Schiffthaler, Kathryn M. Robinson, Stefan Jansson, Nathaniel R. Street, Pär K. Ingvarsson

## Abstract

The rate of meiotic recombination is one of the central factors determining levels of linkage disequilibrium and the efficiency of natural selection, and many organisms show a positive correlation between local rates of recombination and levels of nucleotide diversity indicating that linked selection is an important factor determining genome-wide levels of nucleotide diversity. Several methods for estimating recombination rates from segregating polymorphisms in natural populations have recently been developed. These methods have been extensively used in part because they are relatively simple to implement even in many non-model organisms, but also because they potentially offer higher resolution than traditional map-based methods. However, thorough comparisons of LD and map-based estimates of recombination are not readily available in plants. Here we present a new, high-resolution linkage map for *Populus tremula* and use this to estimate variation in recombination rates across the *P. tremula* genome. We compare these results to recombination rates estimated based on linkage disequilibrium in a large number of unrelated individuals. We also assess how variation in recombination rates is associated with genomic features, such as gene density, repeat density and methylation levels. We find that recombination rates obtained from the two methods largely agree, although the LD-based method identify a number of genomic regions with very high recombination rates that the map-based method fail to detect. Linkage map and LD-based estimates of recombination rates are positively correlated and show similar correlations with other genomic features, showing that both methods can accurately infer recombination rate variation across the genome.

## Introduction

Meiotic recombination (hereafter recombination) is an important evolutionary force that directly alters levels of linkage disequilibrium (e.g. Wright 1931). Recombination therefore has important consequences for how effective natural selection is at removing deleterious mutations or increasing the frequency of beneficial mutations (Felsenstein 1974). Recombination rates are known to vary between species, among individuals within species and among different regions in a genome (Nachman 2002).

Local recombination rates have been shown to be positively correlated with neutral genetic diversity across a wide range of organisms (reviewed in Nachman 2002). One possible explanation for such an association is that cross-over events and/or associated processes, such as gene conversion and double-strand break repair, have direct mutagenic effects and thus act to increase nucleotide polymorphism (e.g. Kulathinal et al. 2008). An alternate explanation is that natural selection has indirect effects on sites linked to a site under selection and therefore also acts to reduce diversity on these sites (Begun and Aquadro 1992). Since recombination breaks down linkage disequilibrium, areas of high recombination are characterized by a rapid decay of linkage disequilibrium and linked selection will hence impact fewer sites in the vicinity of a selected site in these regions (Begun and Aquadro 1992). Conversely, in areas of low recombination rates, linkage disequilibrium will be extensive and indirect selection will impact a larger genomic region. Variation in recombination rates across the genome will generate an association between recombination and sequence diversity. Local variation in recombination rates is therefore an important factor for understanding how natural or artificial selection shapes sequence diversity across the genome of an organism.

Traditionally, recombination rates have been estimated from the relationship of marker positions in linkage maps (Stapley et al. 2017) and more recently recombination rates have also been linked to physical regions of a genome through whole genome sequencing (Nachman 2002). However, producing linkage maps is time consuming and may even be infeasible in some species as it requires controlled crossing of known parents and the establishment of a large segregating progeny population (Stapley et al. 2017). Therefore, methods have been developed that infer recombination rates from linkage disequilibrium (LD) between segregating polymorphisms in individuals sampled from natural populations (e.g. McVean et al. 2004, Chan et al. 2012). Due to the relative ease of obtaining sequence information with modern sequencing methods even from wild populations, these LD-based methods for estimating recombination rates have been widely employed (e.g. McVean et al. 2004, Kulathinal et al. 2008 Silva-Junior and Grattapaglia 2015, Wang et al. 2016, Booker et al. 2017). Detailed knowledge of local variation in recombination rates can be used to infer the action of linked selection by establishing a correlation between the levels of nucleotide diversity and recombination rates across the genome of an organism (McVean et al. 2004, Chan et al. 2012). Using polymorphism data to infer recombination rates and then using these inferred recombination rates to explain variation in genetic diversity could be problematic, but simulations and studies performed using well-established animal model species such as *Mus musculus* (Booker et al. 2017) and in a number of *Drosophila* species (Kulathinal et al. 2008, Chan et al. 2012) suggest that indirect methods for estimating recombination rates are not strongly affected by natural selection. However, comparisons of LD-based and genetic linkage map-based methods for estimating recombination rates are not readily available plant species. Genome structure, and in particular local rates of recombination, show large scale differences between plants and animals (Haenel et al. 2018), and it would therefore also be valuable to assess how well indirect methods for inferring recombination rates perform in plants.

Local variation in recombination rates is known to be associated with a number of different genomic features, such as gene density, repeat density, and cytosine methylation, although the magnitude and direction of these associations are still under debate. Recombination rates have been shown to be both positively and negatively correlated with gene density (positively: e.g. Wang et al. 2016, negatively: e.g. Giraut et al. 2011), GC-content (positively: Kim et al. 2007, negatively: Giraut et al. 2011), repeat density and methylation levels (positively: e.g. Rodgers-Melnick et al. 2015, negatively: Giraut et al. 2011). Characterizing associations between recombination rates and various genomic features at the genus or species level is thus important to avoid making incorrect assumptions about the strength and/or direction of these associations.

The genus *Populus* has emerged as an important model system for forest trees due to its rapid growth rate, ability to generate natural clones and a manageable genome size of ca. 480 Mbp distributed across a haploid set of 19 (2*n*=38) chromosomes (Taylor 2002, Lin et al. 2018). Furthermore, both large and small scale synteny is highly conserved across species in the genus, enabling the transfer of genetic resources between species within the genus (Jansson and Douglas 2007). Further interest in *Populus* has been spurred by their economical (e.g. Taylor 2002) and ecological importance (e.g. Kouki et al. 2004) and over the past two decades a growing number of the ca. 40 species in the genus have been fully sequenced, including *Populus trichocarpa* (Black cottonwood) (Tuskan et al. 2006), *P. euphratica* (Ma et al. 2013) and *P. tremula* (European aspen) (Lin et al. 2018). Similarly, linkage maps have been produced for many of the species in the genus (e.g. Paolucci et al. 2010, Tong et al. 2016), including *Populus tremula* (Zhigunov et al. 2017). However, these maps have been relatively coarse, utilizing a few hundred up to a few thousand markers and typically employing mapping populations consisting of fewer than 300 progenies. Consequently, many of these maps have failed to resolve the expected 19 linkage groups typical for the genus and there is thus a need for developing a high-resolution, fine-scale linkage maps for the whole genus.

*P. tremula* is of special interest within the genus as it has the largest distribution of any tree species in Eurasia, spanning from Spain and Scotland in the west to pacific China and Russia in the east, Iceland and northern Scandinavia in the north to northern Africa and southern China in the south (Luquez et al. 2008). Such extensive geographic distribution means *P. tremula* has adapted to a great variety of different environments, making it a promising species for studying the effects of spatially varying selection and adaptation (e.g. Farmer 1996, Luquez et al. 2008, Wang et al. 2018). To study these effects, it is important to fully understand the genomic landscape of recombination rate variation. Although genome-wide variations in recombination rates have previously been studied in *P. tremula* (e.g. Wang et al. 2016), thus far these analyses have exclusively relied on LD-based methods for estimating recombination rates.

Here we present a newly developed, fine-scale genetic map for *Populus tremula* and use this map to anchor scaffolds from the current draft genome assembly (Potra v1.1, Lin et al. 2018) to chromosomes. We then use this new resource to estimate local variation in recombination rates and use these to assess the correlation with recombination rates inferred from nucleotide polymorphism data from a larger sample of individuals. Finally, we assess how different genomic features, such as gene density, repeat content and methylation levels are associated with the different estimates of local recombination rate.

## Material and methods

### Plant material

In 2013, a controlled F_1_ cross was performed between two unrelated *P. tremula* individuals (UmAsp349.2 x UmAsp229.1) from the Umeå Aspen (UmAsp) collection that consists of c. 300 individuals collected in the vicinity of Umeå in northern Sweden (Fracheboud et al. 2009). This cross yielded 764 full sib progenies that were planted and monitored in a common garden at the Forestry Research Institute of Sweden’s research station in Sävar, 20 km northeast of Umeå (63.9N 20.5E). In addition, we utilized SNP data for 94 individuals of *P. tremula* belonging to the SwAsp collection that consists of 116 individuals sampled from 12 local populations across Sweden (6-10 individuals per population, Luquez et al. 2008). The SNP data has previously been described in Wang et al. (2018) and consists of 4,425,109 SNPs with a minor allele frequency exceeding 5%.

### DNA extraction, sequence capture and genetic map creation

In 2015 leaf samples were collected from all progenies of the F_1_ cross. DNA was extracted using the Qiagen Plant Mini kit according to manufactures guidelines and sent to Rapid Genomics (http://www.rapid-genomics.com) for genotyping using sequence capture probes. The probe set contain 45,923 probes of 120 bases each that were designed to target unique genic regions in the v1.1 *P. tremula* genome assembly (Lin et al. 2018), as well as an additional 70 probes that were designed to specifically target the putative sex determination region on chromosome 19 of the *P. trichocarpa* genome assembly v3.0 (https://phytozome.jgi.doe.gov/pz/portal.html). Parents and all offspring were subjected to sequence capture and subsequently sequenced on an Illumina HighSeq 2000 using paired-end (2×100bp) sequencing to an average depth of 15x per sample. All sequence capture data was delivered from Rapid Genomics in the spring of 2016. In addition, two parents of the F_1_ cross were whole-genome re-sequenced to an average depth of 15x on an Illumina HiSeq 2500 platform with paired-end sequencing (2×150 bp) at the National Genomics Infrastructure at the Science for Life Laboratory in Stockholm, Sweden.

All raw sequencing reads were mapped against the complete *P. tremula* v.1.1 reference genome using `BWA-MEM` v.0.7.12 (Li and Durbin 2009) using default parameters. Following read mapping, PCR duplicates were marked using `Picard` (http://broadinstitute.github.io/picard/) and local realignment around indels was performed using `GATK RealignerTargetCreator` and `IndelRealigner` (McKenna et al. 2010; DePristo et al. 2011). Genotyping was performed using `GATK HaplotypeCaller` (version 3.4-46, (DePristo et al. 2011; Van der Auwera et al. 2013) with a diploid ploidy setting and gVCF output format. `CombineGVCFs` was then run on batches of ~200 gVCFs to hierarchically merge samples into a single gVCF and a final SNP call was performed using `GenotypeGVCFs` jointly on the combined gVCF file, using default read mapping filters.

To obtain informative markers that could be used in the creation of a linkage map, markers were filtered in several steps. First, the vcf file was filtered with `bcftools` (Narasimhan et al. 2016) to only include bi-allelic SNPs, without low quality tags and with a minor allele frequency (MAF) > 0.25 in the parents. All SNPs outside the extended probe regions (120 bp ± 100 bp) and SNPs having a genotype depth (DP) falling outside the range of 10-100 were removed. Progeny genotypes were then filtered using a custom awk script, retaining only genotype calls matching the possible variants available in the Punnet square based on parental genotypes. Genotypes that did not match this criterion were recoded as missing data. Genotyping information was then extracted from the vcf file and all remaining filtering steps were performed in R (R Core Team 2018).

For the map construction we only used markers where both genotyping methods in the parents (capture probes and WGS) showed concordance, where at least one of the parents was heterozygous and where no more than 20% of the progeny had missing data. A chi-square test for segregation distortion was performed on all remaining markers and all markers with a significance level > 0.005 were kept. Finally, only the best marker, in terms of lowest level of missing data and most balanced segregation pattern was kept for each probe. Genotypes and marker segregation pattern were then recoded to `BatchMap` input format (Schiffthaler et al. 2017). The resulting file contained 764 F_1_ progeny and 19,520 probe markers, segregating either in the mother (cross type D1.10), the father (cross type D2.15) or both parents (cross type B3.7).

Framework linkage maps were created using `BatchMap` (Schiffthaler et al. 2017), a parallelized implementation of `OneMap` (Margarido et al. 2007), using the pseudo test cross strategy. Pairwise estimates of recombination frequency were calculated between all markers using a LOD score of 8 and a maximum recombination fraction (`max. rf`) of 0.35. To reduce the number of redundant markers in the map, identical markers (showing no recombination events between them) were collected into bins where one representative marker (the marker with lowest amount of missing data) per bin was used in subsequent analyses. Following binning, markers were grouped into linkage groups (LGs) using a LOD threshold of 12 and further split into a maternal (D1.10 and B3.7 markers) and paternal (D2.15 and B3.7 markers) mapping population with B3.7 markers acting as a bridge between the two parental maps. Marker ordering along the LGs was estimated using 16 rounds of the `RECORD` (Recombination count and ordering) algorithm (Van Os et al. 2005) parallelized over 16 cores. Genetic distance estimates were calculated using three rounds of the ‘map batches’ approach (Schiffthaler et al. 2017) using the Kosambi mapping function. In successive rounds, markers were rippled in sliding windows of eleven, nine and seven markers, respectively, using 32 ripple cores and 2 phasing cores (Margarido et al. 2007, Schiffthaler et al. 2017).

A consensus map of the two parental framework maps was created with the R-package `LPmerge` (Endelmann et al. 2014) using a maximal interval setting ranging from one to ten and equal weight to the two parental maps. The consensus map with the lowest mean root mean square error (RMSE) was set as the best consensus map for each LG.

In order to estimate the correspondence between different LGs from the linkage map and chromosomes in *P. trichocarpa*, probe sequences were mapped against the masked *P. trichocarpa* genome assembly v3.0 using `BLASTn` (Altschul et al. 1990). For each probe with a corresponding marker in the consensus map, the BLAST hit with the highest bitscore value was considered to be the homologous region of the *P. trichocarpa* genome. The number of homologous regions observed between the *P. tremula* LGs and the *P. trichocarpa* chromosomes were used to assign *P. tremula* LGs to corresponding *P. trichocarpa* chromosomes (Figure S1).

### Physical assembly

We used the Python software `AllMaps` (Tang et al. 2015) to create physical chromosomes from the *P. tremula* genome assembly v.1.1 based on the framework genetic maps. Briefly, `AllMaps` uses information from linkage maps to physically anchor scaffolds from a genome assembly into chromosomes. All markers that had been placed into bins at the beginning of the linkage map creation were reintroduced to the final parental framework maps by placing them at the same chromosome and genetic distance as the bin representative marker. All scaffolds in the framework maps that had markers mapped to more than one chromosome (340 scaffolds), or where markers were mapped to different positions on a single LG but more than 20 centiMorgans (cM) apart (19 scaffolds), were split and placed using the corresponding positions of the markers (Figures S2, S3 and S4). To achieve as accurate scaffold splits as possible, assembly gaps and gene annotations were considered. For each scaffold region anchored in the framework maps, the largest assembly gap outside gene models were chosen to split scaffolds. If no assembly gaps were present within the split region, the scaffolds were split in the middle of an intergenic region by artificially creating a gap of size 1 bp. However, if the split region was positioned within a single gene model, the gene models were split either at the largest assembly gap or by artificially creating a gap of size 1 bp in the middle of the region (Figure S5). The python software `jcvi` (Tang et al. 2015) was used to physically split and rename multi-mapped scaffolds. One scaffold, *Potra001073,* contained two markers that could not be split using the rules above since they were positioned in two overlapping gene models appearing on the same strand. These markers were therefore removed from the map. After splitting scaffolds, the linkage map marker positions were translated to the new split scaffold assembly positions using UCSC `liftOver` (Hinrichs et al. 2006) and used as input to `AllMaps`.

`AllMaps` was run according to instructions (https://github.com/tanghaibao/jcvi/wiki/ALLMAPS) using the parental framework maps, here after referred to as the ‘female’ and ‘male’ map, respectively. The maps were merged into the input bed file and weighed equally (1) for scaffold ordering. After ordering, the built-in gap length estimation in `AllMaps` was run to produce more precise lengths for the chromosomes. The chromosome-scale assembly produced will be referred to as *P. tremula* v1.2.

### Linkage map-based recombination map

The parental framework maps as well as the consensus map were edited with custom awk scripts to match the input format specified in the manual of the R-package MareyMap (Rezvoy et al. 2007). All genetic maps were converted to bed format using a custom awk script for easy lift over to the new physical assembly with the UCSC `liftOver` tool (Hinrichs et al. 2006). The lift-over was performed with the ‘—`bedPlus`’ option enabled to carry over extra columns and then recoded back to `MareyMap` input format. Some of the male chromosomes were reversed relative to the female and consensus maps (see negative *ρ*-values in Figure S6). This was done by taking the absolute values of the genetic distance column after subtracting the maximum value of genetic distance from all the values in the column using a custom-made Python script. The edited maps were read into `MareyMap` and two obvious outliers caused by a splitting oversight on chr5 (Figures S2 and S6) and an artificial gap caused by `LPmerge` when creating the consensus map from the parental framework maps in chr16 (Figures S2 and S6) were removed. Finally, we used the ‘sliding window’ method in `MareyMap` to estimate recombination rate in windows of 1Mbp, with a step size of 250 kbp and a minimum number of SNP’s per window of 8, in order to avoid regions with large gaps being assigned recombination values.

### LD-based recombination map

To estimate LD-based recombination rates, the vcf-file with data for the 94 SwAsp individuals from Wang et al. (2018) was lifted over to v1.2 genome coordinates by first recoding the file to bed format using `vcf2bed` from the `BEDOPS` toolkit (Neph et al. 2012), lifted over with UCSC `liftOver` (Hinrichs et al. 2006) and finally recoded back to vcf format. The resulting vcf file was filtered using `vcftools` (Danecek et al. 2011), retaining only bi-allelic SNPs with a minor allele frequency greater than 0.05 (maf > 0.05) and that showed no evidence for deviations from Hardy-Weinberg equilibrium (p > 0.002).

We used `LDhelmet` v.1.10 (Chan et al. 2012) to produce a LD-based recombination map. `LDhelmet` handles a maximum of 25 diploid individuals (ie. 50 haplotypes), and we therefore sampled a random subset of 25 individuals from the 94 re-sequenced SwAsp individuals utilizing the `vcftools` ‘`--max-indv`’-option. The subset vcf was split into separate files according to chromosomes to avoid memory issues when running `LDhelmet`. Full FASTA-files were produced for the 25 individuals by reassigning SNP-positions at the corresponding sites within the reference FASTA for *P. tremula* v1.2 using the ‘`vcf-consensus`’ script from `vcftools`. This was done twice per individual to produce both copies of the chromosome. Individual files were then concatenated together to generate a single FASTA file per chromosome with data for all 25 individuals.

The `LDhelmet` preparatory files were produced as suggested in the `LDhelmet` v1.10 manual (Chan et al. 2012). We produced Padé coefficients and lookup tables separately for each of the 19 chromosomes. The configuration files were created using a window size of 50 and the lookup table was produced using the recommended values found in the `LDhelmet` manual (-t 0.01 -r 0.0 0.1 10.0 1.0 100.0). Similarly, the Padé coefficients tables were produced using the recommended values from the manual (-t 0.01-x 11). To further lighten the computational load during the recombination estimation step, we opted to use the SNPs and pos files for our chromosomes. These files were produced for each chromosome separately using the ‘`–ldhelmet`’-option for `vcftools` v0.1.15. We used the recommended values from the `LDhelmet` manual as *Populus tremula* has similar levels of nucleotide polymorphism and extent of linkage disequilibrium as *Drosophila melanogaster,* on which the recommended settings in `LDhelmet` are based upon (Wang et al. 2016)

`LDhelmet` outputs estimates of recombination in units of ρ/bp whereas the genetic map is in units of cM. We therefore converted the `LDhelmet` results to cM distances following the method outlined in Booker et al. (2017) to be able to make comparisons between the recombination rates estimated using the two methods. The conversion assumes that since the physical size of a chromosome is constant for the two methods, the cumulative genetic distance in either cM or ρ should be the same but on different scales. The cumulative ρ was calculated by multiplying the ρ/bp estimates with the distance in bp between the adjacent SNP’s and then summed across chromosomes. Knowing the cumulative ρ and corresponding cM-values, it is possible to derive a ‘scaling factor’ to calculate cM values from the corresponding ρ values. The resulting cM values were read into `MareyMap` and recombination rates were estimated as described earlier for the genetic map-based recombination map.

### Correlation of recombination rate estimates, genetic correlates of recombination rate and model of recombination rate

We compared recombination rates inferred from the consensus genetic map or from the sequence data by calculating correlations across 1 Mb windows. We also assessed correlations between the two recombination rates and a number of genomic features, including gene density, repeat density, GC-content, substitution density, neutral diversity and methylation.

Gene and repeat density were estimated using `bedtools` (Quinlan 2014) ‘`makewindows`’ option to split the chromosomes into 50 kbp chunks and then using the ‘`annotate`’ option to calculate density of the repeat and gene elements respectively using gff files containing locations of these elements (available at ftp://plantgenie.org/Data/PopGenIE/Populus_tremula/v1.1/gff3/). GC-content was calculated from the FASTA file produced from AllMaps using an awk script (modified from: https://www.biostars.org/p/70167/#70172). The original script was modified to have window functionality across a FASTA sequence and to take into consideration sequence gaps. Windows with more than 80% gaps (N) were discarded to avoid biased results.

Substitutions relative to *P. trichocarpa* were estimated from a vcf-file with SNP-calls for a single *Populus trichocarpa* individual mapped against the *Populus tremula* reference genome v1.1. Comparisons of putative substitutions were then made against a list of SNP positions from the 94 SwAsp *P. tremula* data set using the vcftools option ‘`--exclude-positions`’. Substitutions relative to *P. tremuloides* were inferred in a similar way using a vcf-file containing SNP calls from five *P. tremuloides* individuals mapped against the *P. tremula* reference genome v1.2 (Lin et al. 2018). Both the options ‘`--exclude-positions`’ and ‘`–positions`’ were used to obtain the substitutions fixed in both aspen species relative to *P. trichocarpa* (hereafter referred to as ‘old’ substitutions) and substitutions fixed in *P. tremula* only (‘new’ substitutions) respectively. Files containing old, new and the total number of substitutions were used to calculate substitution densities in windows of 50 kbp using the ‘`--SNPdensity`’ option in `vcftools`. Finally, neutral genetic diversity was estimated using `vcftools` with the ‘`--windowed-pi`’ option in 50 kbp windows using only 4-fold degenerate sites or intergenic sites located at least 2 kbp from an annotated gene. For all these calculations data for all 94 SwAsp samples were used.

Methylation levels were estimated using bisulfite sequencing data from six SwAsp individuals. The six individuals were bisulfite sequenced using two biological replicates per individual using paired-end (2×150) sequencing on an Illumina HiSeq X at the National Genomics Infrastructure facility at Science for Life Laboratory in Uppsala, Sweden. Based on recommendation from the NGI facility, samples were sequenced to an average depth of 60x as approximately 50% of the bisulfite sequencing data cannot be mapped uniquely due to excessive damage induced by the bisulfate treatment. The raw bisulfite-sequencing reads were trimmed using `trimGalore` v. 0.4.4 (https://github.com/FelixKrueger/TrimGalore), a wrapper around `Cutadapt` (Martin 2011) and `FastQC` (Andrews 2010), with a paired-end trimming mode and otherwise default settings. In order to obtain as accurate methylation calls as possible, polymorphic substituted versions of the *P. tremula* v1.1 assembly (Lin et al. 2018) were created for each sample separately using `bcftools` consensus (Narasimhan et al. 2016) and bisulfite converted and indexed with `bismark_genome_preparation` (Krueger and Andrews 2011). Trimmed reads for all individuals were mapped against the corresponding converted reference genomes using `Bismark` (Krueger and Andrews 2011) and `Bowtie2` (Langmead and Salzberg 2012). In order to remove optical duplicates from the BAM files, we ran `deduplicate_bismark` with default settings before methylation levels were extracted using `bismark_methylation_extractor` with the following settings: `--bedGraph --gzip --comprehensive --CX --scaffolds --buffer size 20G`, which produced both context files and coverage files. For one of the sequenced individuals (SwAsp046), subsequent analyses suggested that the two biological replicates were actually derived from two genetically distinct individuals, likely due to a sampling mix-up in the common garden. For all individuals, the biological replicate with largest amount of data available was used in downstream analyses. The `Bismark` results were lifted over to v1.2 coordinates using UCSC `liftOver` with the ‘ `--bedPlus`’ option. Coverage-files were filtered for low (< 5) and high (> 44) coverage observations to remove spurious results due to low coverage or collapsed duplicate genomic regions, respectively. Following filtering, coverage files for all six samples were merged and the different contexts of methylation (GpG, CHG and CHH) were extracted from the `Bismark` context files.

To produce data sets for all genomic features that were comparable to the two recombination maps, average values were calculated across 1 Mbp window using a step size of 250 kbp using a custom-made Python script. Correlations between the two recombination maps and between the recombination maps and genomic features were calculated using R. We also assessed the independent effects of different variables on the two recombination rate estimates using multiple regression.

## Results

### *P. tremula* linkage maps

14,598 unique markers in the female map and 13,997 unique markers in the male map were distributed across 3,861 and 3,710 scaffolds, respectively. Markers in both parental framework maps grouped into 19 linkage groups (LGs), corresponding to the haploid number of chromosomes in *Populus* (Table 1). Among the mapped scaffolds, 19 scaffolds contained markers that mapped to different positions within the same LG, but that were more than 20 cM apart (Figure S3) and 340 scaffolds contained markers that mapped to two or more LGs (Figures S2 and S4). These scaffolds were split according to criteria described in materials and methods. Additionally, there were 49 scaffolds split within gene models (Figure S5). Two ambiguous markers in scaffold Potra001073 were removed. After splitting 14,596 (12,900 binned) and 13,996 (12,382 binned) markers from 4,184 and 4,011 scaffolds remained. These markers spanned 4072.72 cM and 4053.68 cM for the female and male framework maps, respectively. The parental maps were used to produce a consensus genetic map consisting of 19,519 markers derived from 4,761 scaffolds spanning 4059.00 cM (Table 1). Linkage groups (LG) were assigned to the corresponding *P. trichocarpa* homologs through synteny assessment (Table 1, Figure S1).

**Table 1.** Summary of female male and consensus linkage maps for each chromosome.

| Chr | LG | Female |  |  | Male |  |  | Consensus |  |
| --- | --- | --- | --- | --- | --- | --- | --- | --- | --- |
|  |  | Probe markers | Bin markers | Size (cM) | Probe markers | Bin markers | Size (cM) | Probe markers | Size (cM) |
| 1 | 6 | 1,784 | 1,573 | 498.0 | 1,737 | 1,542 | 499.3 | 2,417 | 498.8 |
| 2 | 3 | 1,054 | 924 | 266.3 | 1,028 | 904 | 268.3 | 1,407 | 266.3 |
| 3 | 8 | 826 | 736 | 229.9 | 790 | 695 | 235.3 | 1,117 | 235.2 |
| 4 | 1 | 813 | 728 | 217.2 | 835 | 736 | 263.0 | 1,128 | 220.5 |
| 5 | 7 | 965 | 842 | 284.5 | 947 | 836 | 270.6 | 1,293 | 271.9 |
| 6 | 14 | 1,241 | 1,108 | 311.1 | 1,116 | 993 | 294.3 | 1,602 | 296.4 |
| 7 | 18 | 572 | 497 | 161.9 | 501 | 450 | 155.1 | 765 | 155.5 |
| 8 | 12 | 895 | 781 | 227.4 | 824 | 726 | 211.0 | 117 | 215.5 |
| 9 | 2 | 697 | 617 | 171.5 | 640 | 578 | 171.6 | 879 | 172.2 |
| 10 | 4 | 1,026 | 901 | 279.0 | 1,015 | 879 | 270.8 | 1,388 | 273.7 |
| 11 | 16 | 515 | 460 | 181.4 | 541 | 480 | 180.7 | 724 | 182.0 |
| 12 | 17 | 515 | 452 | 142.4 | 486 | 429 | 140.7 | 678 | 143.3 |
| 13 | 11 | 586 | 530 | 173.7 | 608 | 531 | 176.4 | 822 | 178.7 |
| 14 | 5 | 733 | 655 | 194.3 | 702 | 609 | 182.0 | 983 | 194.3 |
| 15 | 10 | 553 | 505 | 145.8 | 593 | 532 | 157.9 | 777 | 159.2 |
| 16 | 19 | 383 | 334 | 145.8 | 220 | 195 | 125.8 | 430 | 145.4 |
| 17 | 15 | 488 | 433 | 155.5 | 447 | 394 | 148.9 | 647 | 149.4 |
| 18 | 13 | 591 | 502 | 156.3 | 592 | 531 | 166.2 | 792 | 165.5 |
| 19 | 9 | 361 | 322 | 130.6 | 375 | 342 | 135.8 | 500 | 135.8 |
| Total |  | 14,598 | 12,900 | 4072.7 | 13,997 | 12,382 | 4053.7 | 19,519 | 4059.0 |

### Physical assembly Potra v1.2

The parental framework maps were used to produce a physical map, Potra v1.2, of the *P. tremula* chromosomes that we used to estimate recombination maps (Figure 1, Figure S6). The scaffolds anchored from the parental framework maps spanned 210.7 Mbp and 205.1 Mbp for the female and male respectively. This corresponds to 54.6% and 53.1% of the 385.8 Mbp covered by the v1.1 assembly (Table S1). Of the 4,761 scaffolds with markers, 96.6% could be anchored in the assembly providing a total physical assembly consisting of 223.4 Mbp. This corresponds to 57.9% of the v1.1 *P. tremula* assembly (Lin et al. 2018) (Table 2). 75.7% of the physical map was both anchored and oriented, while the remaining 24.3% was only anchored. 199,967 of the v1.1 assembly scaffolds were not covered by the framework maps and thus could not be anchored to the physical map. There was a clear distinction between the scaffolds we could and could not anchor to the Potra v1.2 assembly. The median length for the 4214 scaffolds anchored in the map was 37 kb and these scaffolds contain 26,808 predicted gene models. Conversely, the median length on unanchored scaffolds was only 0.3 kb and they collectively contain only 8,501 predicted gene models (Table 2). After the initial assembly, gap estimation added 43 Mbp of gap sequences across the genome, increasing the estimated total size of the v1.2 assembly to 265 Mbp. This is approximately 55% of the 479 Mbp genome size estimated for *P. tremula* (Lin et al. 2018).

**Figure 1.**
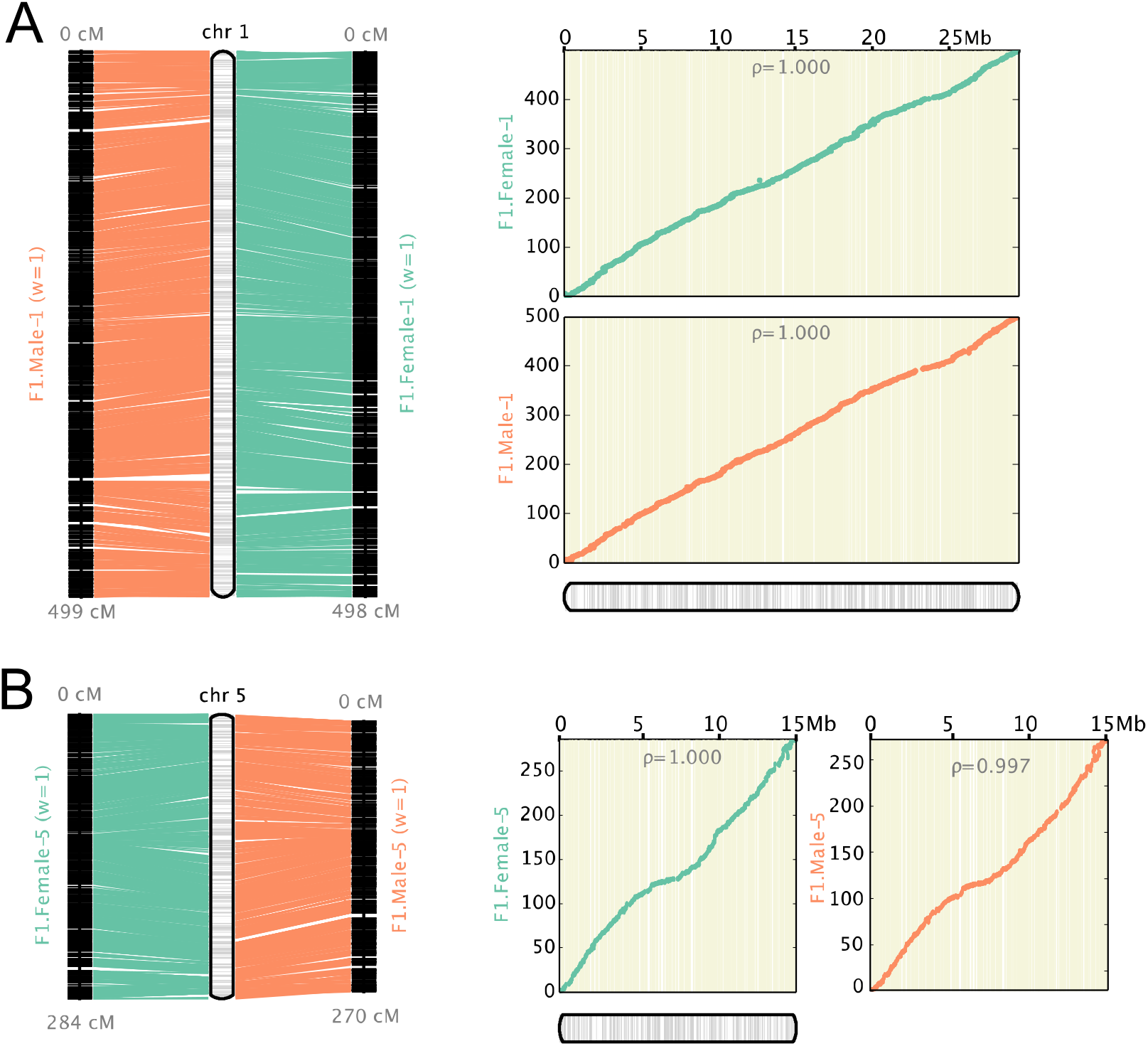
Genetic maps and resulting physical map created by Allmaps for Chr1 and Chr5. The left panel shows the marker distribution (in cM) for the genetic maps and the anchored genomic region (in Mb) for the physical map, while the right panel is showing the Marey maps, i.e. the correspondence between the physical (x-axis) and recombination-based (y-axis) position of markers. The female map is depicted in green and the male map is depicted in orange.

**Table 2.** Summary of physical assembly Potra vl.2.

|  | Anchored | Oriented | Unplaced |
| --- | --- | --- | --- |
| Markers (unique) | 19,302 | 15,597 | 215 |
| Markers per Mb | 86.4 | 92.2 | 1.3 |
| Gene elements | 26,808 (75.9%) | NA | 8501 (24.1%) |
| N50 Scaffolds (bp) | 2,071 | 1,589 | 238 |
| Scaffolds | 4,599 | 2,759 | 200,129 |
| Scaffolds with 1 marker | 1,151 | 0 | 133 |
| Scaffolds with 2 markers | 999 | 787 | 17 |
| Scaffolds with 3 markers | 620 | 384 | 7 |
| Scaffolds with >=4 markers | 1,829 | 1,588 | 5 |
| Total bases | 223,381,844 (57.9%) | 169,187,619 (43.9%) | 162,436,408 (42.1%) |

### Recombination estimates

Recombination estimates were produced based on the consensus linkage map (LMB) and from LD data (LDB) derived from 25 randomly selected individuals from the SwAsp collection. The LMB recombination rate estimates varied between 1.605 cM/Mbp on chromosome 4 to 26.911 cM/Mbp on chromosome 11, while the LDB estimates varied between 1.969 cM/Mbp on chromosome 5 to 231.801 cM/Mbp on chromosome 1. The median estimated recombination rate in the LMB map was 16.0 cM/Mbp with a mean of 15.6 cM/Mbp, whereas the median recombination rate for the LDB map was 14.0 cM/Mbp with a mean of 16.1 cM/Mbp (Table S2, Figure S7). The majority of all recombination rate estimates (97%) for both maps fell in the range of 2 – 27 cM/Mb (Figure 2, Figure S8). There were 20 windows where the LDB estimates are 1.5-15-fold higher compared to the corresponding rates from the LMB estimates and 2-14-fold higher than the mean recombination rate estimate from the LDB map. 13 of these windows had recombination rates exceeding 27 cM/Mbp, while seven were within 2-27 cM/Mbp. Conversely, there were also 23 windows where the LDB estimates were 2-4 times lower than the corresponding LMB estimates (Figure 2, Figures S8 and S9).

**Figure 2.**
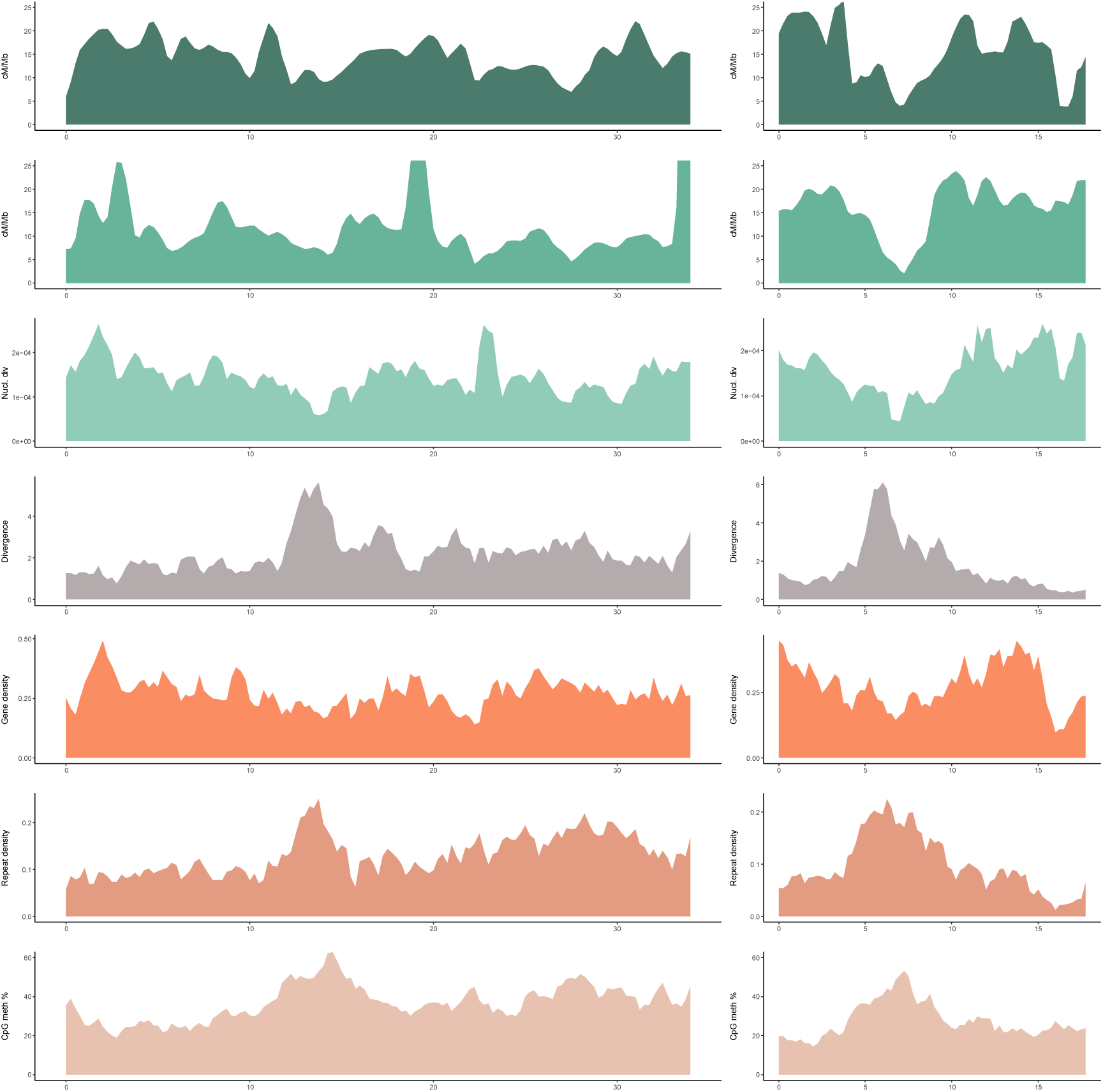
Recombination rates and genomic features calculated in 1Mb windows across chromosome 1 and 5 with a step size of 250kb. A) Recombination rate estimated from the linkage map (cM/Mb) B) Recombination rate estimated from sequence LD data (cM/Mb) C) Nucleotide diversity (1/bp) D) Divergence (sites/Mb) E) Gene density (percentage coding/Mb) F) Repeat density (percentage repeats/Mb) G) CpG methylation (percentage/Mb)

### Correlation of recombination rate estimates, genetic correlates of recombination rate and model of recombination rate

The correlation between recombination rate estimates for the LMB and LDB maps was 0.478 (Spearman’s rank correlation) (Figure 3). This was the strongest positive correlation of all of the correlations calculated for both maps and strongest correlation overall for the LDB map. Correlations between the LMB and LDB recombination maps and neutral diversity were the second strongest positive correlations for both maps, 0.447 and 0.442 respectively. This correlation was the second strongest overall for LDB map. Correlation with neutral diversity was also the only variable where there was no notable decrease in the correlation coefficient from the LMB to the LDB recombination maps (Figure 3).

**Figure 3.**
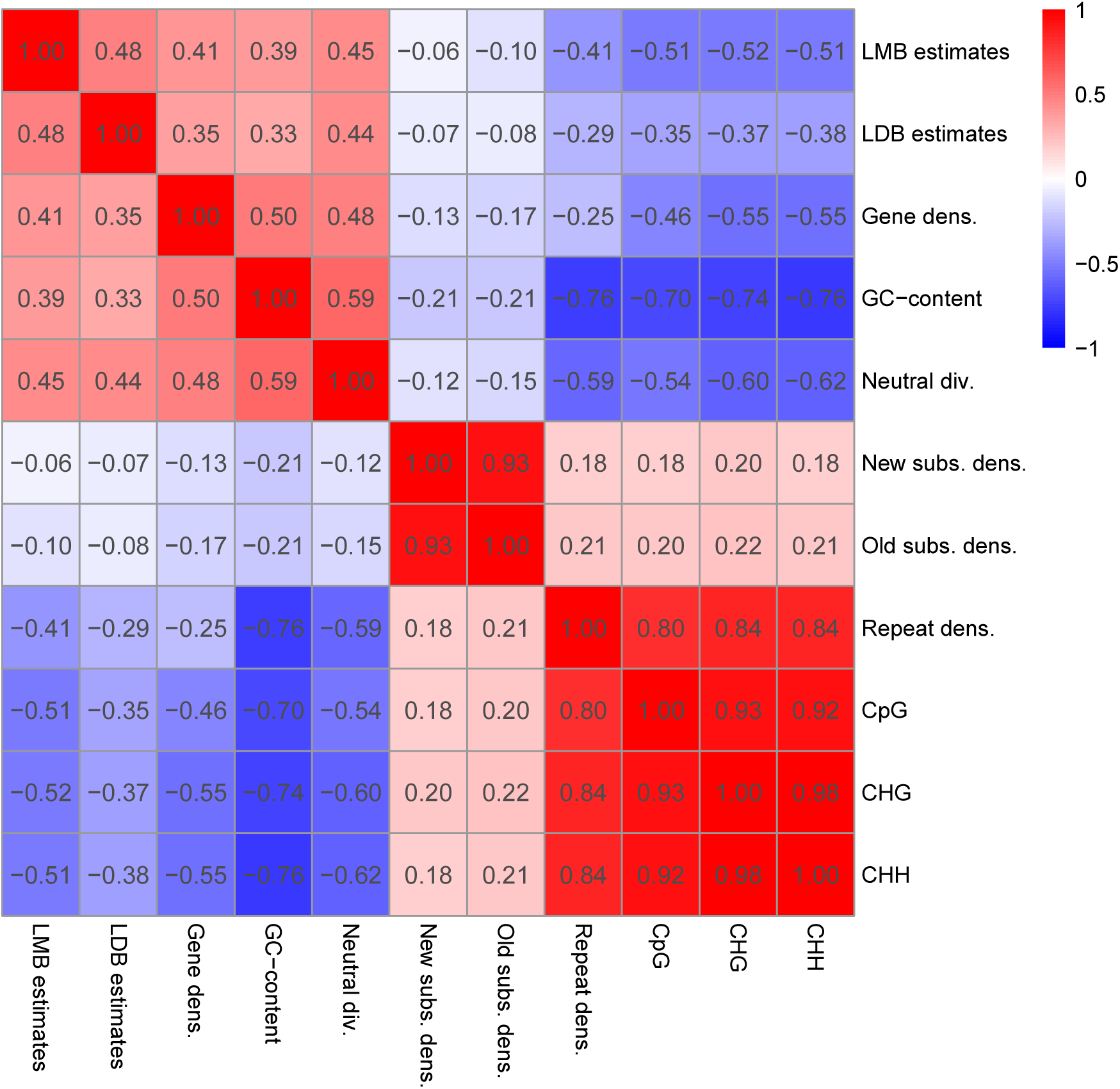
Correlations between recombination rates and genomic features.

For the LMB map we observed strong negative correlations with CHG methylation (−0.515), CHH methylation (−0.511) and CpG methylation (−0.505). For the LDB map the corresponding correlations were −0.379 (CHH), −0.371 (CHG) and −0.353 (CpG). Methylation levels were also strongly correlated with each other (0.918-0.984) and with repeat density (0.800-0.840). Repeat density was also moderately negatively correlated with recombination rate estimates from both the LMB (−0.408) and LDB maps (−0.291). Both recombination maps showed only weak correlations (−0.1< *ρ* <0.1) with either old or new neutral substitution densities (−0.06 --0.1). Neutral substitution densities and neutral diversity showed only weak negative correlations (Figure 3). Overall, the LMB estimates displayed consistently stronger correlations with the different genomic features compared to the LDB estimates. This is in line with 5 % of the total variation being explained by a multiple regression model for the LDB recombination estimates compared to 35 % variation explained for the LMB estimates (Table 3).

**Table 3.**
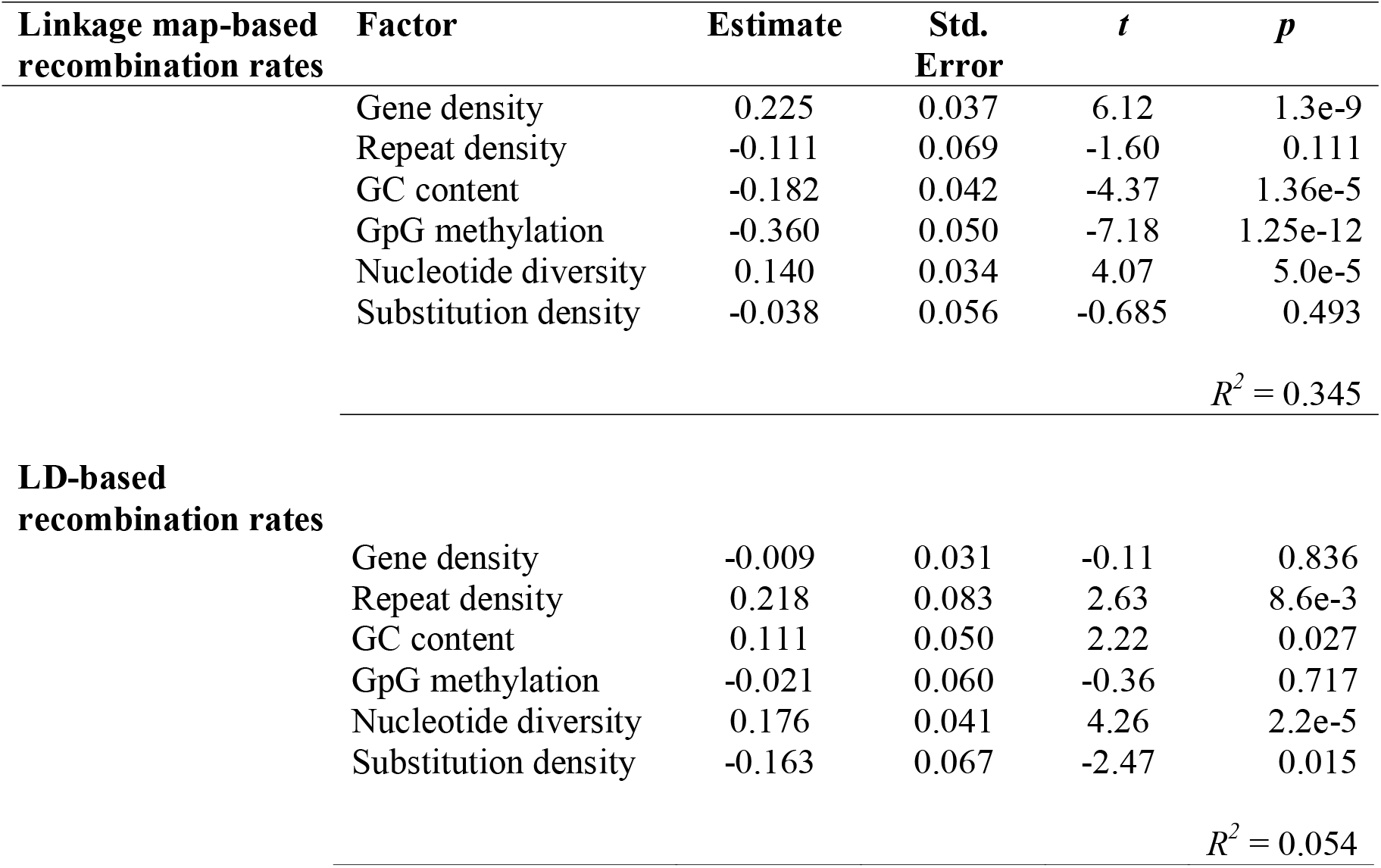
Multiple regression of recombination rate and various genomic

## Discussion

### *P. tremula* fine-scale genetic maps and physical assembly Potra v1.2

The genetic maps presented here are the most marker-dense maps produced for *P. tremula* to date. Our female map is only 20 cM larger than the male map, despite having 518 more informative markers (Table 1) and most chromosomes have size differences of less than 10 cM between sexes (Table 1, Figure S6). However, in cases where we observed differences between the maps for the two sexes exceeding 10 cM, the male map is shorter in all but one case. These results, together with the overall shorter linkage map for the male, could suggest overall lower recombination rates in males, in line with what has been observed in many other highly outcrossing plant species (Lenormand and Dutheil 2005). The high marker density in our framework genetic maps allow us to anchor 57.9 % of the *P. tremula* v1.1 genome assembly on to the expected 19 chromosomes, providing us with the first chromosome-scale assembly for *P. tremula* (Table 2).

The map length in the section *Populus,* to which *P. tremula* belongs (Wang et al. 2014), has previously been estimated to be 1,600-3,500 cM (e.g. Zhang et al. 2004, Paolucci et al. 2010, Zhigunov et al. 2017). The most relevant comparison for our purposes is the recently produced linkage maps in *P. tremula* by Zhigunov et al. (2017). Their map contains 2000 informative markers with an average marker distance of 1.5 cM that were observed in 122 progenies, resulting in a total map length of 3000-3100 cM. Our framework maps are much denser with ca. 12000-13000 informative markers (Table 1) and with an average distance of ~0.3 cM between markers. In addition, our map is based on a mapping population consisting of 764 progenies and we are hence able to achieve a far greater resolution in our maps. However, larger data sets, both with respect to the number of markers and the number of progenies used, increase the risk of genotyping errors. Genotyping errors will ultimately lead to an inflation of map sizes as errors can be interpreted as recombination events during map creation and this could help explain why our maps are roughly 1000 cM longer than those reported by Zhigunov et al. (2017), given that we use 5-7-fold more markers and a mapping population that is six times larger.

On the other hand, our framework genetic maps are similar in size to the ca. 4200 cM and 3800 cM maps presented by Tong et al. (2016) for the more distantly related species *Populus deltoides* and *Populus simonii,* respectively. The large size of these maps led Tong et al. (2016) to suggest that their maps were suffering from inflation due to the difficulty of properly ordering a large number of markers within a linkage group. While we likely also suffer from such size inflation, these issues appear to be less severe in our *P. tremula* parental maps, which contain between 8-14 times the number of markers used in the *P. deltoides* (1601) and *P. simonii* (940) maps and yet yield linkage maps of similar size. One explanation for this is our considerably larger mapping population compared to the *P. deltoides* and *P. simonii* maps (299 progenies). A greater number of segregating progenies helps mitigate the problems of ordering a larger number of markers by increasing resolution of recombination detection.

### Recombination rate estimates

Recombination rates estimated from both the consensus linkage map and from polymorphism data showed substantial variation across all chromosomes on Mbp scales (Figure 2). For the consensus genetic map-based estimates and LD-based estimates, the majority of our observations fell in the range 0-27 cM/Mb (Figure 2, Figure S8) which is similar to what has been observed in other plants such as *Arabidopsis thaliana* (Giraut et al. 2011), *Populus trichocarpa* (Slavov et al. 2012) and *Eucalyptus grandis* (Silva-Junior and Grattapaglia 2015), where recombination rates across chromosomes mostly fall within 0-25 cM/Mb.

We observed a small number of genomic windows where the LDB recombination rates were either 1.5-15-fold higher or lower than the corresponding estimates based on the consensus genetic map (Figure 2, Figure S8). While we do not know for certain what causes these large differences in recombination rates, a possible explanation could be that such windows harbor recombination hotspots or coldspots that the comparatively coarse linkage map fails to detect. Recombination hotspots, with local recombination rates 10 to 100-fold higher than the genome-wide average, have been observed in a number of species, including *Drosophila melanogaster* (Chan et al. 2012), *Arabidopsis thaliana* (Kim et al. 2007), *Zea mays* (He and Dooner 2009), *Oryza sativa* (Si et al. 2015) and *Eucalyptus grandis* (Silva-Junior and Grattapaglia 2015). Similarly, coldspots have been identified in *Zea mays* (He and Dooner 2009) and *Oryza sativa* (Si et al. 2015) among others. Hotspots or coldspots for recombination are, however, often quite restricted in size (Choi and Henderson 2015), spanning only a few kb, and the relatively coarse recombination maps produced here are consequently not suitable for accurate detection of such regions.

The average recombination rate in *P. tremula* is 2-27 times higher than those found in a number of, mostly domesticated, plant and animal species (reviewed in Henderson (2012) and Tiley and Burleigh (2015)), suggesting that *P. tremula* exhibits recombination rates that are among the highest recorded in the animal and plant kingdoms. Of the species covered in these reviews, *P. trichocarpa* (Slavov et al. 2012) makes for the most interesting comparison since it is one of the few undomesticated species listed, is closely related to *P. tremula* and has previously been compared with *P. tremula* (Wang et al. 2016). Despite the close relationship, the average recombination rate in *P. trichocarpa* (Slavov et al. 2012) is less than a third of what we estimated for *P. tremula.* Similar observations were previously made by Wang et al. (2016) who found that population-based recombination rates in *P. trichocarpa* were on average only a quarter of the corresponding values in *P. tremula.* Wang et al. (2016) argued that the differences they observed in recombination rates between *P. tremula* and *P. trichocarpa* could at least partly stem from differences in the effective population size (N_e_) of two species (Wang et al. 2016). In light of this, it would be interesting to perform further comparisons of recombination rates in *P. tremula* with other *Populus* species that have wide distribution ranges and large Ne, such as *P. deltoides* (Tong et al. 2016) or *P. tremuloides* (Wang et al. 2016).

### Correlations between recombination rate and genomic features

Recombination rate estimates from the consensus linkage map and from polymorphism data showed a moderately strong positive correlation (>0.4) (Figure 3). A similar correlation between linkage map and LD-based estimates of recombination was also been observed in house mouse by Booker et al. (2017), suggesting that LDB recombination rate estimates are reliable substitutes for genetic map-based recombination rate estimates.

We observed a strong positive correlation between recombination rate and gene density (0.45 and 0.41 respectively) (Figure 3). This is in line with earlier observations in plants (Tiley and Burleigh 2015, Stapley et al. 2017) and implies that recombination may be linked to gene-dense regions through a higher recruitment of the recombination machinery to euchromatic genome regions. Preferential recruitment of recombination to euchromatic genome regions has also been put forward as an explanation for why recombination rates across plants generally show stronger correlations with gene density compared to genome size (Henderson 2012, Tiley and Burleigh 2015). Studies in plants like *Arabidopsis thaliana* and *Oryza sativa* have shown that while crossover events are enriched in genic regions, they mostly occur in promoters a few hundred bps upstream of the transcription start site or downstream of the transcription termination site (Choi et al. 2013, Marand et al. 2019).

We observed negative correlations between local recombination rates and both repeat density and methylation (Figure 3), in line with earlier results that highlighted the role of chromatin features in establishing crossover locations in plants (Choi et al. 2013, Marand et al. 2019). For instance, Choi et al. (2013) showed that methylation is lower at observed sites of crossovers and Rodgers-Melnick et al. (2015) showed that cross-over density in *Zea mays* is negatively correlated with repeats and CpG methylation. All methylation contexts were highly correlated in our data and also strongly correlated with repeat density (≥0.8, Figure 3), in line with the observation that most repetitive elements in plant genomes are strongly methylated (Saze and Kakutani 2011).

Compared to earlier results from *P. tremula,* we observed a weaker correlation between recombination rates and gene density (Wang et al. 2016). One possible reason for this is likely to be the reference genome used. Wang et al. (2016) based their analyses on the *P. trichocarpa* reference genome whereas our analyses were based on a *de novo* assembly for *P. tremula.* The *P. trichocarpa* assembly, while more contiguous than our current *P. tremula* assembly, is less ideal for these types of analyses since divergence between the two species leads to substantially reduced rates of read mapping primarily in intergenic regions (Lin et al. 2018). Our current assembly, while only representing 55 % of the expected genome size *of P. tremula,* likely offers a more unbiased set of genomic regions where we are able to call genetic variants. In contrast, the data derived from using the *P. trichocarpa* reference genome likely suffers from under-representations of repeat-rich regions and other intergenic regions (Wang et al. 2016).

GC-content was positively correlated with both our recombination rate estimates, similar to what has been observed in humans *(Homo sapiens)* (e.g. Fullerton et al. 2001) and *Arabidopsis thaliana* (Kim et al. 2007) among others. However, when GC-content was included in a multiple regression model with other genomic features, the direct effect of GC content was actually negative for the LMB recombination rate (Table 3). GC-content is strongly correlated with gene density (0.50) in *P. tremula,* and gene density is in turn also strongly positively correlated with recombination (Figure 3). The strand separation needed in the strand invasion of meiotic recombination is harder to achieve in areas with high GC-content due to higher annealing energy and can explain why GC-content has a direct negative effect on recombination rates when effects of gene density are accounted for (Table 3, e.g. Mandel and Marmur 1968).

### Effects of linked selection on patterns of nucleotide diversity in *P. tremula*

Both of our recombination rate estimates were strongly correlated with nucleotide diversity at putatively neutral sites (Figure 3). A positive correlation between local recombination rate and nucleotide polymorphism is usually interpreted as a signature of ubiquitous natural selection acting either through positive (hitchhiking) or negative (background) selection (Begun and Aquadro 1992). Alternatively, such a correlation could also arise if recombination itself is mutagenic (Begun and Aquadro 1992). However, if recombination is mutagenic one also expects to see a correlation between recombination and sequence and divergence at neutral sites (Begun and Aquadro 1992). Our data show little evidence supporting the idea that recombination has a direct mutagenic effect as we observed only a weak and negative correlation between local recombination rates and substitutions at putatively neutral sites (Figure 3). In light of this, and in line with earlier results, we observed that linked selection has pervasive effects on neutral diversity across the *P. tremula* genome (Ingvarsson 2010, Wang et al. 2016).

## Conclusions

Our high-density *Populus tremula* genetic maps and the new chromosome-scale genome assembly we present here provide a valuable resource not only for *P. tremula,* but also for comparative genomics studies within the entire *Populus* genus. We have also presented multiple lines of evidence to support the utility of using LD-based estimates of recombination rates as a proxy for genetic map-based estimates. Estimates of recombination rates derived from the two different approaches were in broad agreement and correlations between the two recombination rate estimates and various genomic features were in broad agreement between the two methods. Booker et al. (2017) and Chan et al. (2012) reported similar results in *Mus musculus* and *Drosophila melanogaster*, and our results suggest that LD-based estimates of recombination are also largely applicable to plants. Our results also suggest that LD-based estimates might be especially useful for identifying fine scale recombination variation and features such as recombination hot-or cold-spots by relying on the relatively high density of SNPs within genomes. Finally, we have further verified and extended the observation that linked selection is an important force shaping genome-wide variation in *P. tremula* by showing that the positive correlation between local recombination rates and nucleotide diversity and neutral sites is robust even when factoring in the effects of other genomic features. Although a positive correlation between recombination and diversity is a hallmark signature of linked selection, the pattern can be established by either positive or negative selection. We have earlier documented evidence for both a reduction in levels of standing variation due to recurrent hitchhiking (Ingvarsson 2010) and a reduction in the efficacy of purifying selection at eliminating weakly deleterious in regions of low recombination (Wang et al. 2016). More work is thus needed to assess the relative importance of positive and negative selection in shaping genome-wide variation in *P. tremula.*

## Acknowledgements

The research has been funded through research grants from the Swedish Research Council (PI), the Knut and Alice Wallenberg Foundation (PI, BS) and ‘Trees and Crops for the Future’ (NRS, PKI and SJ). The authors acknowledge support from the National Genomics Infrastructure in Stockholm funded by Science for Life Laboratory, the Knut and Alice Wallenberg Foundation and the Swedish Research Council, and SNIC/Uppsala Multidisciplinary Center for Advanced Computational Science for assistance with massively parallel sequencing and access to the UPPMAX computational infrastructure through projects b2011141 and SNIC 2017/1-499.

## Literature Cited

Altschul SF, Gish W, Miller W, Myers EW, Lipman DJ. 1990. Basic local alignment search tool. J Mol Biol. 215: 403–410.

Andrews S. 2010. FastQC: a quality control tool for high throughput sequence data. Available online at: http://www.bioinformatics.babraham.ac.uk/projects/fastqc

Van der Auwera GA et al. 2013. From FastQ Data to High-Confidence Variant Calls: The Genome Analysis Toolkit Best Practices Pipeline. Current Protocols in Bioinformatics 43:1–33.

Begun DJ, Aquadro CF. 1992. Levels of naturally occurring DNA polymorphism correlate with recombination rates in D. melanogaster. Nature 356: 519–520.

Booker TR, Ness RW, Keightley PD. 2017. The Recombination Landscape in Wild House Mice Inferred Using Population Genomic Data. Genetics 207: 297–309.

Chan AH, Jenkins PA, Song YS. 2012. Genome-wide fine-scale recombination rate variation in Drosophila melanogaster. PLoS Genet 8: e1003090.

Choi K, Henderson IR. 2015. Meiotic recombination hotspots – a comparative view. Plant j 83: 52–61.

Choi K et al. 2013. Arabidopsis meiotic crossover hot spots overlap with H2A.Z nucleosomes at gene promoters. Nat Genet 45: 1327–1336.

Danecek P et al. 2011. The variant call format and VCFtools. Bioinformatics 27 (15): 2156–2158.

DePristo MA et al. 2011. A Framework for Variation Discovery and Genotyping Using Next-Generation DNA Sequencing Data. Nat Genet 43: 491–98.

Endelman JB, Plomion C. 2014. LPmerge: An R Package for Merging Genetic Maps by Linear Programming. Bioinformatics 30: 1623–24.

Farmer Jr RE. 1996. “The genecology of Populus.” in (Stettler R, Bradshaw T, Heilman P, Hinckley T. eds.) “Biology of Populus and its implications for management and conservation”, pp. 33–55 NRC Research Press, Ottawa, Canada.

Felsenstein J. 1974. The evolutionary advantage of recombination. Genetics 78: 737–756.

Fracheboud Y et al. 2009. The control of autumn senescence in European aspen. Plant physiol 149:1982–1991.

Fullerton SM, Carvalho AB, Clark AG. 2001. Local Rates of Recombination Are Positively Correlated with GC Content in the Human Genome, Mol Biol Evol 18(6): 1139–1142.

Giraut L et al. 2011. Genome-Wide Crossover Distribution in Arabidopsis thaliana Meiosis Reveals Sex-Specific Patterns along Chromosomes. PLoS Genet 7(11): e1002354.

Haenel Q, Laurentino TG, Roesti M, Berner D. 2018. Meta-analysis of chromosome-scale crossover rate variation in eukaryotes and its significance to evolutionary genomics. Mol Ecol 27: 2477–2497.

He L, Dooner HK. 2009. Haplotype structure strongly affects recombination in a maize genetic interval polymorphic for Helitron and retrotransposon insertions. P Natl Acad Sci 106(21): 8410–8416.

Henderson IR. 2012. Control of meiotic recombination frequency in plant genomes. Curr opin plant biol 15: 556–561.

Hinrichs AS et al. 2006. The UCSC Genome Browser Database update 2006. Nucleic Acids Res 1;34: D590–598.

Ingvarsson PK. 2010. Natural selection on synonymous and nonsynonymous mutations shapes patterns of polymorphism in Populus tremula. Mol biol evol 27: 650–660.

Jansson S, Douglas CJ. 2007. Populus: a model system for plant biology. Annu rev plant biol 58: 435–458.

Kim S et al. 2007. Recombination and linkage disequilibrium in Arabidopsis thaliana. Nat genet 39(9): 1151–1155.

Kouki J, Arnold K, Martikainen P. 2004. Long-term persistence of aspen – a key host for many threatened species – is endangered in old-growth conservation areas in Finland. J Nat Conserv 12: 41–52.

Krueger F, Andrews SR. 2011. Bismark: a flexible aligner and methylation caller for Bisulfite-Seq applications. Bioinformatics 1;27(11): 1571–1572.

Kulathinal RJ, Bennett SM, Fitzpatrick CL, Noor MAF. 2008. Fine-scale mapping of recombination rate in Drosophila refines its correlation to diversity and divergence. P Natl Acad Sci USA 105: 10051–10056.

Langmead B, Salzberg S. 2012. Fast gapped-read alignment with Bowtie 2. Nat Methods 9: 357–359.

Lenormand T, Dutheil J. 2005. Recombination difference between sexes: a role for haploid selection. PLoS Biol 3: e63.

Li H., Durbin R. 2009. Fast and Accurate Short Read Alignment with Burrows-Wheeler Transform. Bioinformatics 25: 1754–60.

Lin Y-C et al. 2018. Functional and evolutionary genomic inferences in Populus through genome and population sequencing of American and European aspen. P Natl Acad Sci USA 115: E10970–E10978.

Luquez V et al. 2008. Natural phenological variation in aspen (Populus tremula): the SwAsp collection. Tree genet genomes 4: 279–292.

Mandel M, Marmur J. 1963. Use of ultraviolet absorbance-temperature profile for determining the quanine plus cytosine content of DNA. Methods Enzymol 518 (1962): 195–206.

Ma T et al. 2013. Genomic insights into salt adaptation in a desert poplar. Nat Commun 4: 2797.

Marand AP et al. 2019. Historical Meiotic Crossover Hotspots Fueled Patterns of Evolutionary Divergence in Rice. Plant Cell 31: 645–662.

Margarido GRA, Souza AP, Garcia AAF. 2007 OneMap: Software for Genetic Mapping in Outcrossing Species. Hereditas 144: 78–79.

Martin M. 2011. Cutadapt removes adapter sequences from high-throughput sequencing reads. EMBnet.journal 17: 10–12.

McKenna A et al. 2010. The Genome Analysis Toolkit: A MapReduce Framework for Analyzing Next-Generation DNA Sequencing Data. Genome Res 20: 1297–1303.

McVean GAT et al. 2004. The fine-scale structure of recombination rate variation in the human genome. Science 304: 581–584.

Nachman MW. 2002. Variation in recombination rate across the genome: evidence and implications. Curr opin genet dev 12: 657–663.

Narasimhan V et al. 2016. BCFtools/RoH: a hidden Markov model approach for detecting autozygosity from next-generation sequencing data. Bioinformatics 32;11: 1749–1751.

Neph S et al. 2012. BEDOPS: high-performance genomic feature operations. Bioinfomatics 28(14): 1919–1920.

Paolucci I et al. 2010. Genetic linkage maps of Populus alba L. and comparative mapping analysis of sex determination across Populus species. Tree Gen Genomes 6: 863.

Quinlan AR. 2014. BEDTools: The Swiss-Army Tool for Genome Feature Analysis. Current Protocols in Bioinformatics 47: 11.12.1–11.12.34.

R Core Team. 2018. R: A language and environment for statistical computing. R Foundation for statistical Computing, Vienna, Austria.

Rezvoy C, Charif D, Guéquen L, Marais GAB. 2007. MareyMap: an R-based tool with graphical inter-face for estimation recombination rates. Bioinformatics 23(16): 2188–2189

Rodgers-Melnick E et al. 2015. Recombination in diverse maize is stable, predictable, and associated with genetic load. P Natl Acad Sci USA 112: 3823–3828.

Saze H, Kakutani T. 2011. Differentiation of epigenetic modifications between transposons and genes. Curr opin plant biol 14: 81–87.

Schiffthaler B, Bernhardsson C, Ingvarsson PK, Street NR. 2017. BatchMap: A parallel implementation of the OneMap R package for fast computation of F1 linkage maps in outcrossing species. PLoS One 12: e0189256.

Silva-Junior OB, Grattapaglia D. 2015. Genome-wide patterns of recombination, linkage disequilibrium and nucleotide diversity from pooled resequencing and single nucleotide polymorphism genotyping unlock the evolutionary history of Eucalyptus grandis. New Phytol 208: 830–845.

Si W et al. 2015. Widely distributed hot and cold spots in meiotic recombination as shown by the sequencing of rice F 2 plants. New Phytol 206: 1491–1502.

Slavov GT et al. 2012. Genome Resequencing Reveals Multiscale Geographic Structure and Extensive Linkage Disequilibrium in the Forest Tree Populus Trichocarpa. New Phytol 196:713–25.

Stapley J, Feulner PGD, Johnston SE, Santure AW, Smadja CM. 2017. Variation in recombination frequency and distribution across eukaryotes: patterns and processes. Philos T R Soc B 372: 20160455–S14.

Tang H, Krishnakumar V, Li J, Zhang X. 2015. Jcvi: JCVI utility libraries. Zenodo. Available online: http://dx.doi.org/10.5281/zenodo.31631

Tang H et al. 2015. ALLMAPS: robust scaffold ordering based on multiple maps. Genome biol 16: 3.

Taylor G. 2002. Populus: arabidopsis for forestry. Do we need a model tree? Ann Bot-London 90(6): 681–689.

Tiley GP, Burleigh JG. 2015. The relationship of recombination rate, genome structure, and patterns of molecular evolution across angiosperms. BMC Evol Biol 15: 194.

Tong C et al. 2016. Construction of High-Density Linkage Maps of Populus deltoides x P. simonii Using Restriction-Site Associated DNA Sequencing. PLoS One 11(3): e0150692.

Tuskan GA et al. 2006. The Genome of Black Cottonwood, Populus Trichocarpa (Torr. & Gray). Science 313(5793): 1596–1604

Van Os H, Stam P, Visser RGF, Van Eck HJ. 2005. RECORD: A Novel Method for Ordering Loci on a Genetic Linkage Map. Theor Appl Genet 112: 30–40.

Wang J et al. 2018. A major locus controls local adaptation and adaptive life history variation in a perennial plant. Genome biol 19: 72.

Wang Z et al. 2014. Phylogeny Reconstruction and Hybrid Analysis of Populus (Salicaceae) Based on Nucleotide Sequences of Multiple Single-Copy Nuclear Genes and Plastid Fragments. PLoS One 9(8): e103645.

Wang J, Street NR, Scofield DG, Ingvarsson PK. 2016. Natural Selection and Recombination Rate Variation Shape Nucleotide Polymorphism Across the Genomes of Three Related Populus Species. Genetics 202: 1185–1200.

Wright S. 1931. Evolution in mendelian populations. Genetics 16: 97–159.

Zhang D et al. 2004. Genetic mapping in (Populus tomentosa x Populus bolleana) and P. tomentosa Carr. using AFLP markers Theor Appl Genet 108: 657.

Zhigunov AV et al. 2017. Development of F1 Hybrid Population and the High-Density Linkage Map for European Aspen (Populus Tremula L.) using RADseq Technology. BMC Plant Biol 17(Suppl 1): 180.

