## Supplementary Figures for "Constructing a high-density linkage map to infer the genomic landscape of recombination rate variation in European Aspen *(Populus tremula)*"

**Figure S1.** Comparison between the physical chromosomes of *P. trichocarpa* and the probe-marker position (in cM) of the *P. tremula* consensus LGs. LG – chromosome comparisons with more than 150 markers-pairs were considered as homologous and all LGs were therefore renamed to the homologous chromosome name in *P. trichocarpa*.

**Figure S2.** Circos plot of the consensus map. A) Marker distribution over the 19 chromosomes (Chr 1- Chr 19). Each black vertical line represents a marker (19,519 in total) in the map and is displayed according to the marker positions in cM. Track B-C visualizes multi marker scaffolds, where each line is a pairwise position comparison of probe-markers from the same scaffold. B) Position comparisons of probe-markers from the same scaffold that are located on the same chromosome. Grey lines indicate probe-markers that are located < 20 cM from each other and red lines indicate probe-markers >= 20 cM apart. C) Position comparisons of probe-markers from the same scaffold that are mapping to different chromosomes. Orange lines indicated probe-markers from the same scaffold split over 2 chromosomes, while dark blue lines indicated probe-markers split over 3 chromosomes.

**Figure S3.** Physical position (in bp) versus genetic position (in cM) for all probe-markers in the parental maps that are located on scaffolds showing signs of splits within a chromosome (maximum distance between probe markers from the same scaffold > 20 cM). Red dots indicate position in the female map while blue dots indicate position in the male map. Grey dots indicate probe-markers positioned on other chromosomes (these splits are further investigated in Figure S2). All gene-models positioned on the scaffold are visualized as horizontal black bars in the bottom on the figure. Green vertical bars indicate gap positions in the assembly (N:s). Blue or red vertical lines represent the assembly gap or artificial gap, respectively, that the scaffold was split at before creation of physical chromosomes with Allmaps. Scaffold names and chromosome belonging are represented in the title of each plot.

**Figure S4.** Physical position (in bp) versus chromosome grouping (1-19) for all probe-markers in the consensus map that are located on scaffolds showing signs of splits between chromosomes. All gene-models positioned on the scaffold are visualized as horizontal black bars in the bottom on the figure. Green vertical bars indicate gap positions in the assembly (N:s). Blue or red vertical lines represent the assembly gap or artificial gap, respectively, that the scaffold was split at before creation of physical chromosomes with Allmaps. Scaffold name is represented in the title of each plot.

**Figure S5.** Scaffolds showing the breakpoint splits within gene-models. The genomic positions of gene-models (exons plus introns) are represented with thin black lines, where gene models on the plus strand are represented above gene models on the minus strand. Exons from gene-models on the plus strand are represented with red boxes while exons from gene-models on the minus strand are represented with green boxes. Breakpoint decisions are represented with blue vertical lines. Scaffold name is represented in the title of each plot.

**Figure S6.** Genetic maps and resulting physical map created by Allmaps. For each of the 19 chromosomes, the left panel shows the marker distribution (in cM) for the genetic maps and the anchored genomic region (in Mb) for the physical map, while the right panel is showing the correspondence between the physical (x-axis) and recombination-based (y-axis) position of markers. The female map is depicted in green and the male map is depicted in orange.

**Figure S7.** Summary statistics for genetic linkage map based (purple) and sequenced based (orange) estimates of recombination rate (cM/Mb) in windows of 1 Mb. Red dots correspond to the average recombination rate over all windows and grey dots are outliers.

**Figure S8.** Manhattan plots for all genomic features.

**Figure S9.** Correlation between LD-based and map-based recombination rate

Supplementary Figure 1

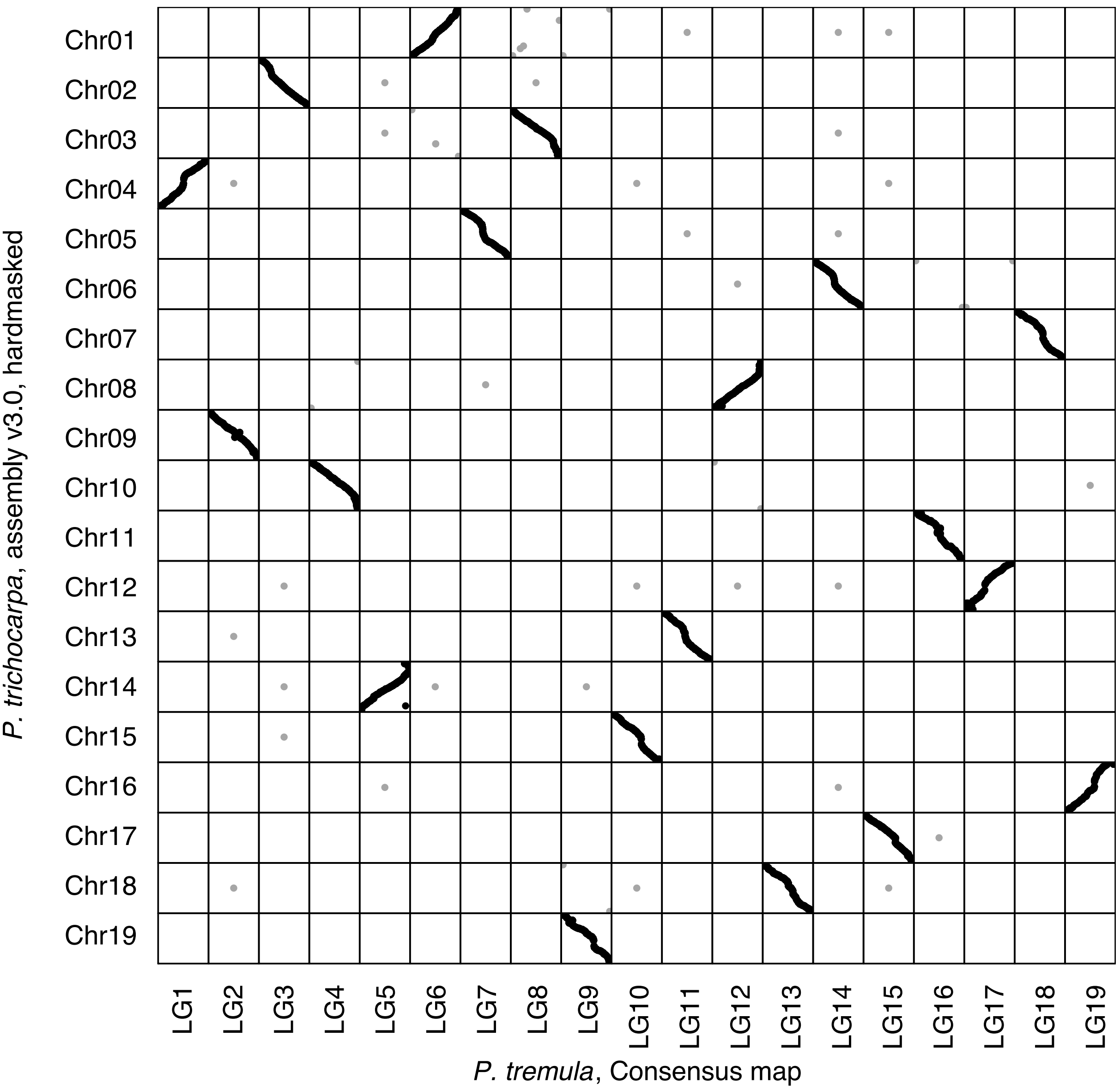

Supplementary Figure 2

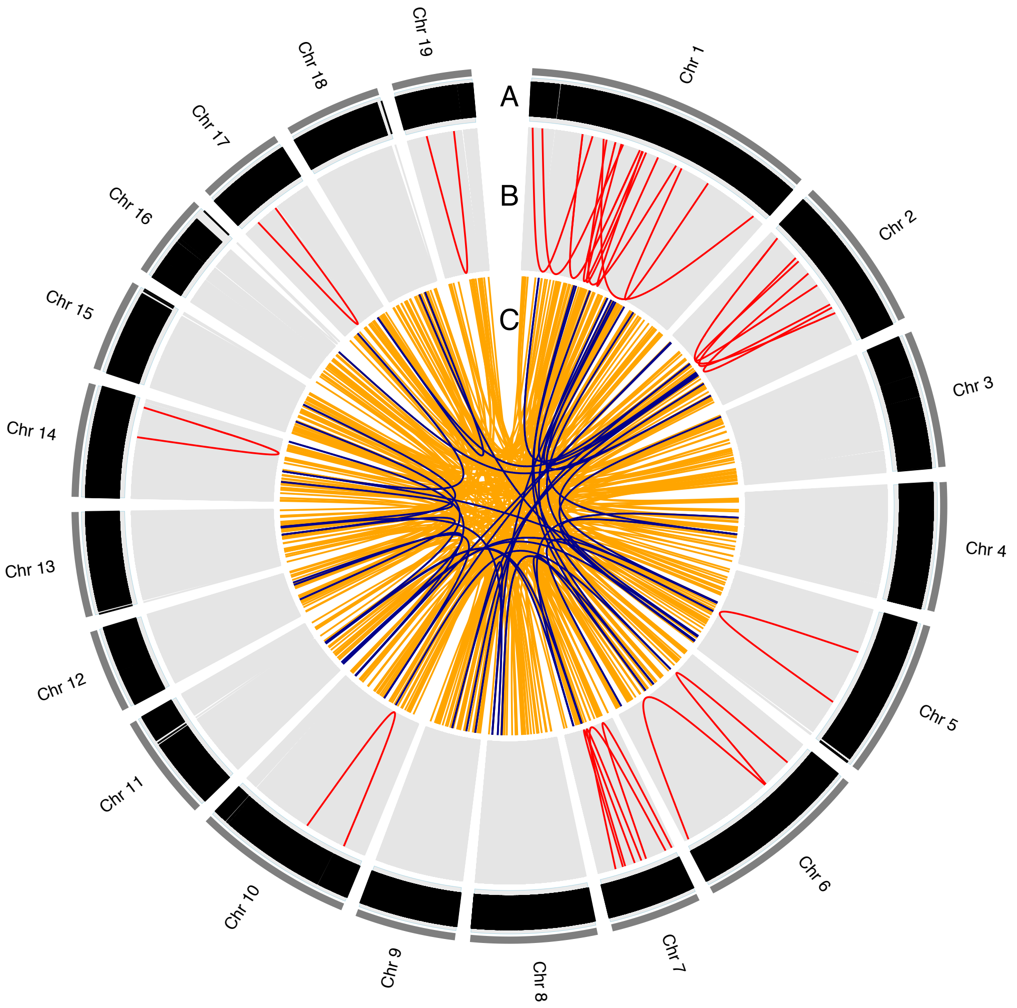

Supplementary Figure 3

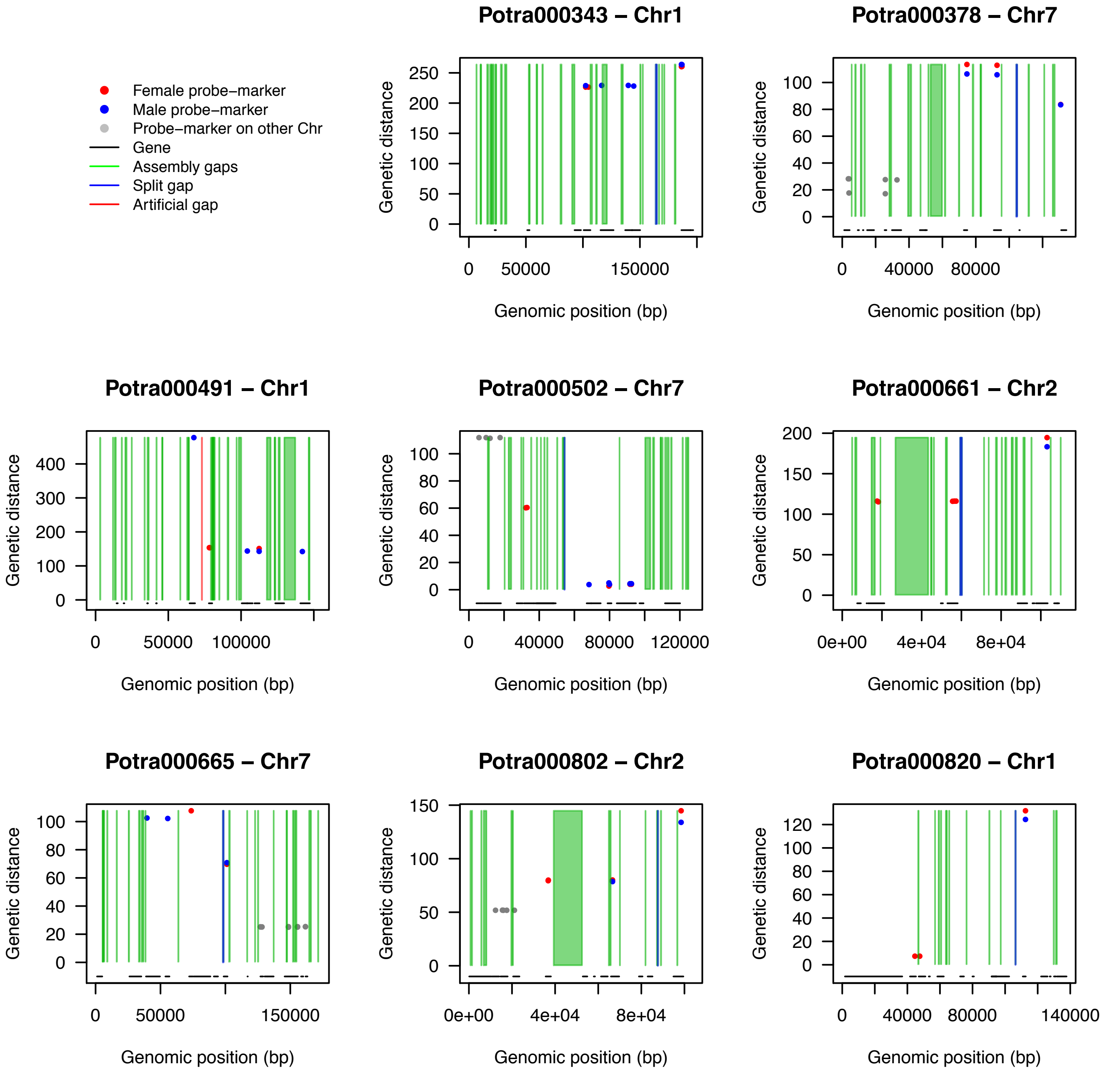
Female probe−marker
Male probe−marker
Probe−marker on other Chr
Gene
Assembly gaps
Split gap
Artificial gap
Potra000343 – Chr1
Potra000378 – Chr7
Potra000491 – Chr1
Potra000502 – Chr7
Potra000661 – Chr2
Potra000665 – Chr7
Potra000802 – Chr2
Potra000820 – Chr1
Genetic distance
Genomic position (bp)

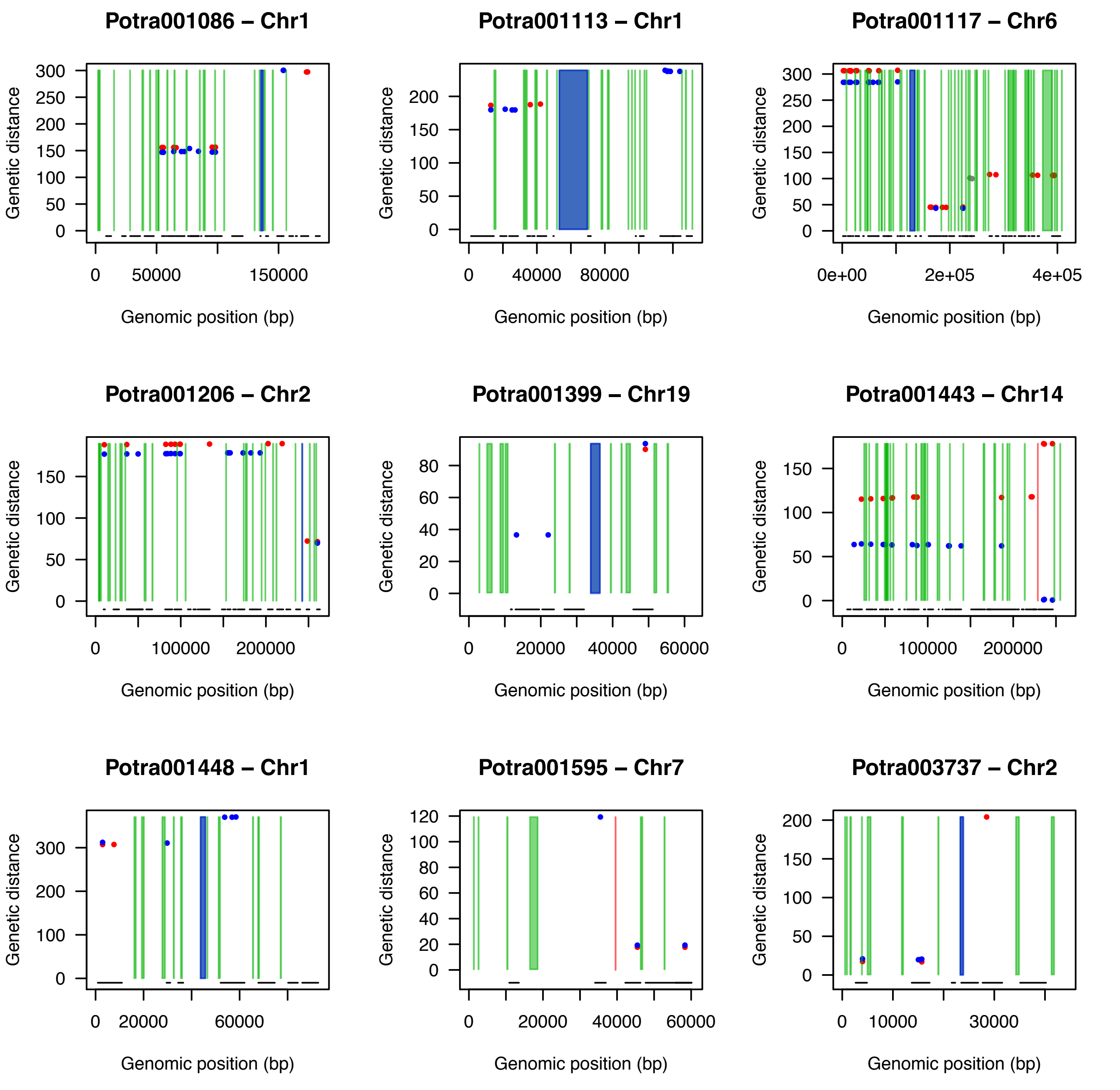
Potra001086 – Chr1
Potra001113 – Chr1
Potra001117 – Chr6
Potra001206 – Chr2
Potra001399 – Chr19
Potra001443 – Chr14
Potra001448 – Chr1
Potra001595 – Chr7
Potra003737 – Chr2
Genetic distance
Genomic position (bp)

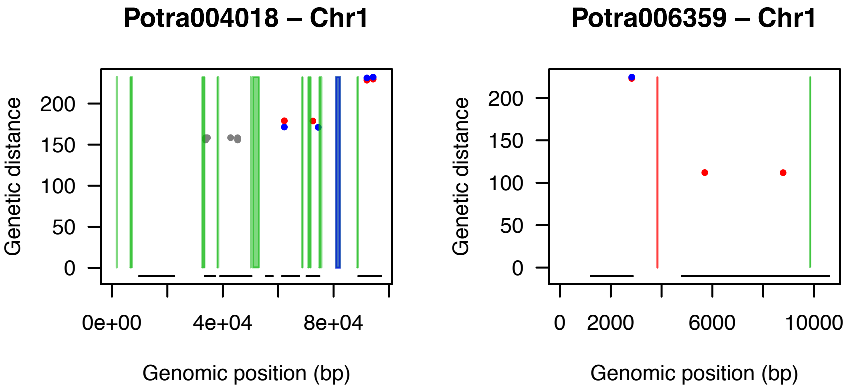
Potra004018 – Chr1
Genetic distance
0
50
100
150
200
0e+00
4e+04
8e+04
Genomic position (bp)
Potra006359 – Chr1
Genetic distance
0
50
100
150
200
0
2000
6000
10000
Genomic position (bp)

Supplementary Figure 4

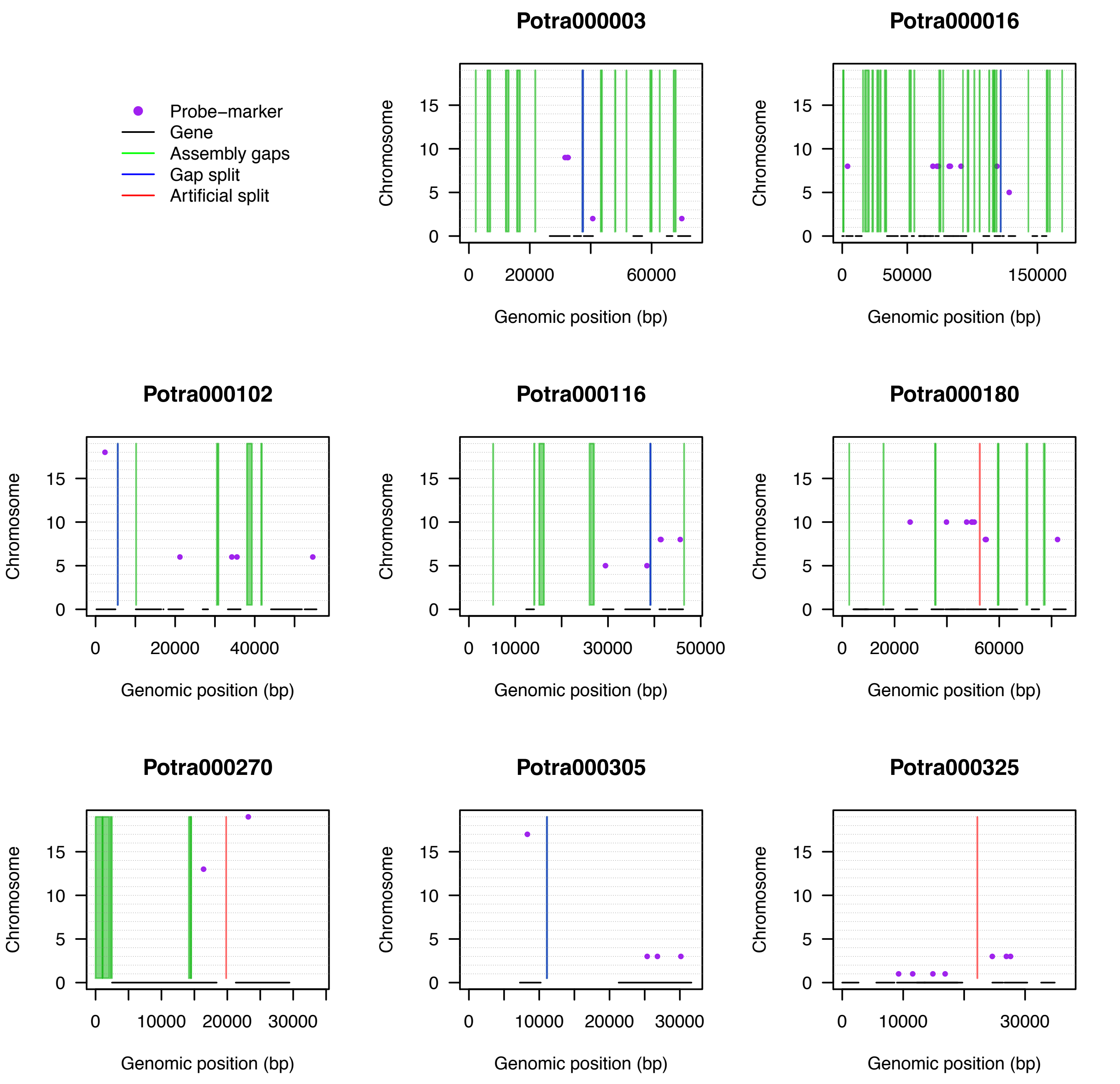

Probe−marker
Gene
Assembly gaps
Gap split
Artificial split
Potra000003
Potra000016
Potra000102
Potra000116
Potra000180
Potra000270
Potra000305
Potra000325
Chromosome
Genomic position (bp)

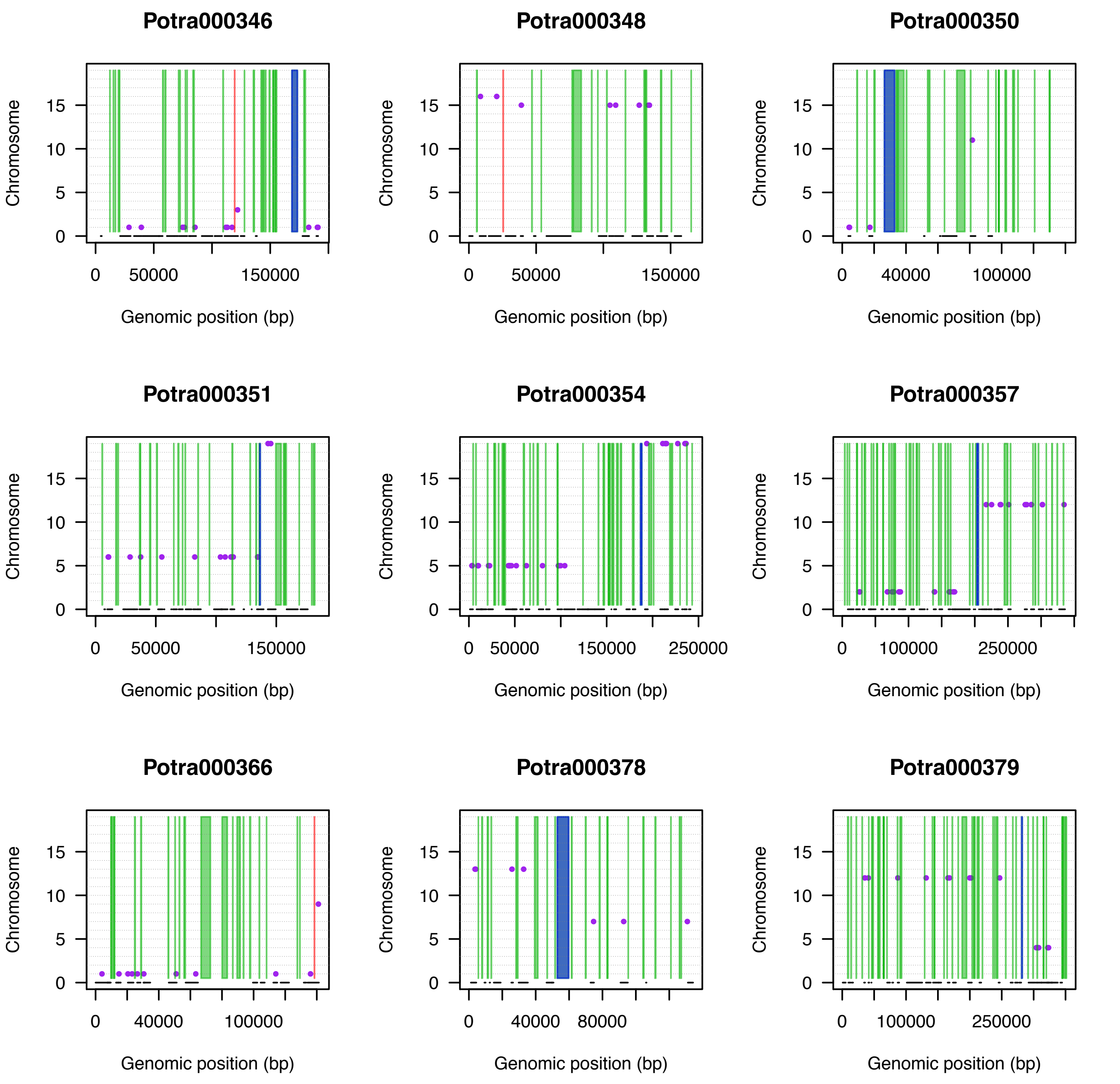
Potra000346
Potra000348
Potra000350
Potra000351
Potra000354
Potra000357
Potra000366
Potra000378
Potra000379
Chromosome
0
5
10
15
Genomic position (bp)
0
50000
150000
0
50000
150000
0
40000
100000
0
50000
150000
0
50000
150000
250000
0
100000
250000
0
40000
100000
0
40000
80000
0
100000
250000

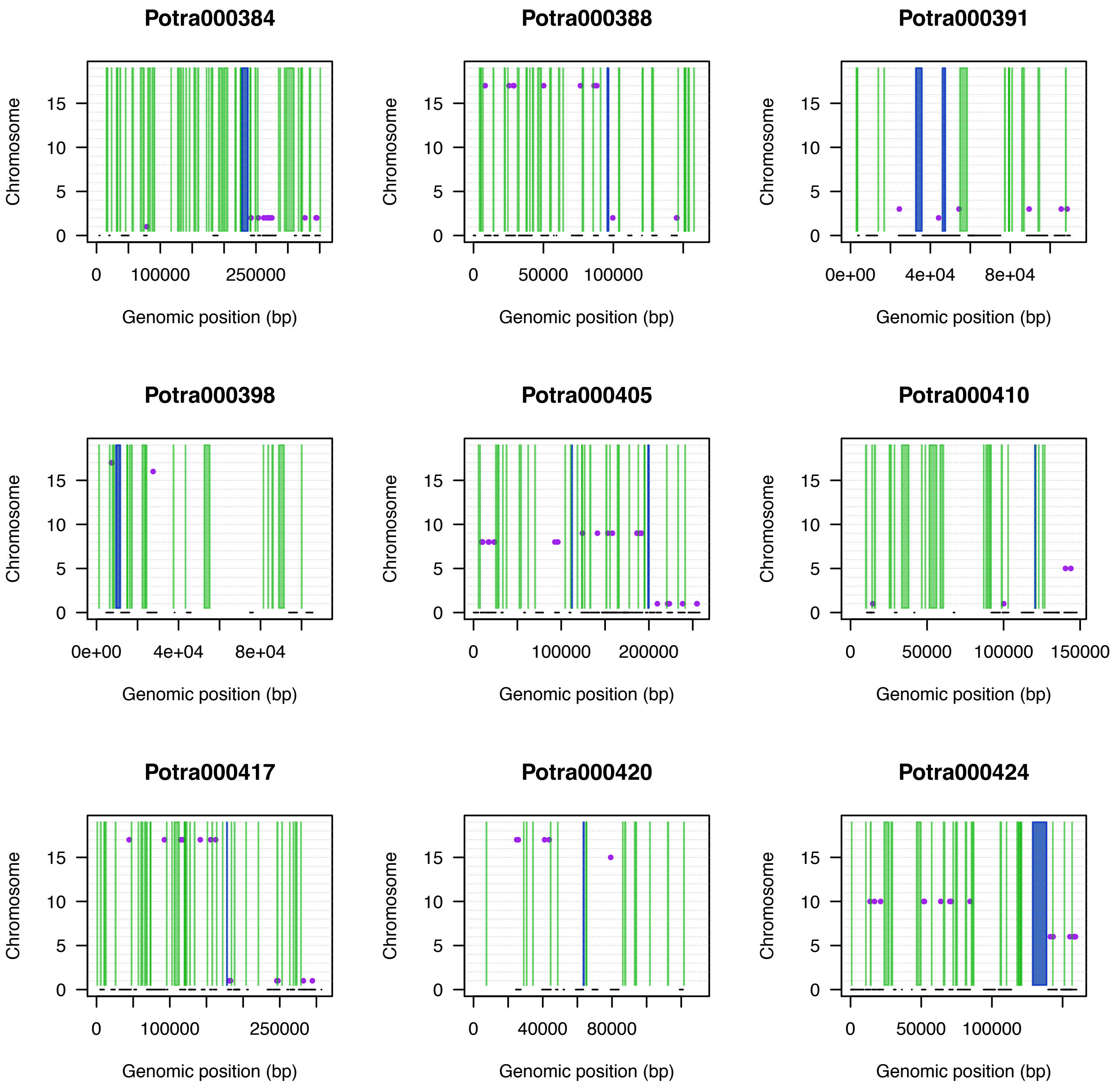
Potra000384
Potra000388
Potra000391
Potra000398
Potra000405
Potra000410
Potra000417
Potra000420
Potra000424
Chromosome
0
5
10
15
Genomic position (bp)
0
100000
250000
50000
0e+00
4e+04
8e+04
200000
150000
40000
80000

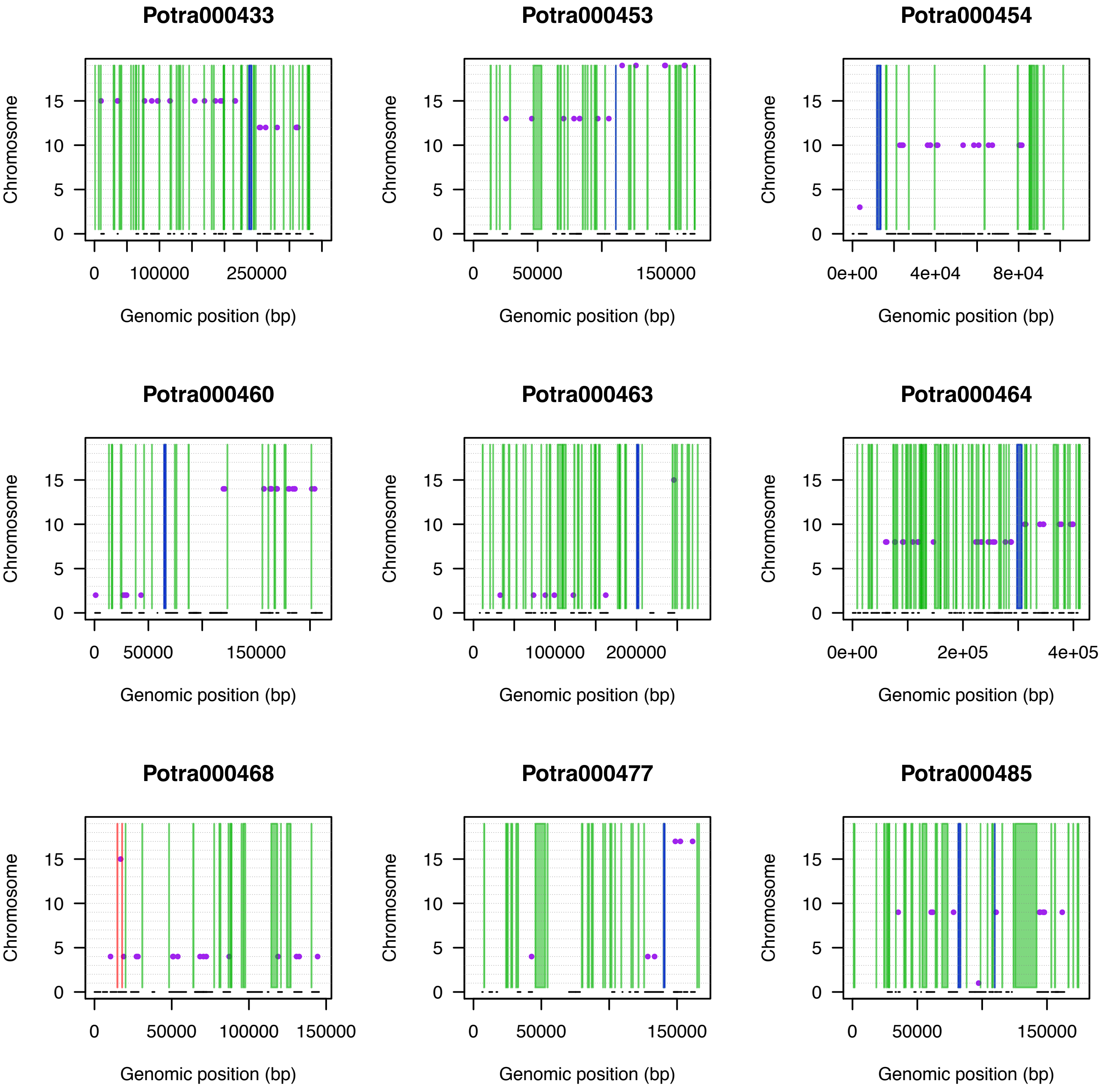
Potra000433
Potra000453
Potra000454
Potra000460
Potra000463
Potra000464
Potra000468
Potra000477
Potra000485
Chromosome
0
5
10
15
0
100000
250000
50000
150000
0e+00
4e+04
8e+04
200000
2e+05
4e+05
Genomic position (bp)

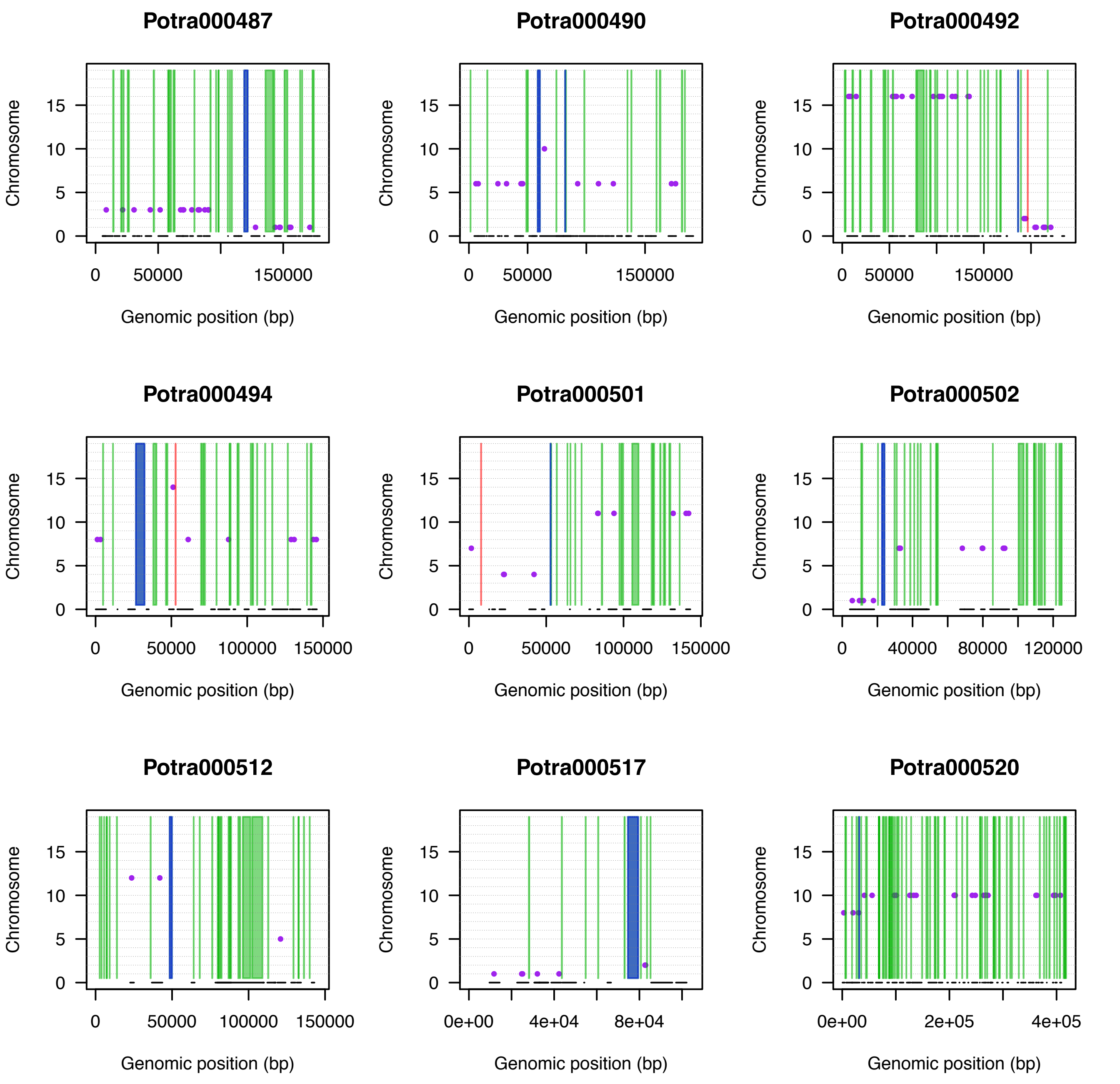
Potra000487
Potra000490
Potra000492
Potra000494
Potra000501
Potra000502
Potra000512
Potra000517
Potra000520
Chromosome
Genomic position (bp)
0
5
10
15
0
50000
100000
150000
0
40000
80000
120000
0e+00
4e+04
8e+04
2e+05
4e+05

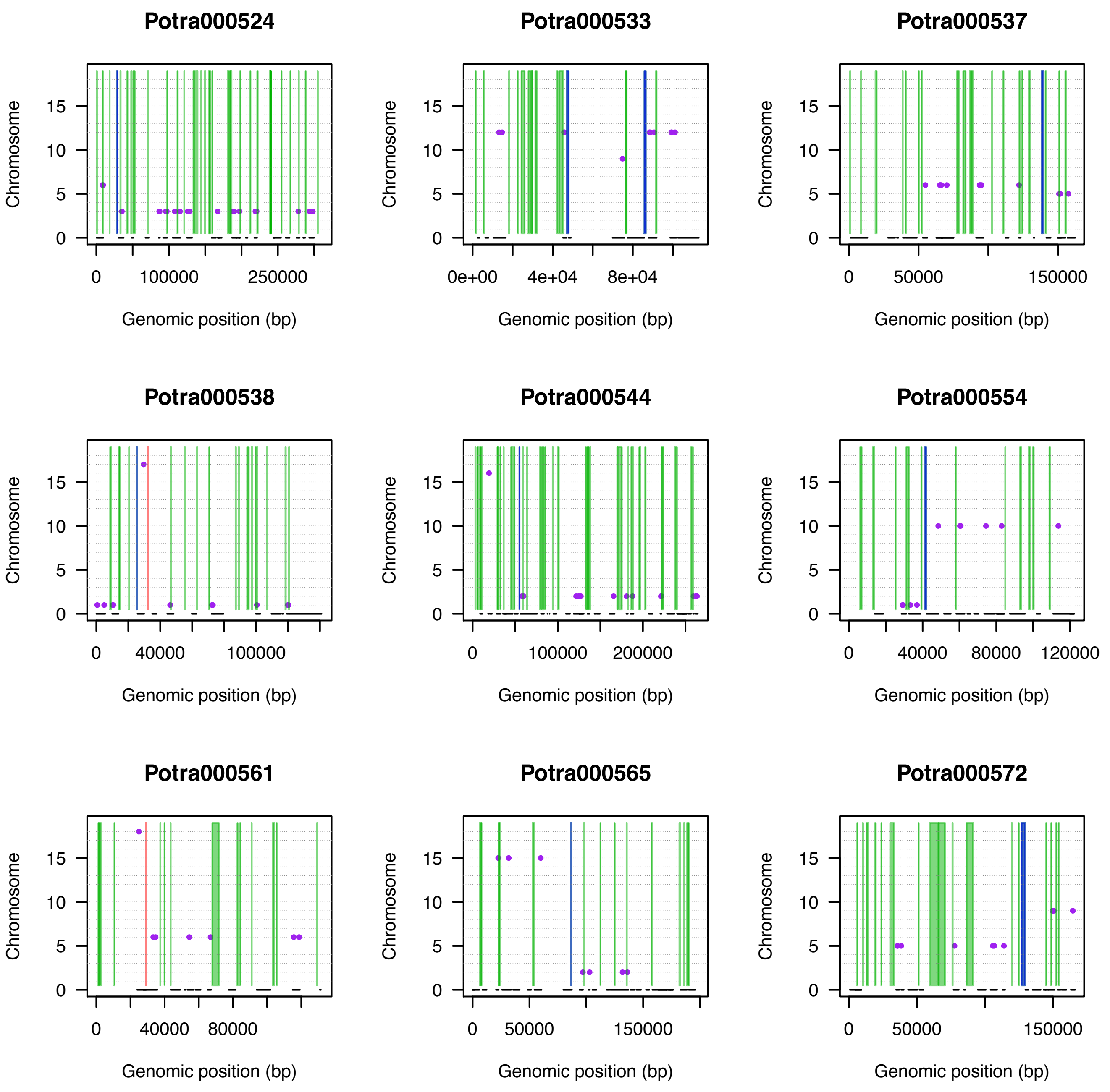

Potra000524
Potra000533
Potra000537
Potra000538
Potra000544
Potra000554
Potra000561
Potra000565
Potra000572
Chromosome
0
5
10
15
Genomic position (bp)
0
100000
250000
0e+00
4e+04
8e+04
50000
150000
40000
200000
80000
120000

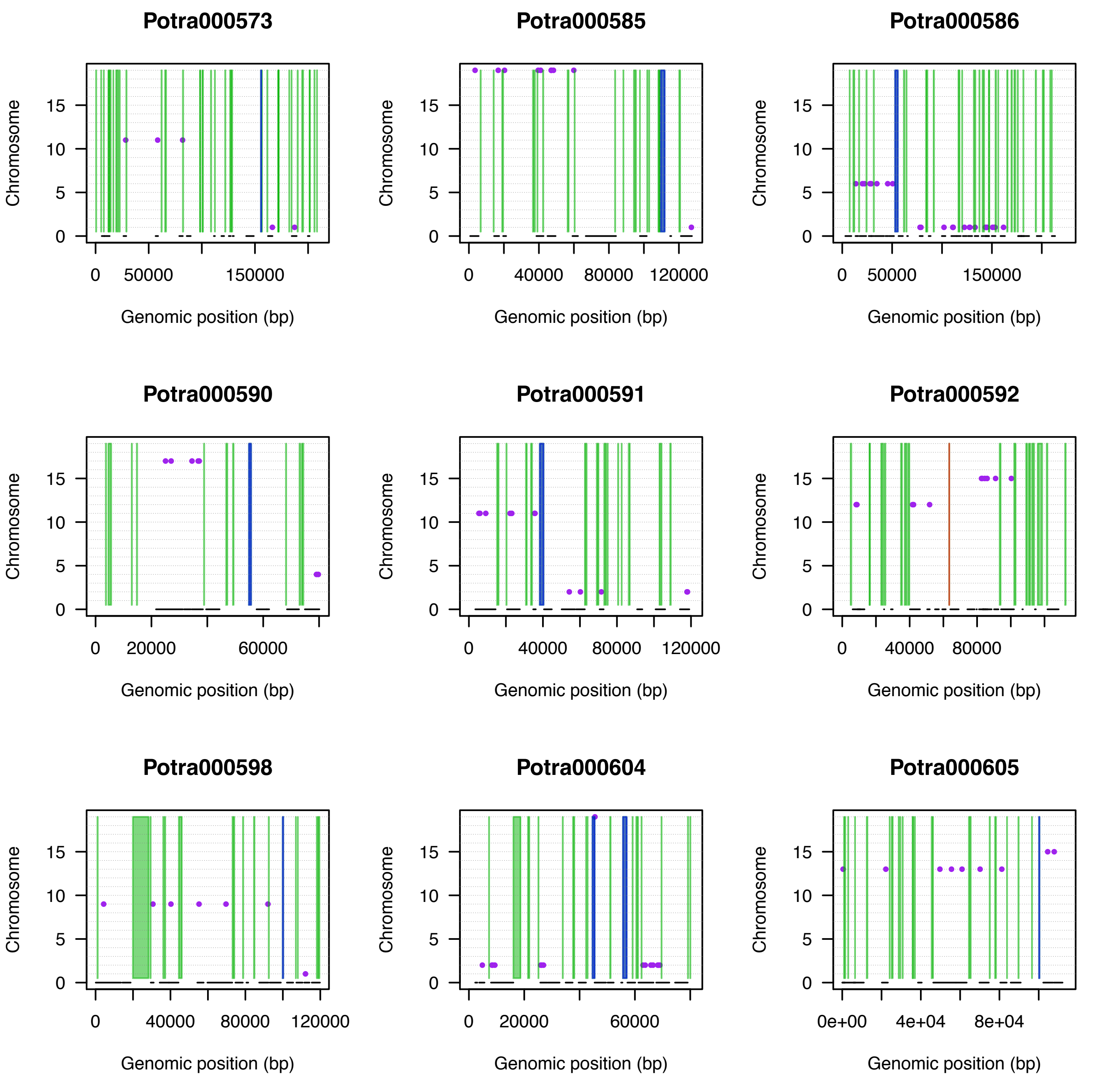

Potra000573
Potra000585
Potra000586
Potra000590
Potra000591
Potra000592
Potra000598
Potra000604
Potra000605
Chromosome
Genomic position (bp)
0
5
10
15
0
50000
150000
40000
80000
120000
20000
60000
0e+00
4e+04
8e+04

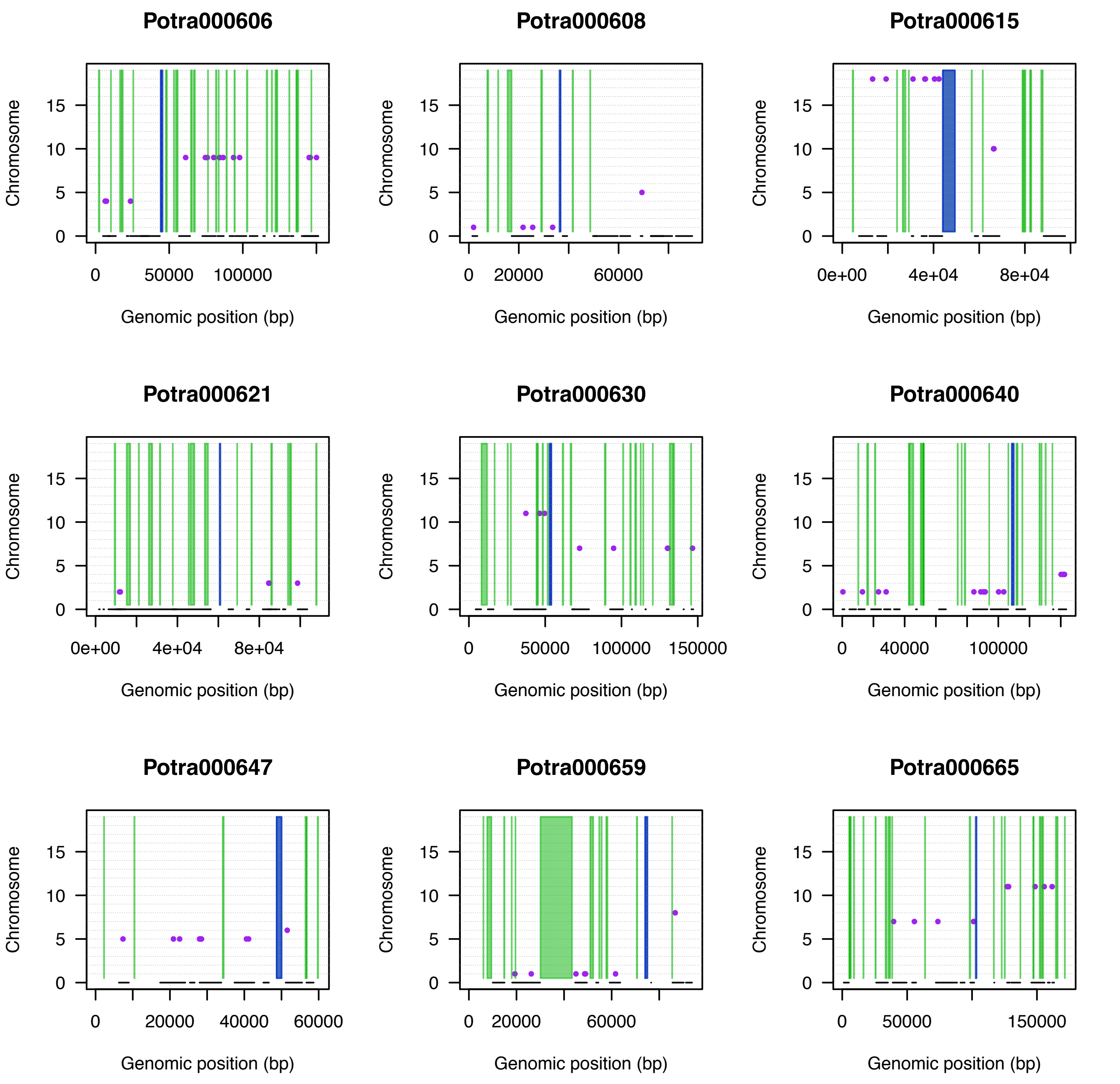

Potra000606
Potra000608
Potra000615
Potra000621
Potra000630
Potra000640
Potra000647
Potra000659
Potra000665
Chromosome
Genomic position (bp)

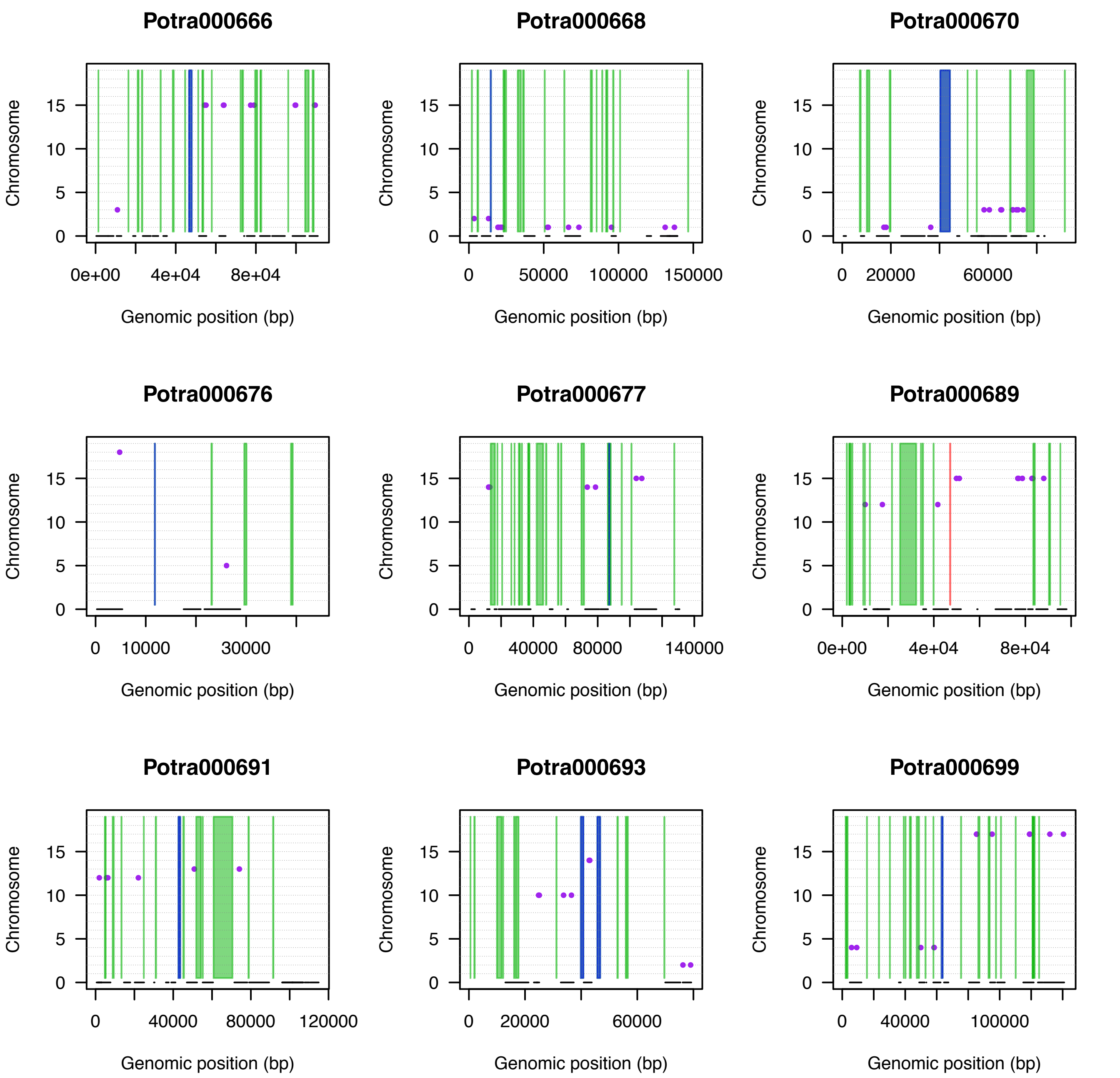

Potra000666
Potra000668
Potra000670
Potra000676
Potra000677
Potra000689
Potra000691
Potra000693
Potra000699
Chromosome
0
5
10
15
Genomic position (bp)
0e+00
4e+04
8e+04
0
50000
100000
150000
20000
60000
10000
30000
40000
80000
140000
120000
100000

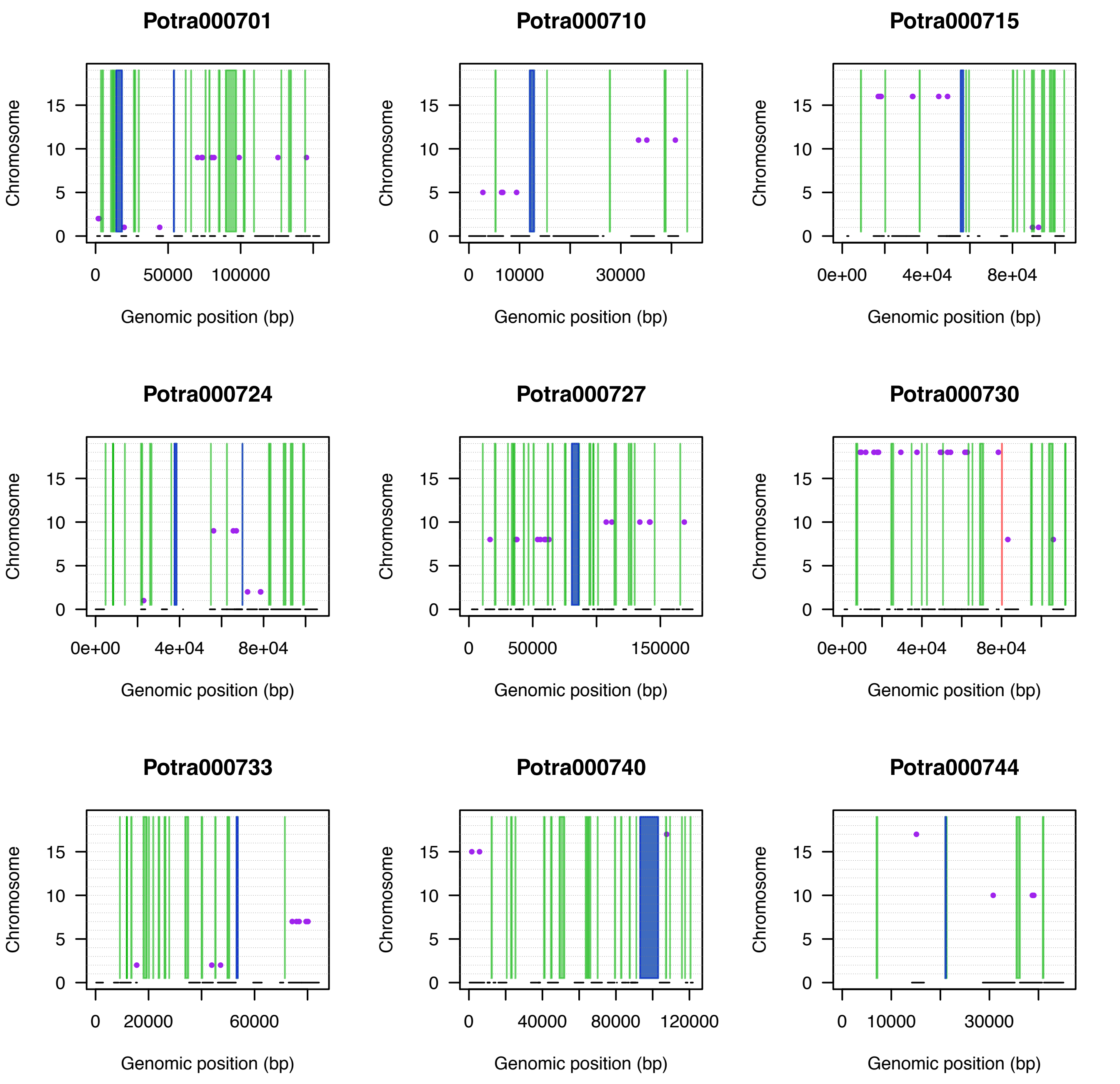

Potra000701
Potra000710
Potra000715
Potra000724
Potra000727
Potra000730
Potra000733
Potra000740
Potra000744
Chromosome
0
5
10
15
Genomic position (bp)
0
50000
100000
10000
30000
0e+00
4e+04
8e+04
150000
20000
60000
40000
80000
120000

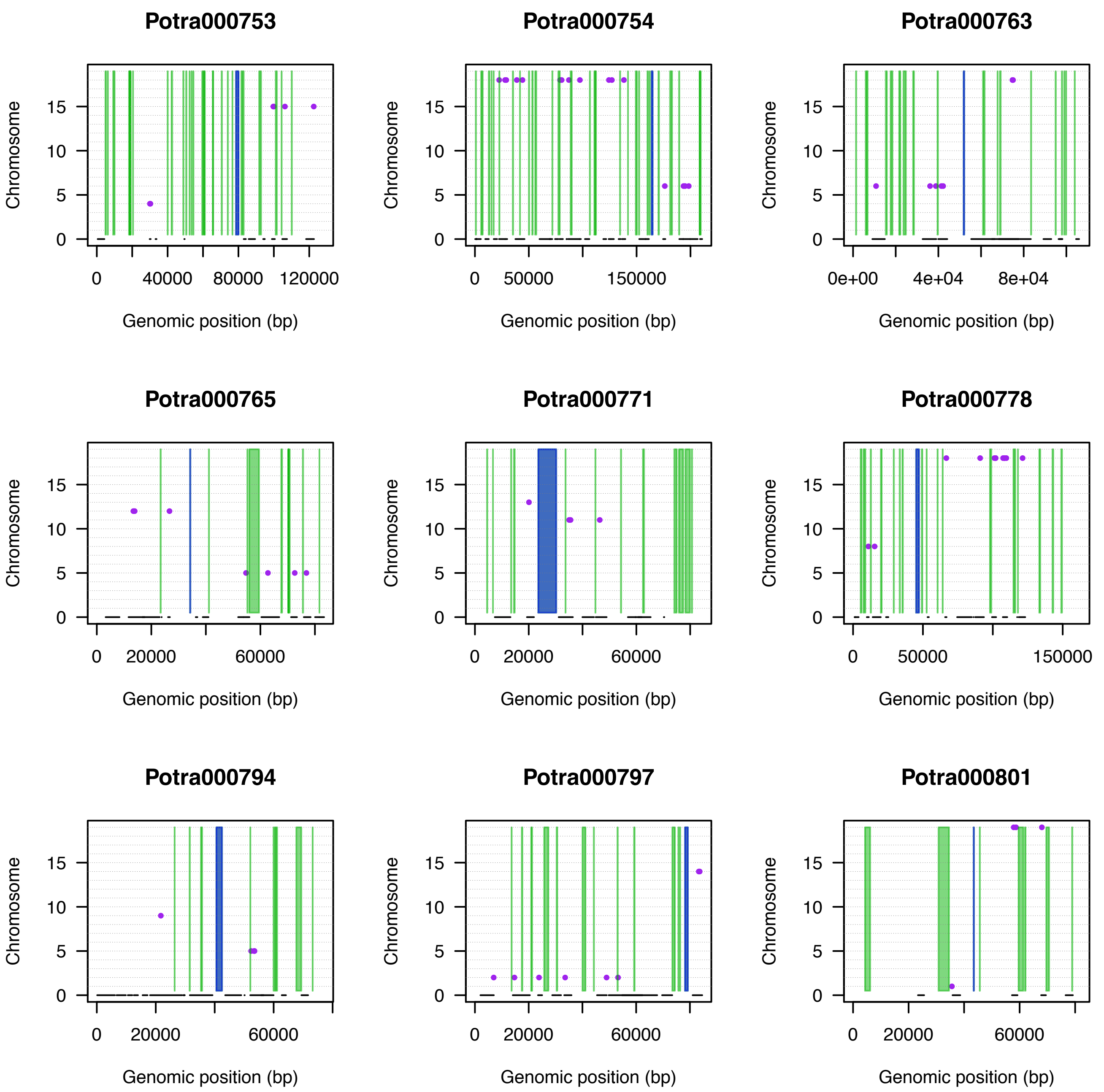

Potra000753
Potra000754
Potra000763
Potra000765
Potra000771
Potra000778
Potra000794
Potra000797
Potra000801
Chromosome
0
5
10
15
Genomic position (bp)
0
40000
80000
120000
50000
150000
0e+00
4e+04
8e+04
20000
60000

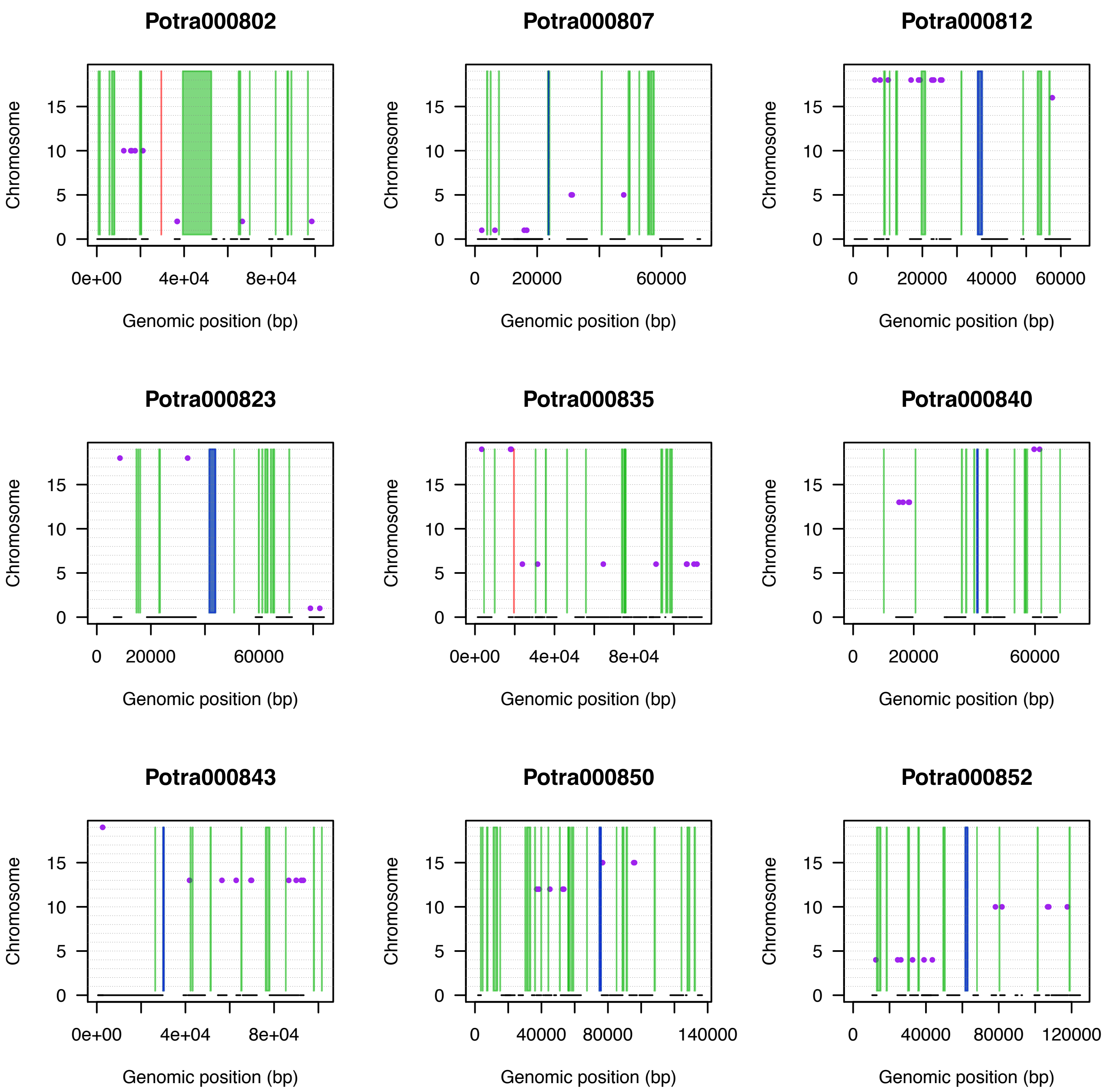

Potra000802
Potra000807
Potra000812
Potra000823
Potra000835
Potra000840
Potra000843
Potra000850
Potra000852
Chromosome
Genomic position (bp)
0
5
10
15
0e+00
4e+04
8e+04
0
20000
40000
60000
80000
120000
140000

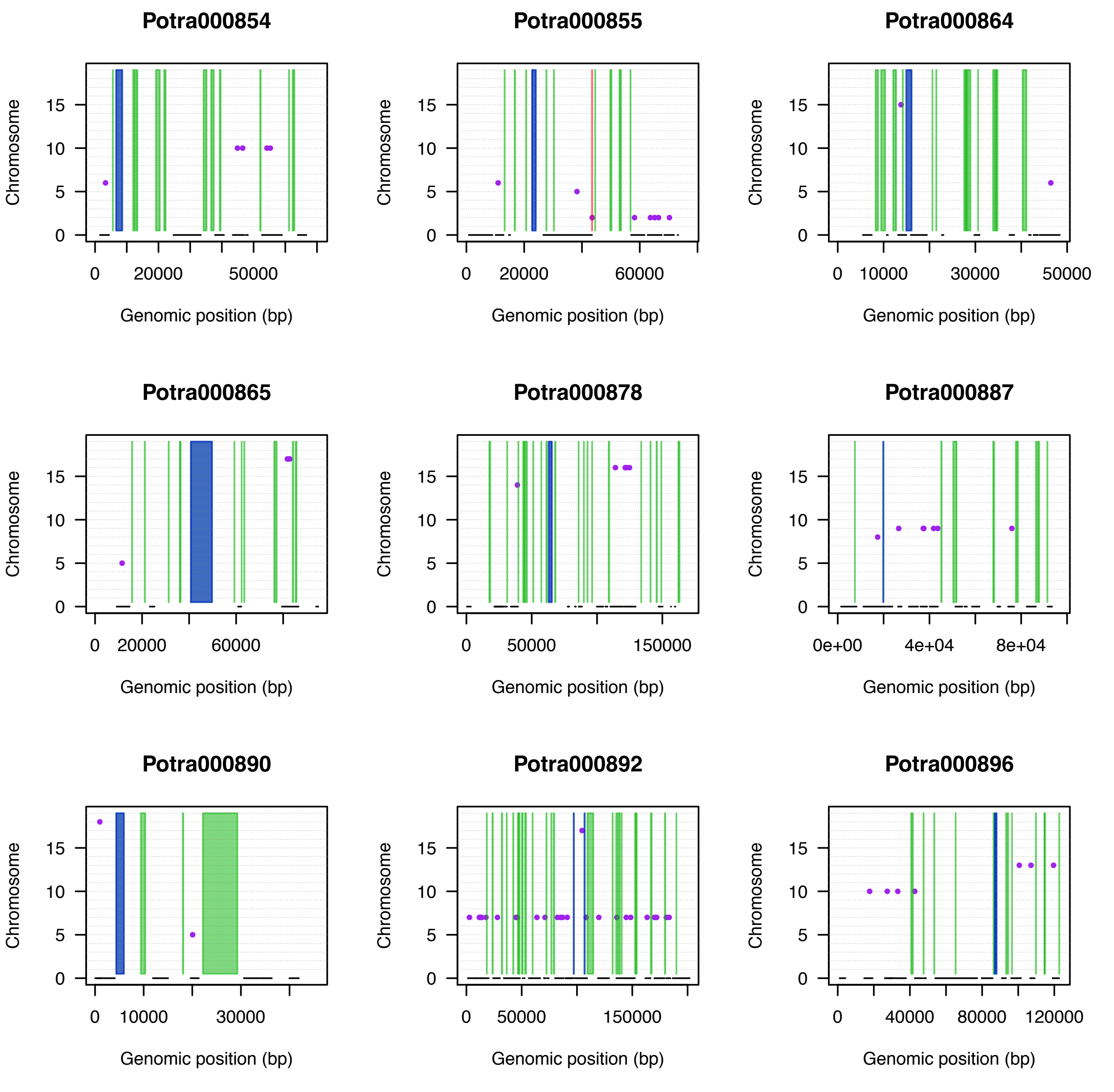

Potra000854
Potra000855
Potra000864
Potra000865
Potra000878
Potra000887
Potra000890
Potra000892
Potra000896
Chromosome
Genomic position (bp)

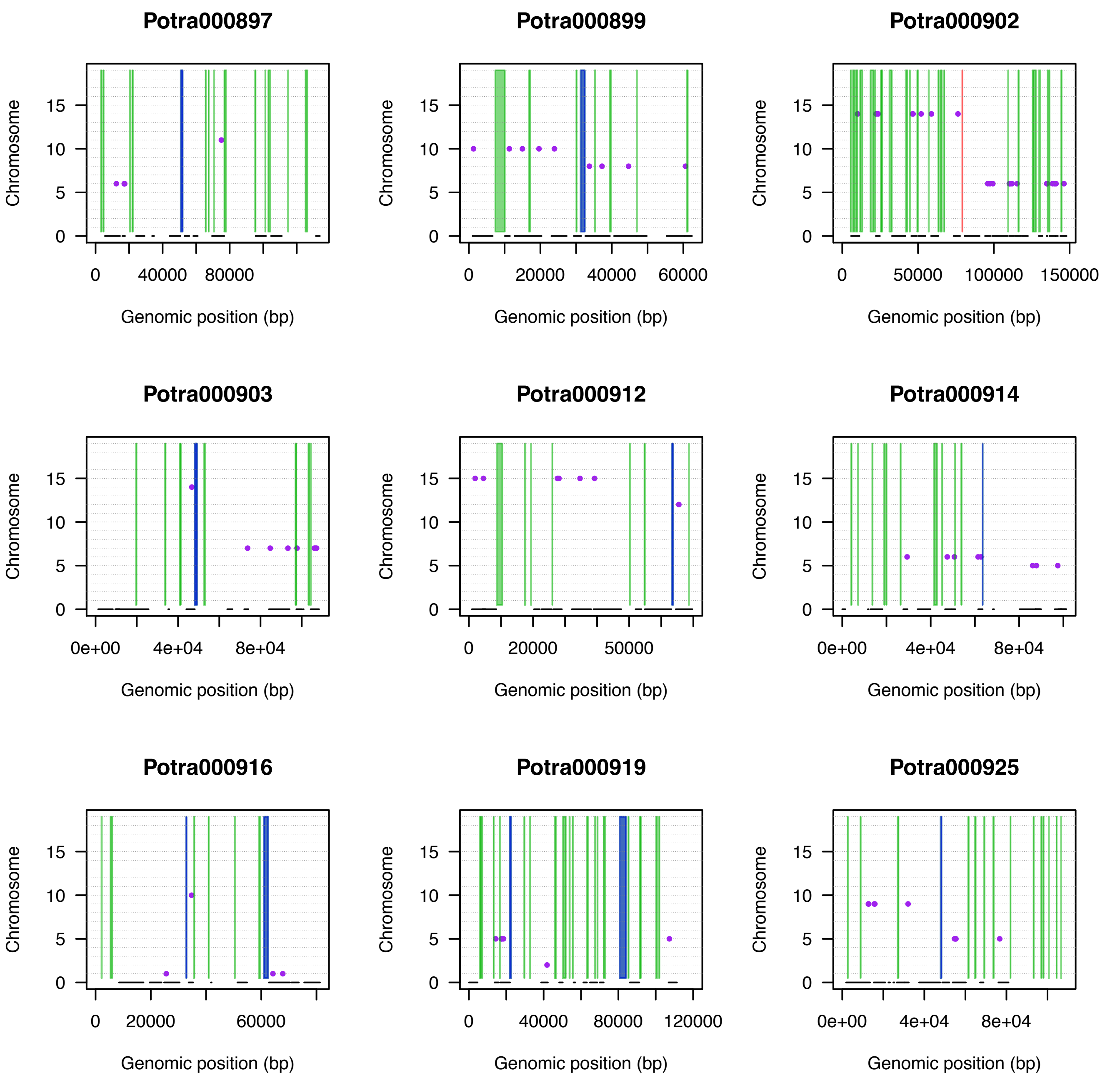

Potra000897
Potra000899
Potra000902
Potra000903
Potra000912
Potra000914
Potra000916
Potra000919
Potra000925
Chromosome
Genomic position (bp)

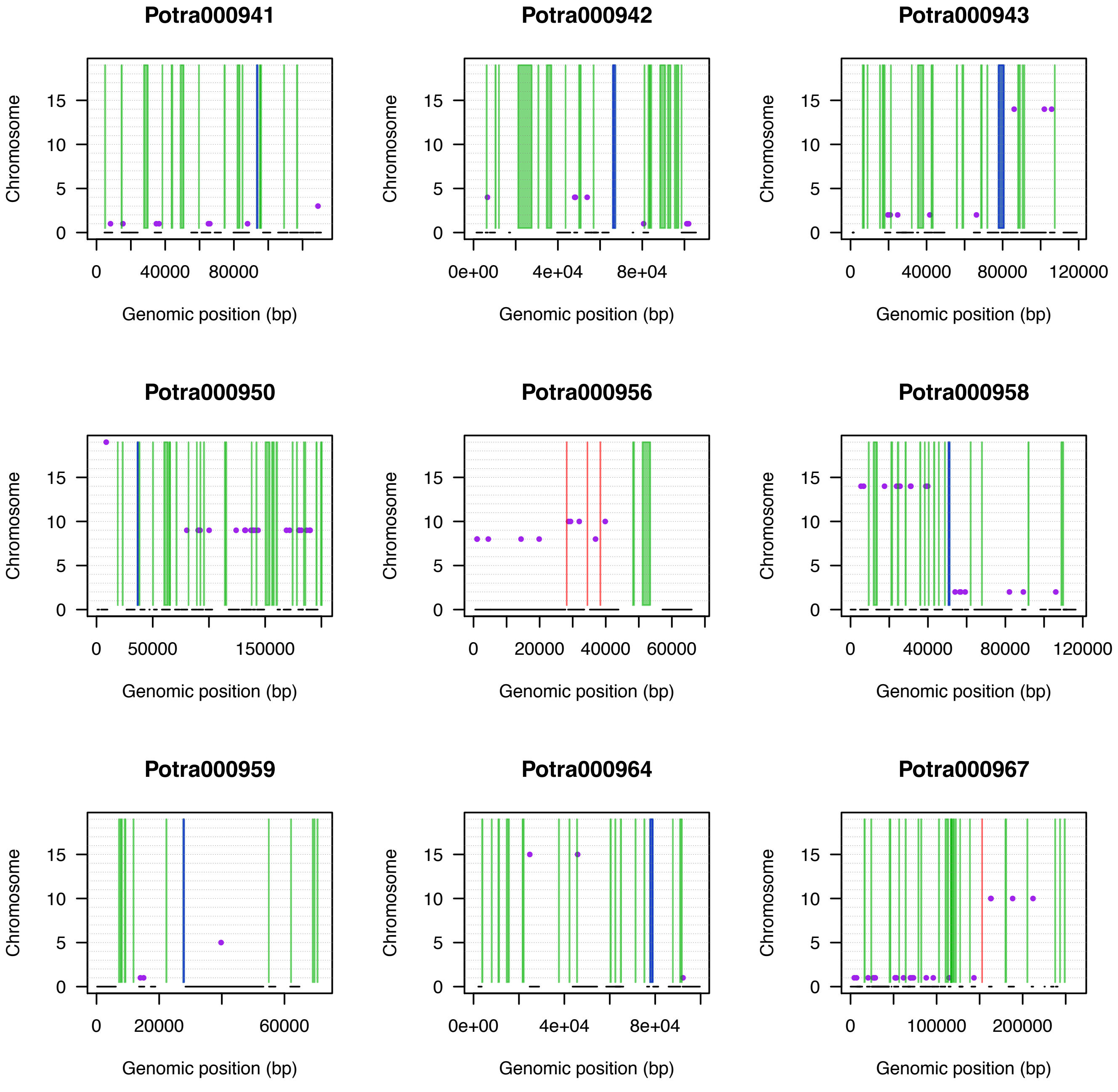

Potra000941
Potra000942
Potra000943
Potra000950
Potra000956
Potra000958
Potra000959
Potra000964
Potra000967
Chromosome
0
5
10
15
Genomic position (bp)
0
40000
80000
120000
0e+00
4e+04
8e+04
50000
150000
20000
60000
100000
200000

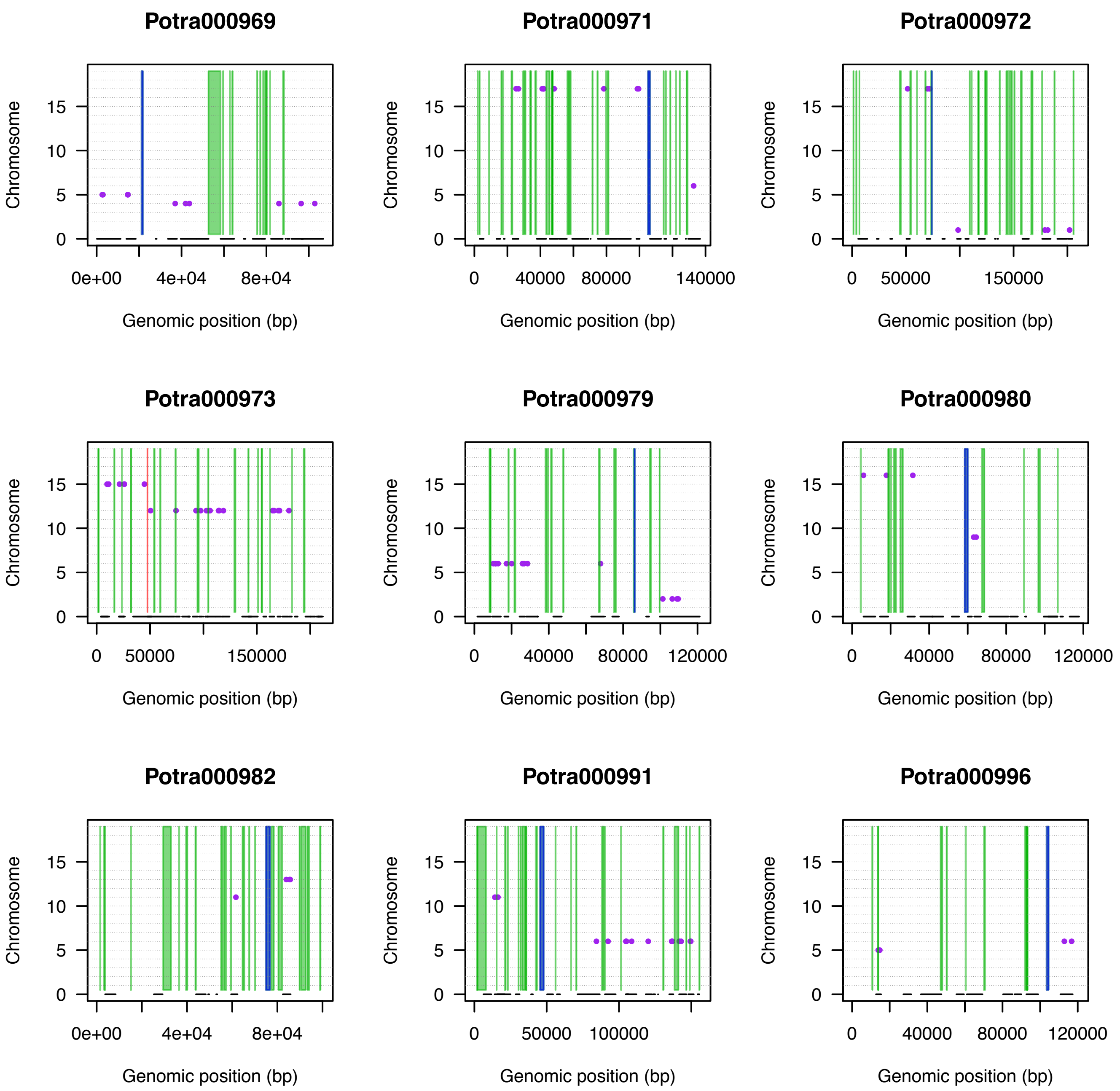
Potra000969
Potra000971
Potra000972
Potra000973
Potra000979
Potra000980
Potra000982
Potra000991
Potra000996
Chromosome
Genomic position (bp)
0
5
10
15
0e+00
4e+04
8e+04
0
40000
80000
140000
0
50000
150000
0
40000
80000
120000
0
50000
100000

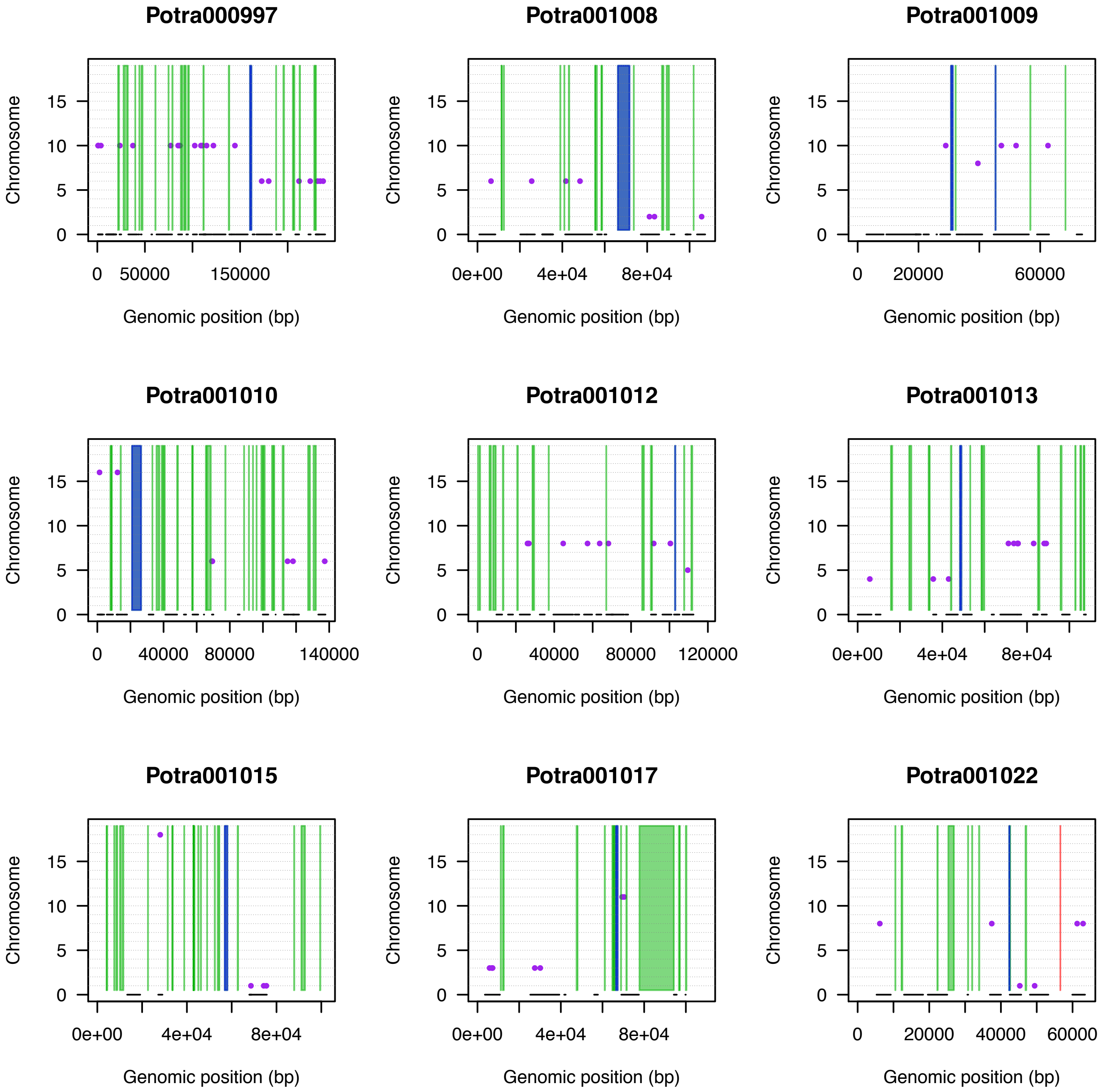
Potra000997
Potra001008
Potra001009
Potra001010
Potra001012
Potra001013
Potra001015
Potra001017
Potra001022
Chromosome
0
5
10
15
Genomic position (bp)
0
50000
150000
0e+00
4e+04
8e+04
20000
60000
40000
80000
140000
120000

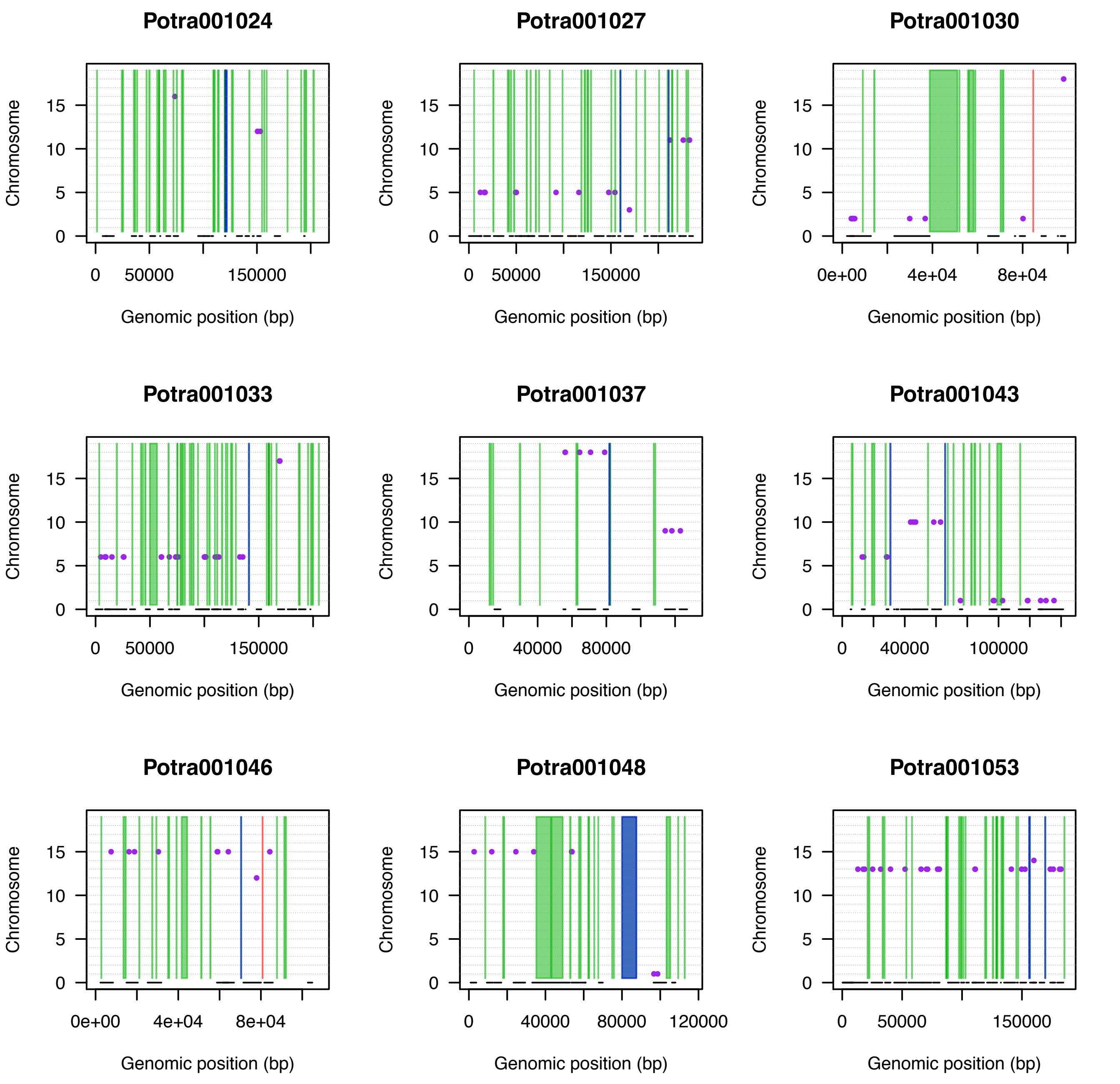
Potra001024
Potra001027
Potra001030
Potra001033
Potra001037
Potra001043
Potra001046
Potra001048
Potra001053
Chromosome
0
5
10
15
Genomic position (bp)
0
50000
150000
0e+00
4e+04
8e+04
40000
80000
100000
120000

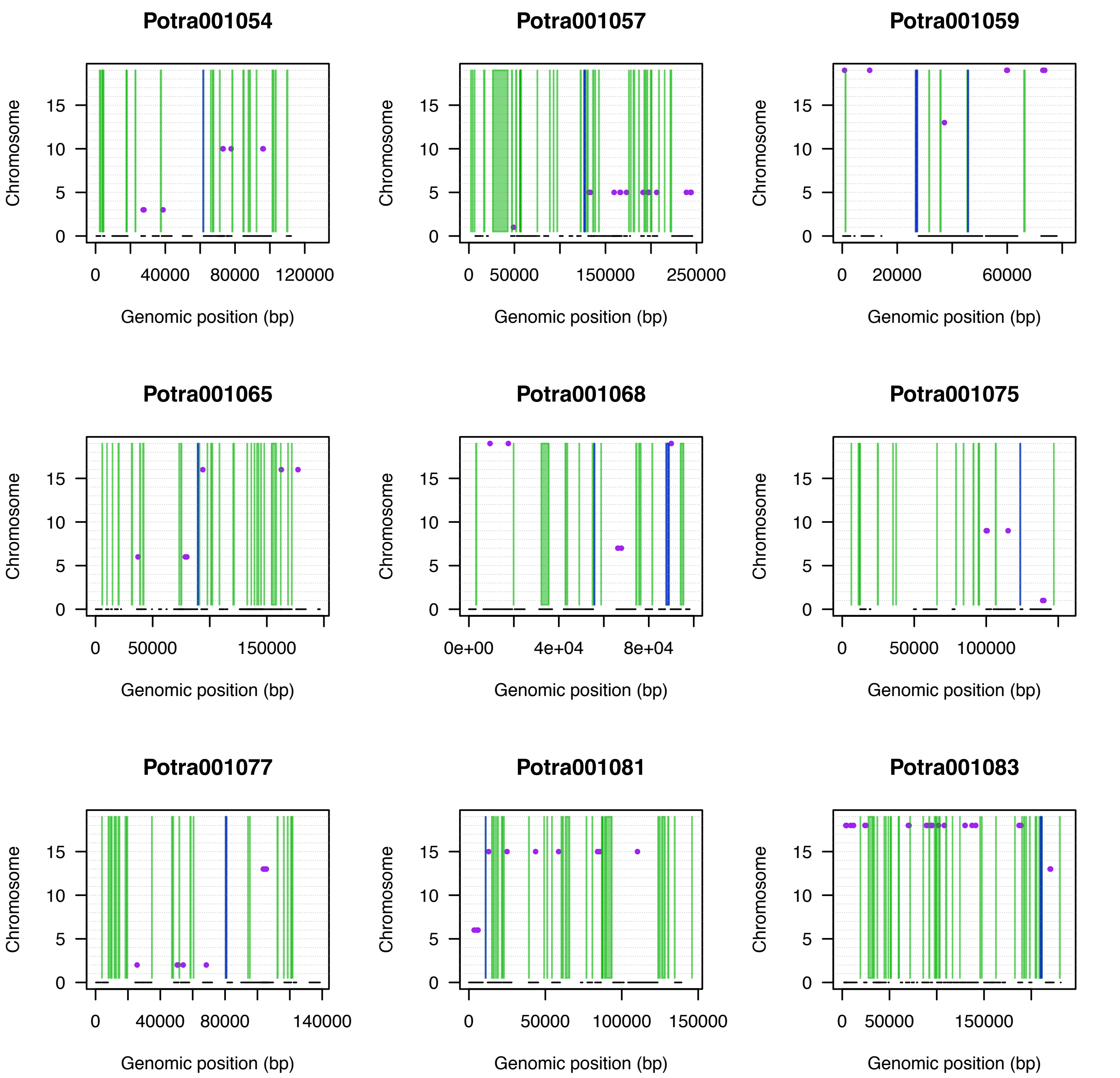
Potra001054
Potra001057
Potra001059
Potra001065
Potra001068
Potra001075
Potra001077
Potra001081
Potra001083
Chromosome
Genomic position (bp)
0
5
10
15
0
40000
80000
120000
50000
150000
250000
20000
60000
0e+00
4e+04
8e+04
100000
140000

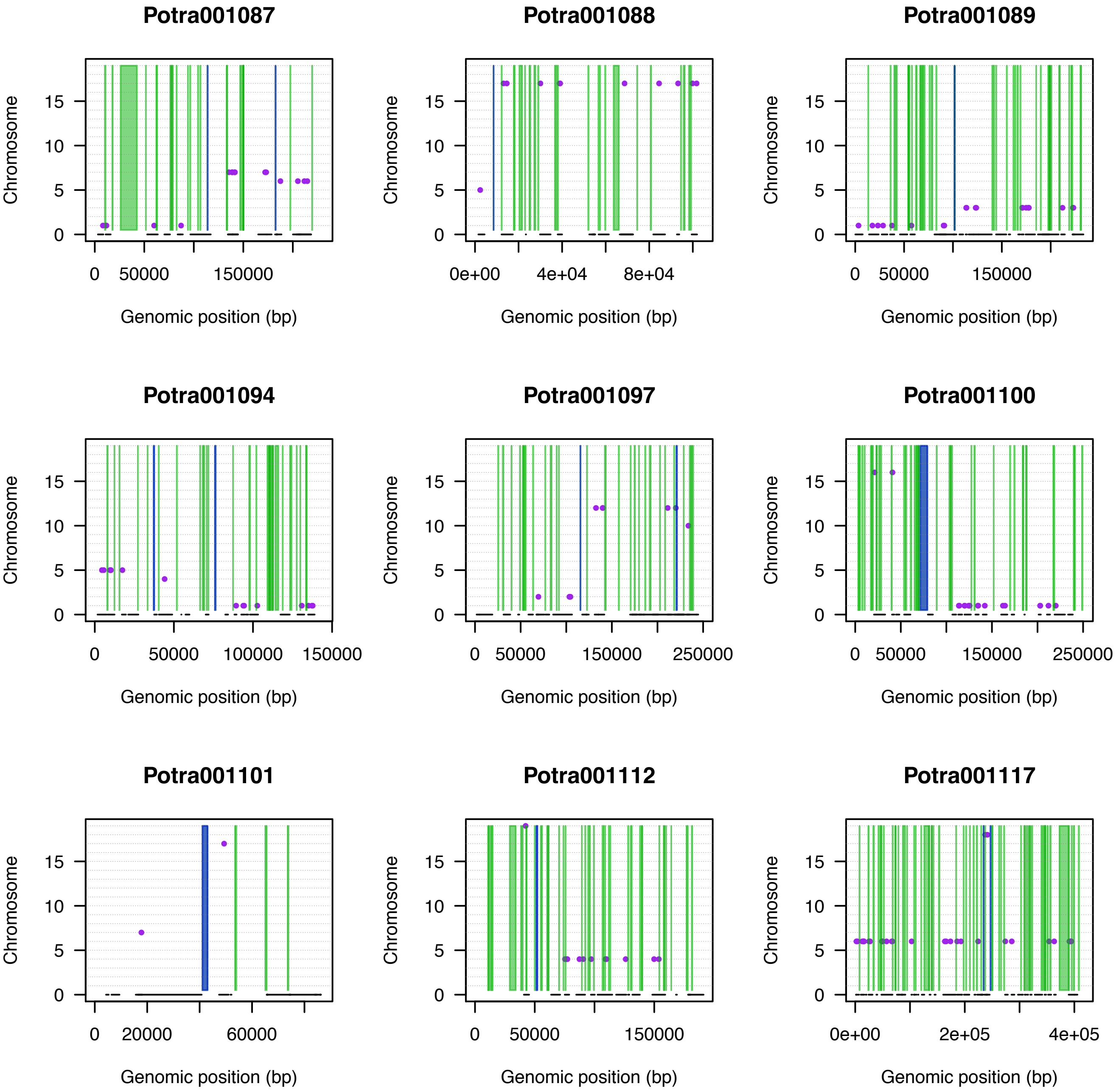
Potra001087
Potra001088
Potra001089
Potra001094
Potra001097
Potra001100
Potra001101
Potra001112
Potra001117
Chromosome
Genomic position (bp)

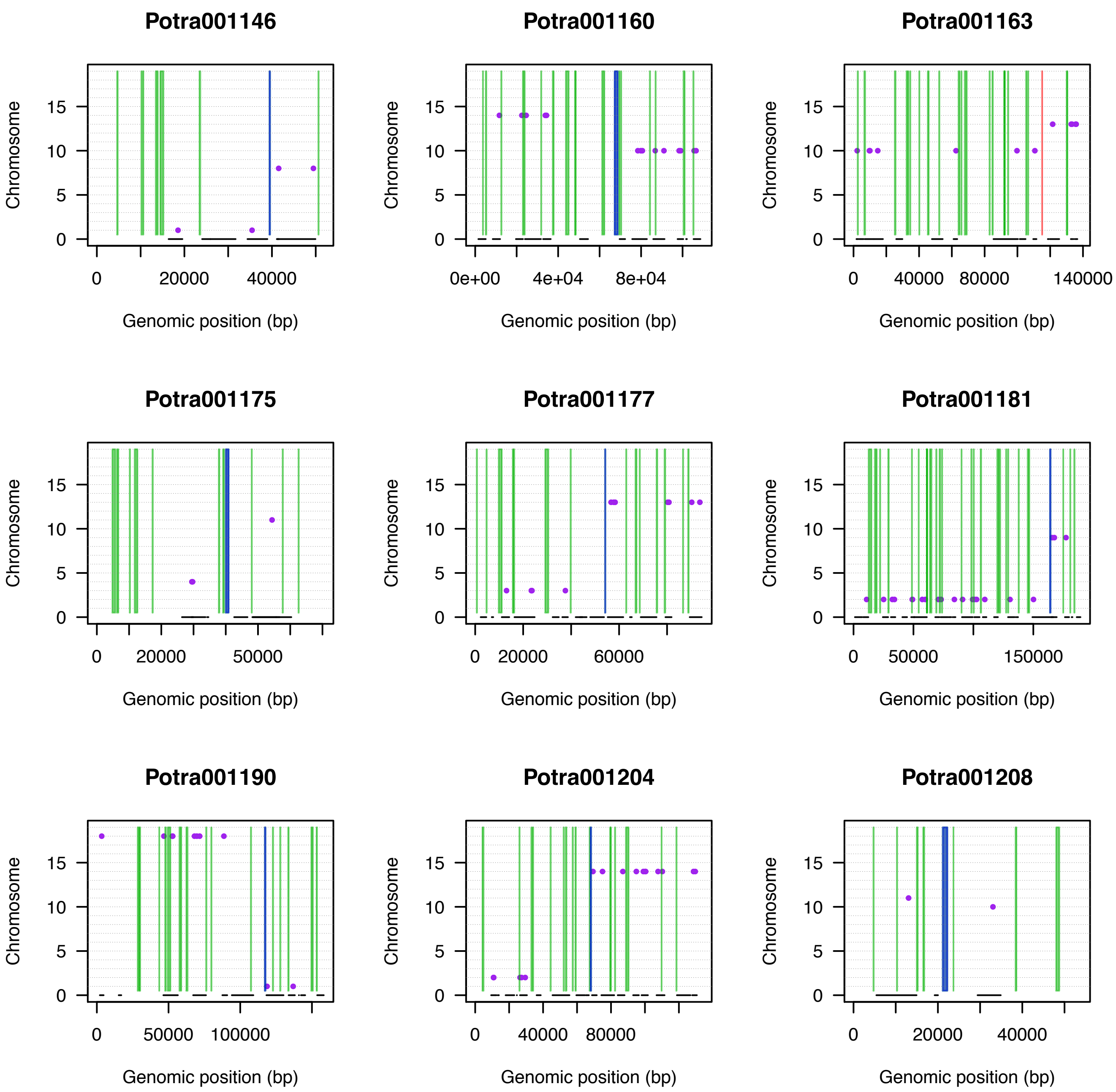

Potra001146
Potra001160
Potra001163
Potra001175
Potra001177
Potra001181
Potra001190
Potra001204
Potra001208
Chromosome
0
5
10
15
Genomic position (bp)
0
20000
40000
0e+00
4e+04
8e+04
80000
140000
50000
60000
150000
100000

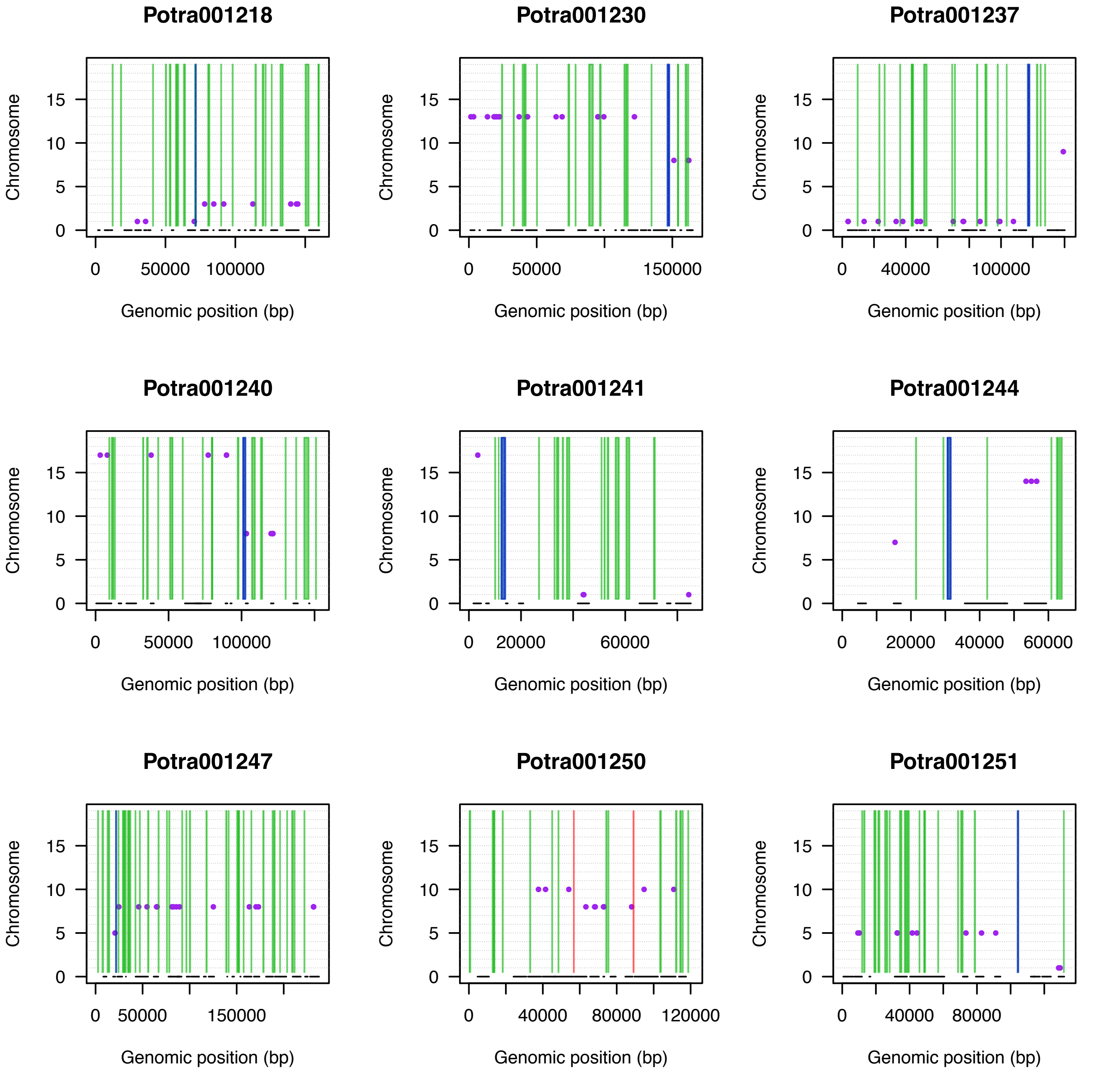

Potra001218
Potra001230
Potra001237
Potra001240
Potra001241
Potra001244
Potra001247
Potra001250
Potra001251
Chromosome
0
5
10
15
Genomic position (bp)
0
50000
100000
150000
0
40000
100000
0
50000
100000
0
20000
60000
0
20000
40000
60000
0
50000
150000
0
40000
80000
120000
0
40000
80000

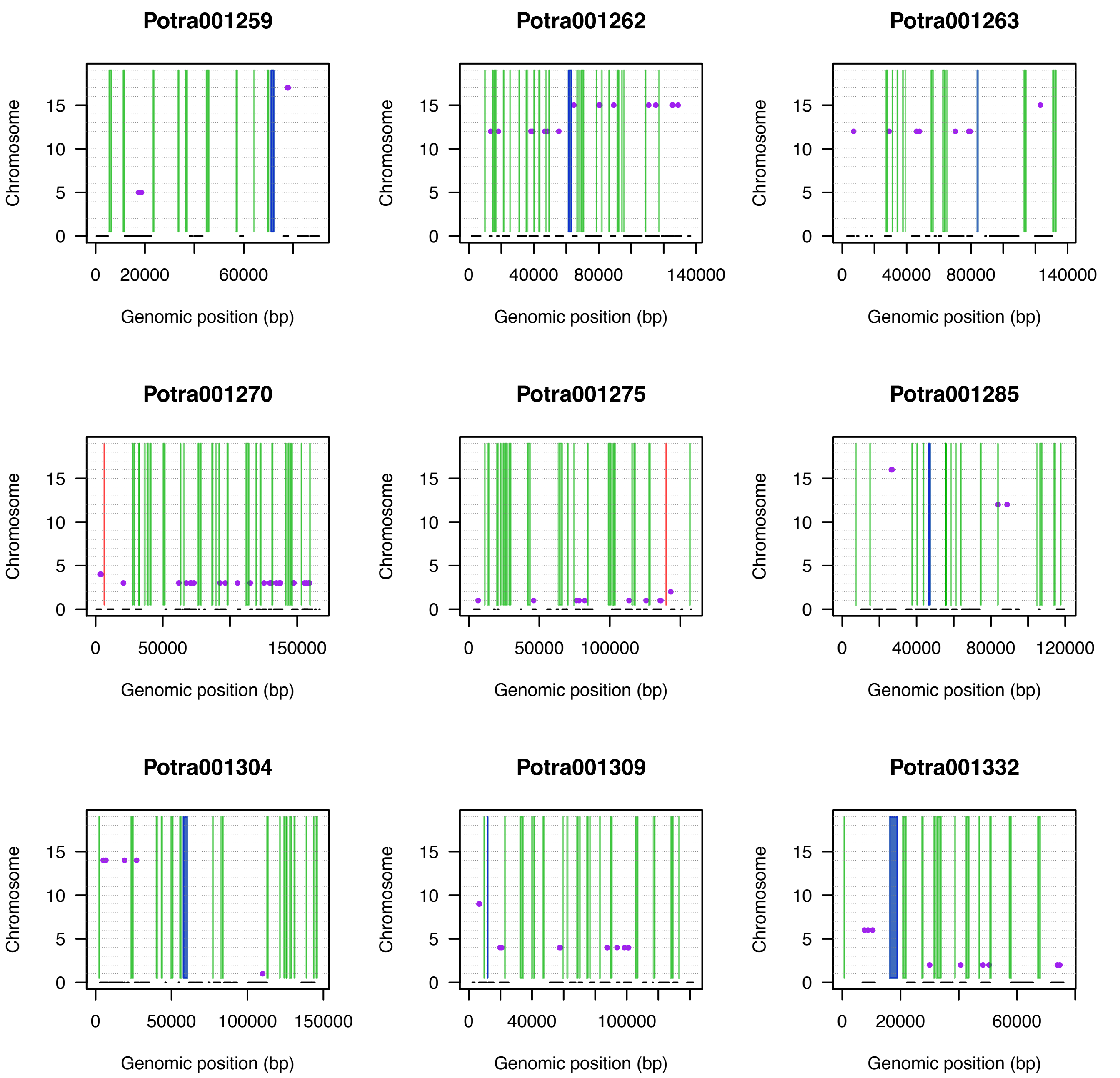
Potra001259
Potra001262
Potra001263
Potra001270
Potra001275
Potra001285
Potra001304
Potra001309
Potra001332
Chromosome
Genomic position (bp)

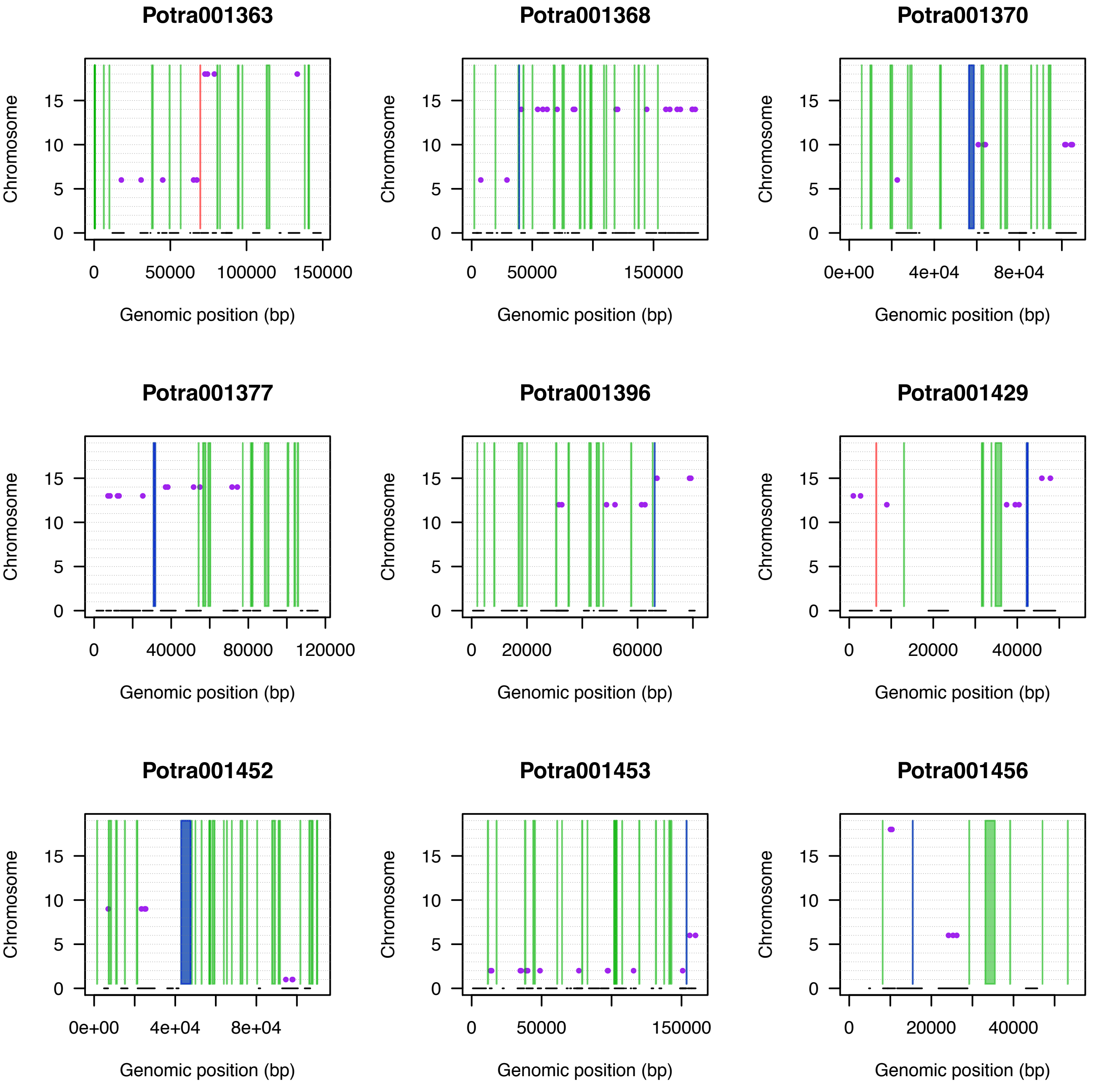
Potra001363
Potra001368
Potra001370
Potra001377
Potra001396
Potra001429
Potra001452
Potra001453
Potra001456
Chromosome
0
5
10
15
Genomic position (bp)
0
50000
100000
150000
0e+00
4e+04
8e+04
40000
80000
120000
20000
60000

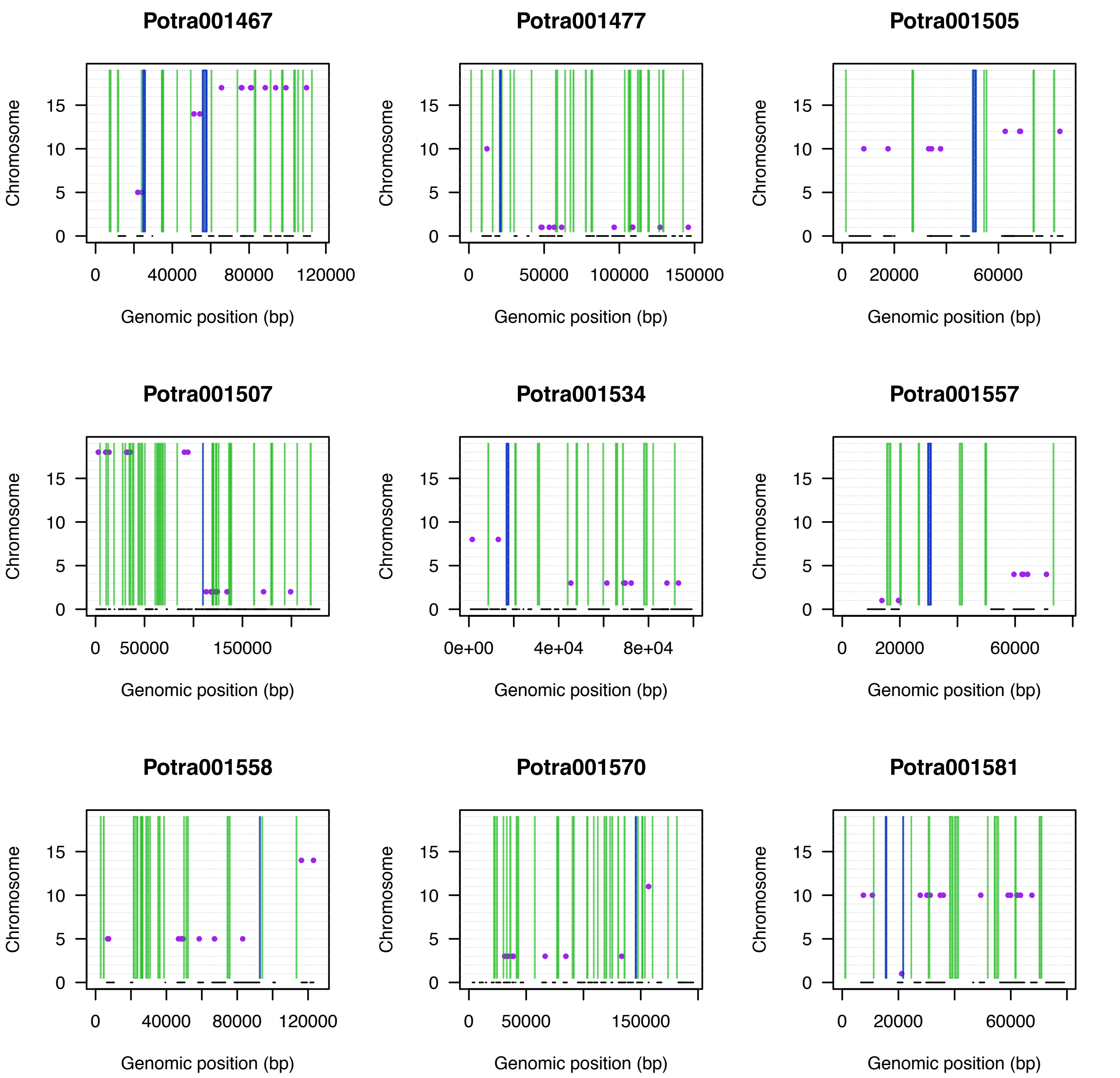
Potra001467
Chromosome
0
5
10
15
0
40000
80000
120000
Genomic position (bp)
Potra001477
Chromosome
0
5
10
15
0
50000
100000
150000
Genomic position (bp)
Potra001505
Chromosome
0
5
10
15
0
20000
60000
Genomic position (bp)
Potra001507
Chromosome
0
5
10
15
0
50000
150000
Genomic position (bp)
Potra001534
Chromosome
0
5
10
15
0e+00
4e+04
8e+04
Genomic position (bp)
Potra001557
Chromosome
0
5
10
15
0
20000
60000
Genomic position (bp)
Potra001558
Chromosome
0
5
10
15
0
40000
80000
120000
Genomic position (bp)
Potra001570
Chromosome
0
5
10
15
0
50000
150000
Genomic position (bp)
Potra001581
Chromosome
0
5
10
15
0
20000
60000
Genomic position (bp)

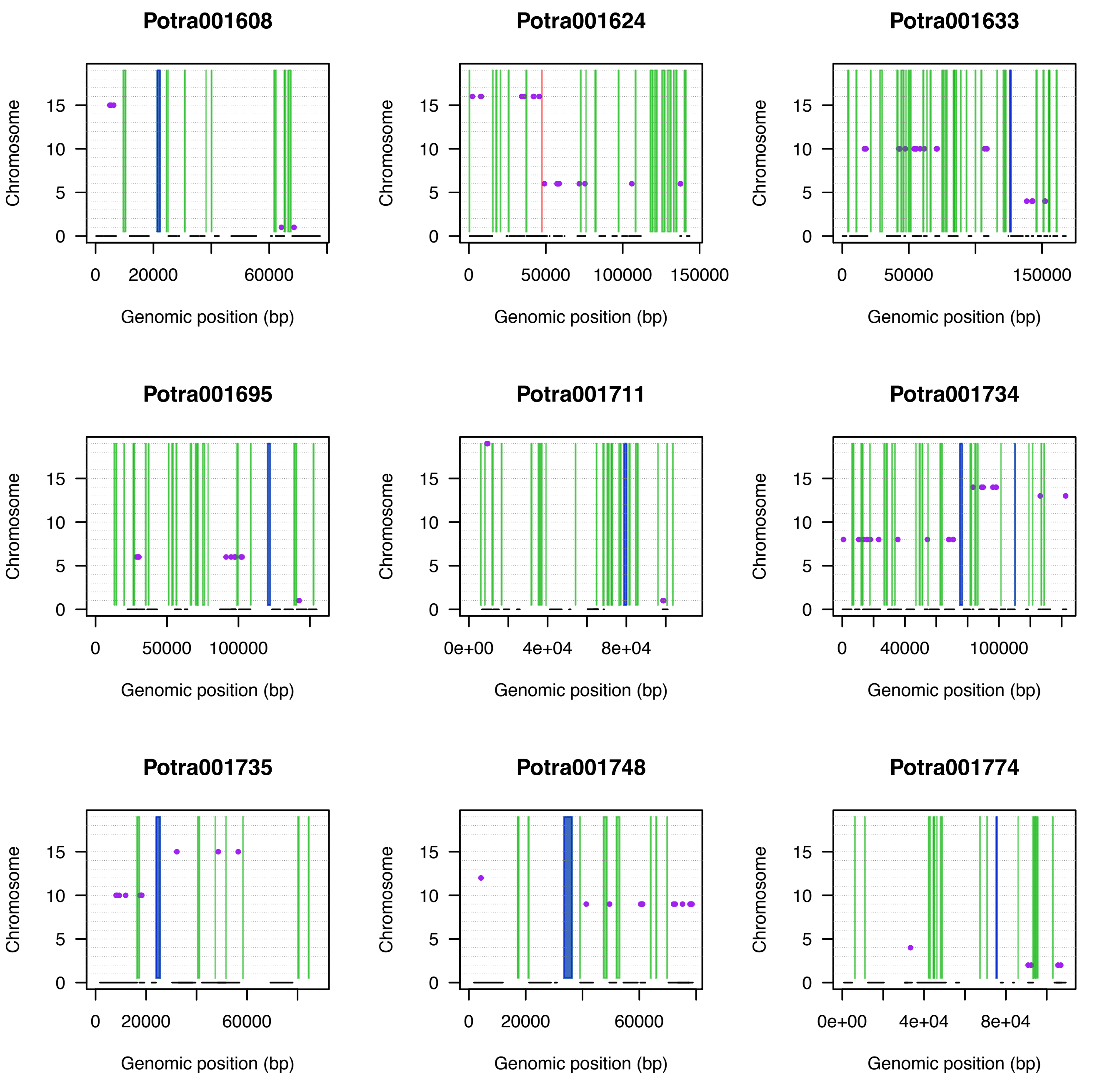
Potra001608
Potra001624
Potra001633
Potra001695
Potra001711
Potra001734
Potra001735
Potra001748
Potra001774
Chromosome
Genomic position (bp)
0
5
10
15
0
20000
60000
50000
100000
150000
0e+00
4e+04
8e+04
40000

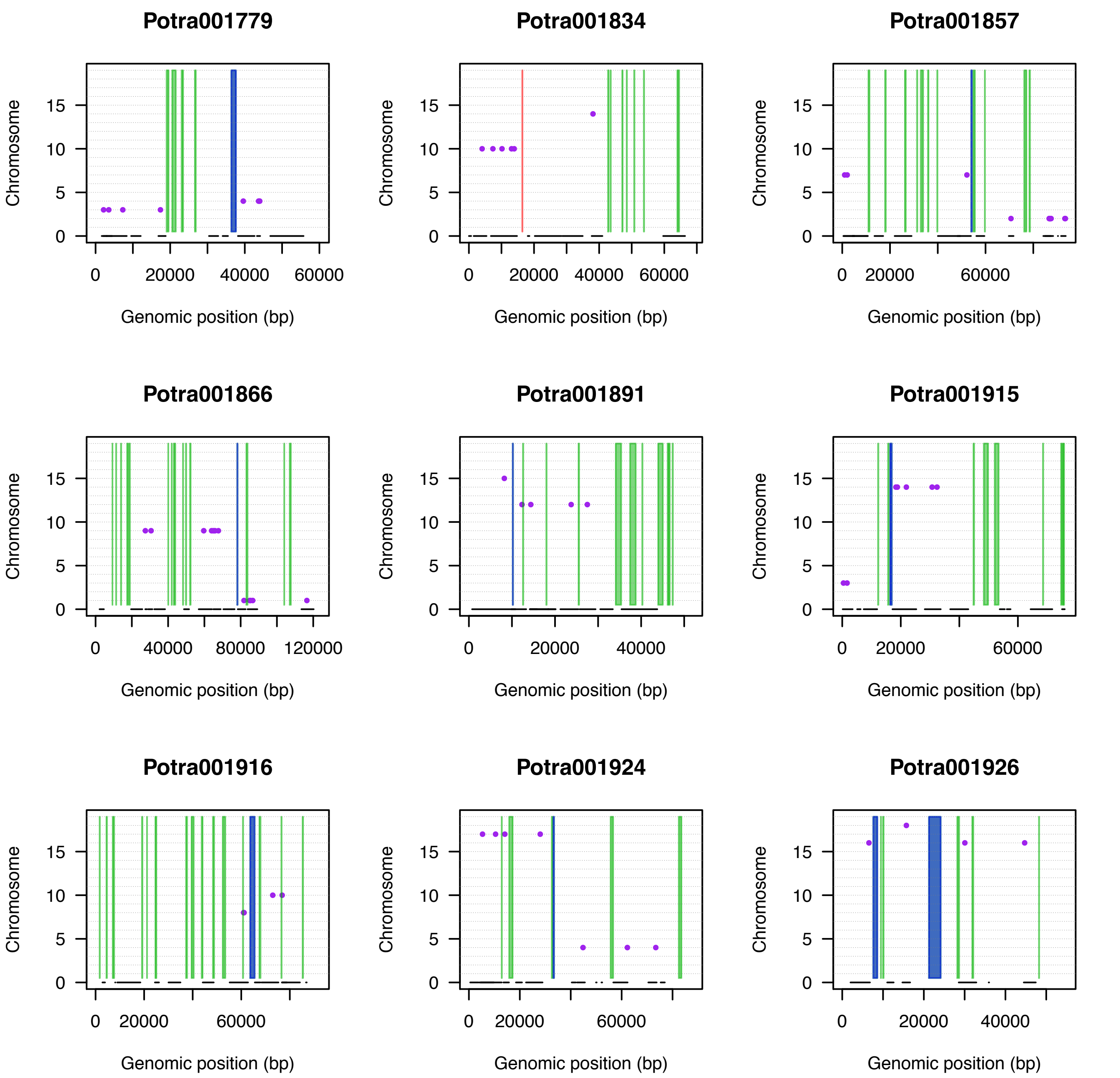
Potra001779
Potra001834
Potra001857
Potra001866
Potra001891
Potra001915
Potra001916
Potra001924
Potra001926
Chromosome
0
5
10
15
0
20000
40000
60000
80000
120000
Genomic position (bp)

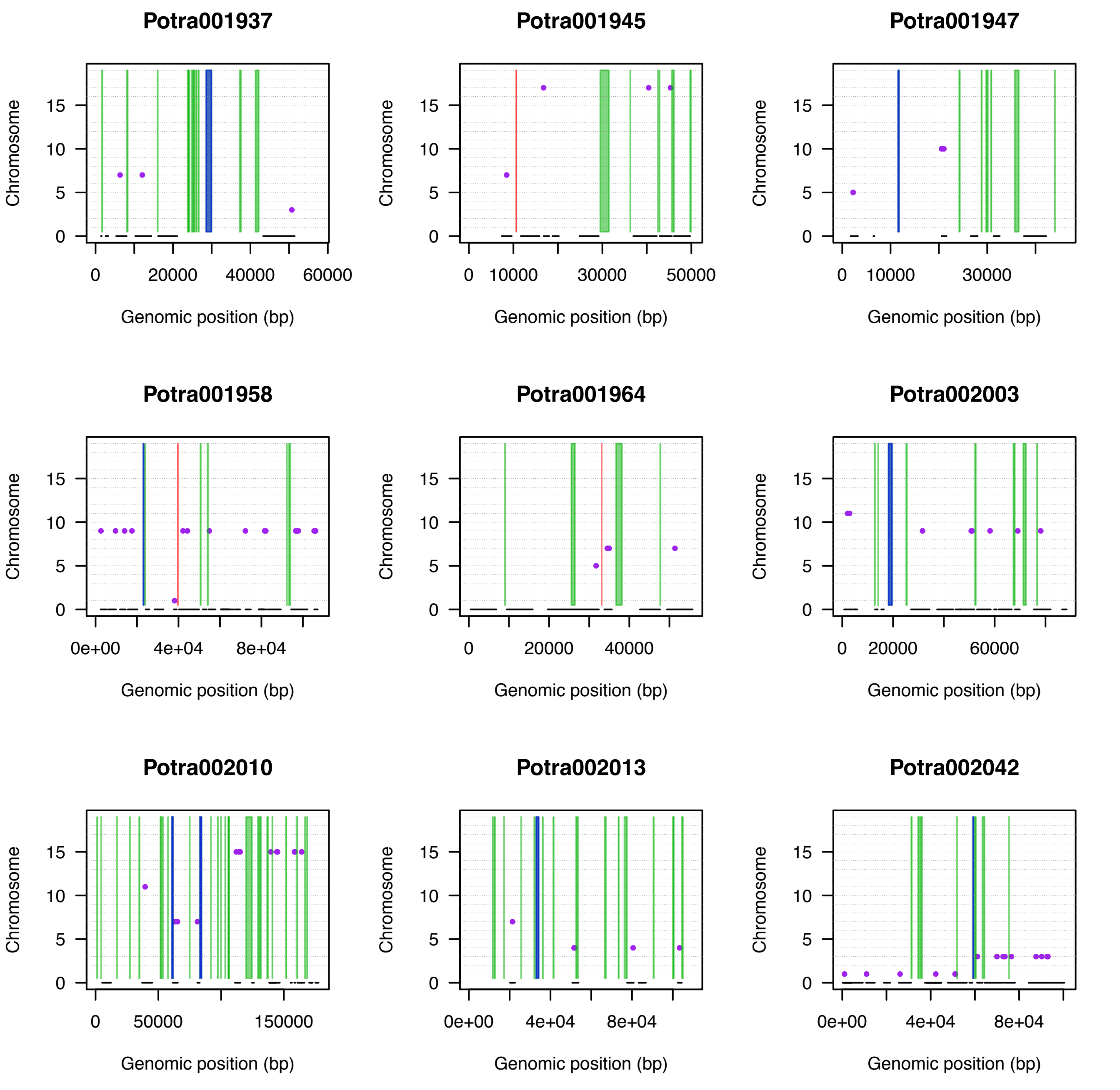

Potra001937
Potra001945
Potra001947
Potra001958
Potra001964
Potra002003
Potra002010
Potra002013
Potra002042
Chromosome
Genomic position (bp)

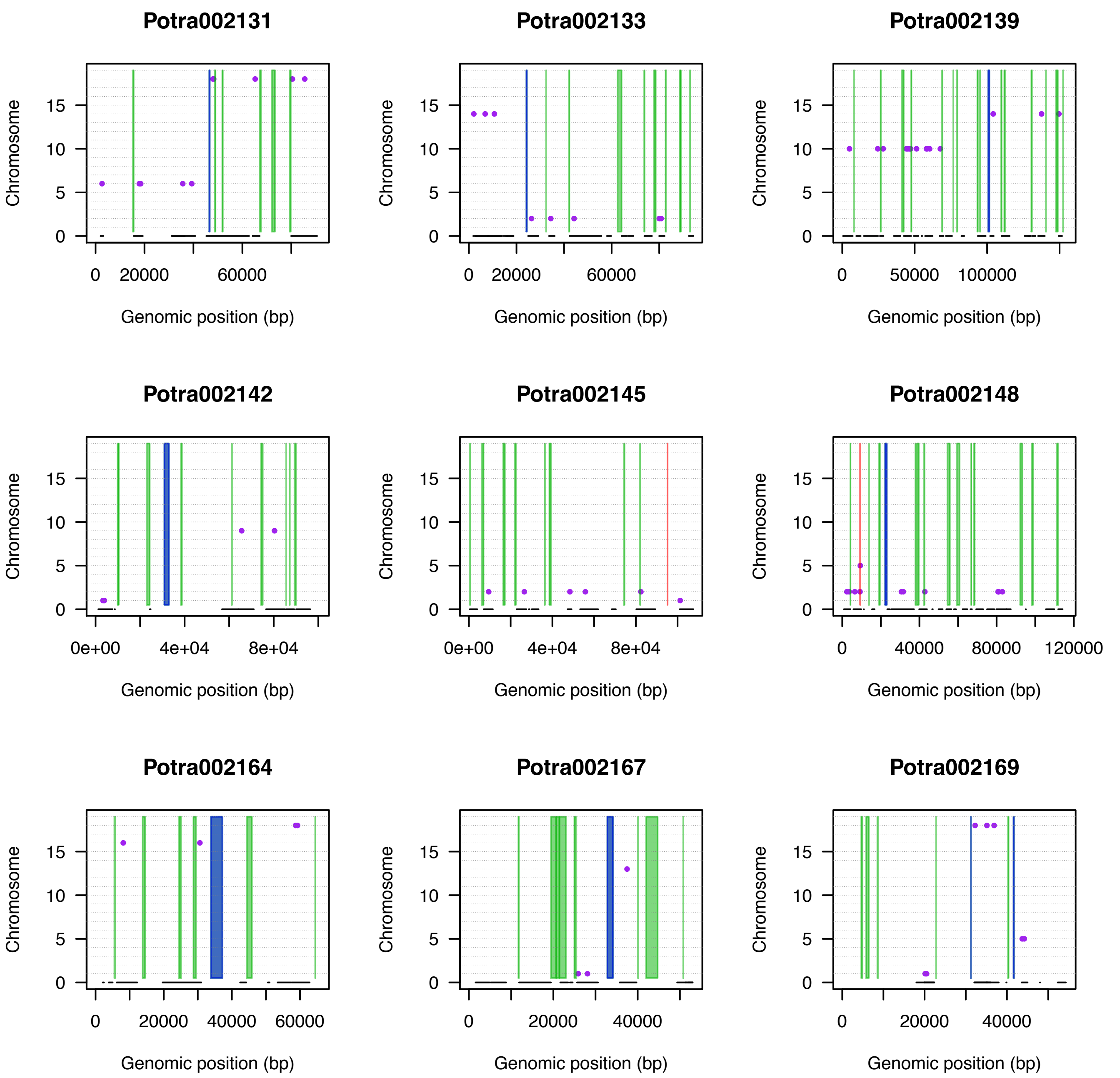
Potra002131
Potra002133
Potra002139
Potra002142
Potra002145
Potra002148
Potra002164
Potra002167
Potra002169
Chromosome
Genomic position (bp)
0
5
10
15
0
20000
60000
50000
100000
0e+00
4e+04
8e+04
40000
80000
120000
40000
60000

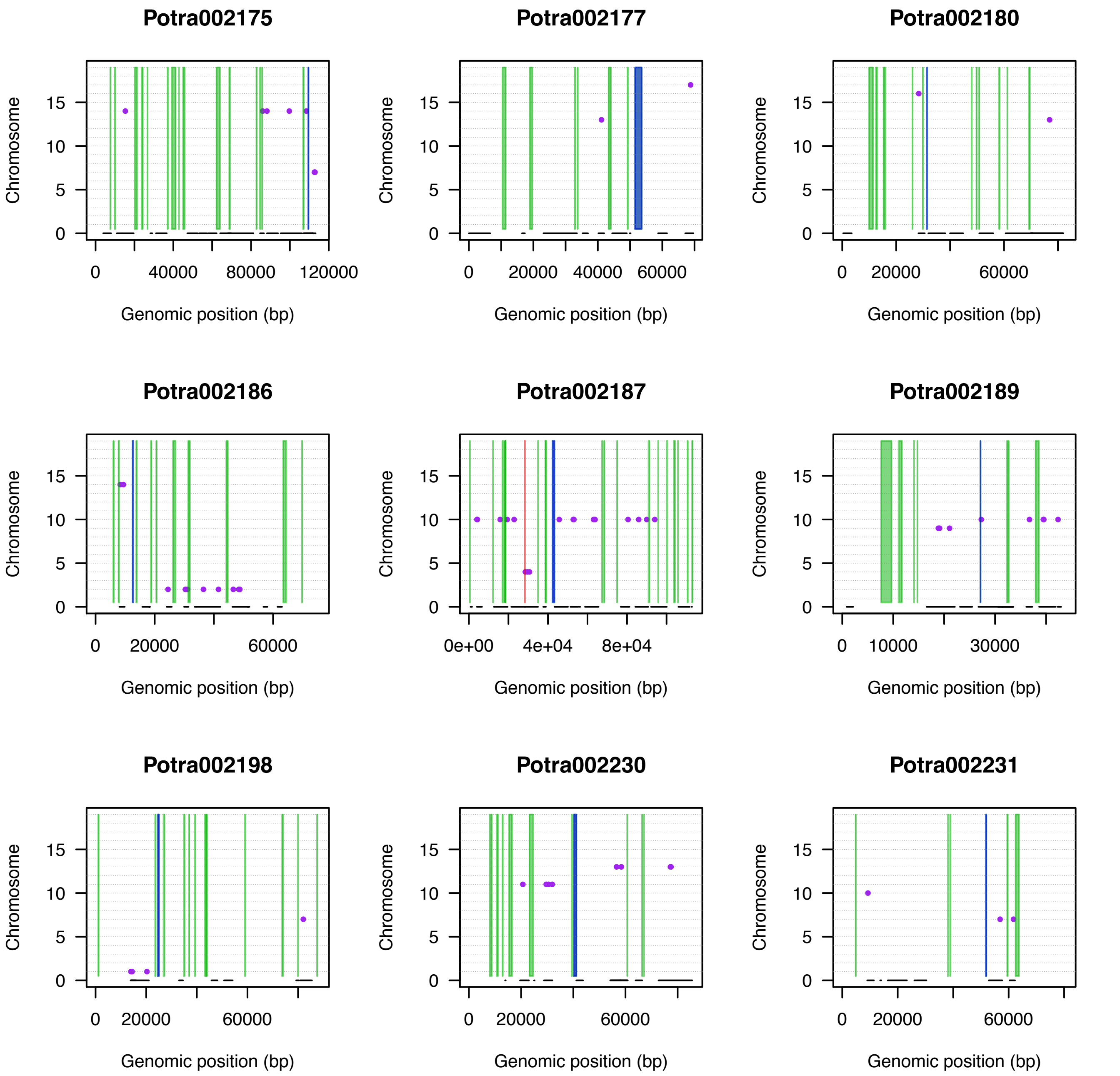

Potra002175
Potra002177
Potra002180
Potra002186
Potra002187
Potra002189
Potra002198
Potra002230
Potra002231
Chromosome
0
5
10
15
Genomic position (bp)
0
40000
80000
120000
20000
60000
10000
30000
0e+00
4e+04
8e+04

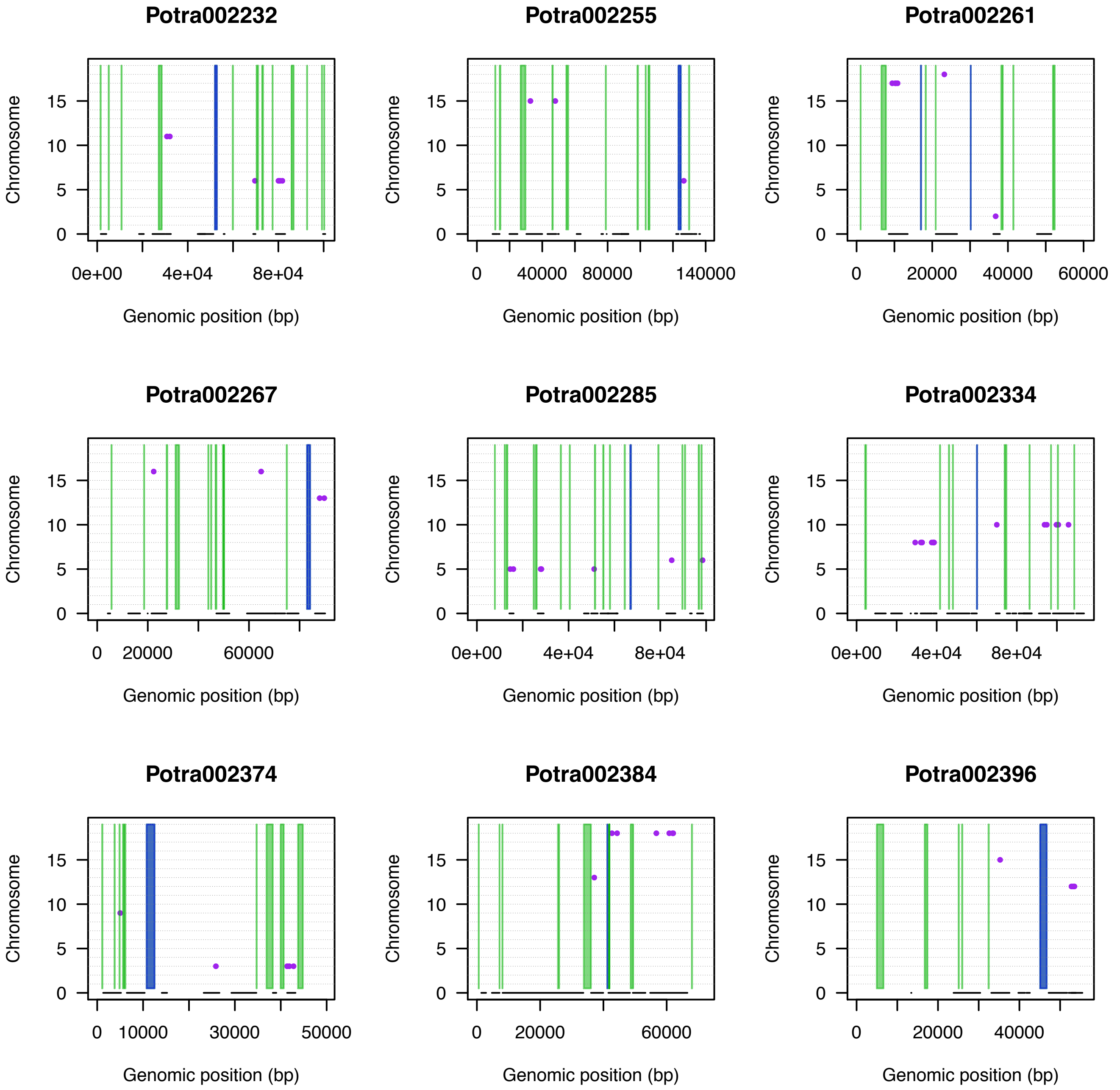
Potra002232
Potra002255
Potra002261
Potra002267
Potra002285
Potra002334
Potra002374
Potra002384
Potra002396
Chromosome
0
5
10
15
Genomic position (bp)
0e+00
4e+04
8e+04
0
40000
80000
140000
0
20000
40000
60000
0
20000
60000
0
10000
30000
50000

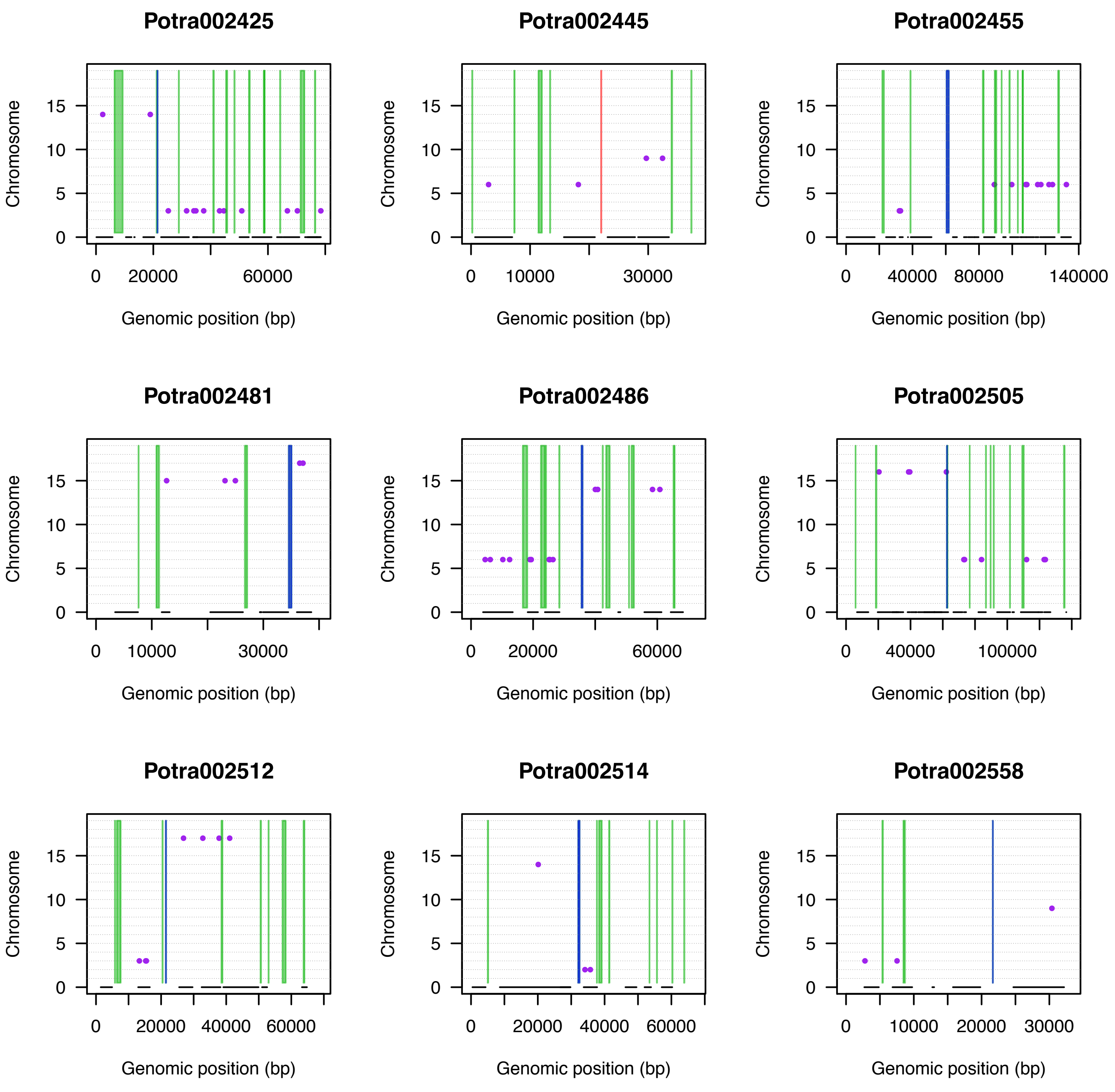

Potra002425
Potra002445
Potra002455
Potra002481
Potra002486
Potra002505
Potra002512
Potra002514
Potra002558
Chromosome
Genomic position (bp)

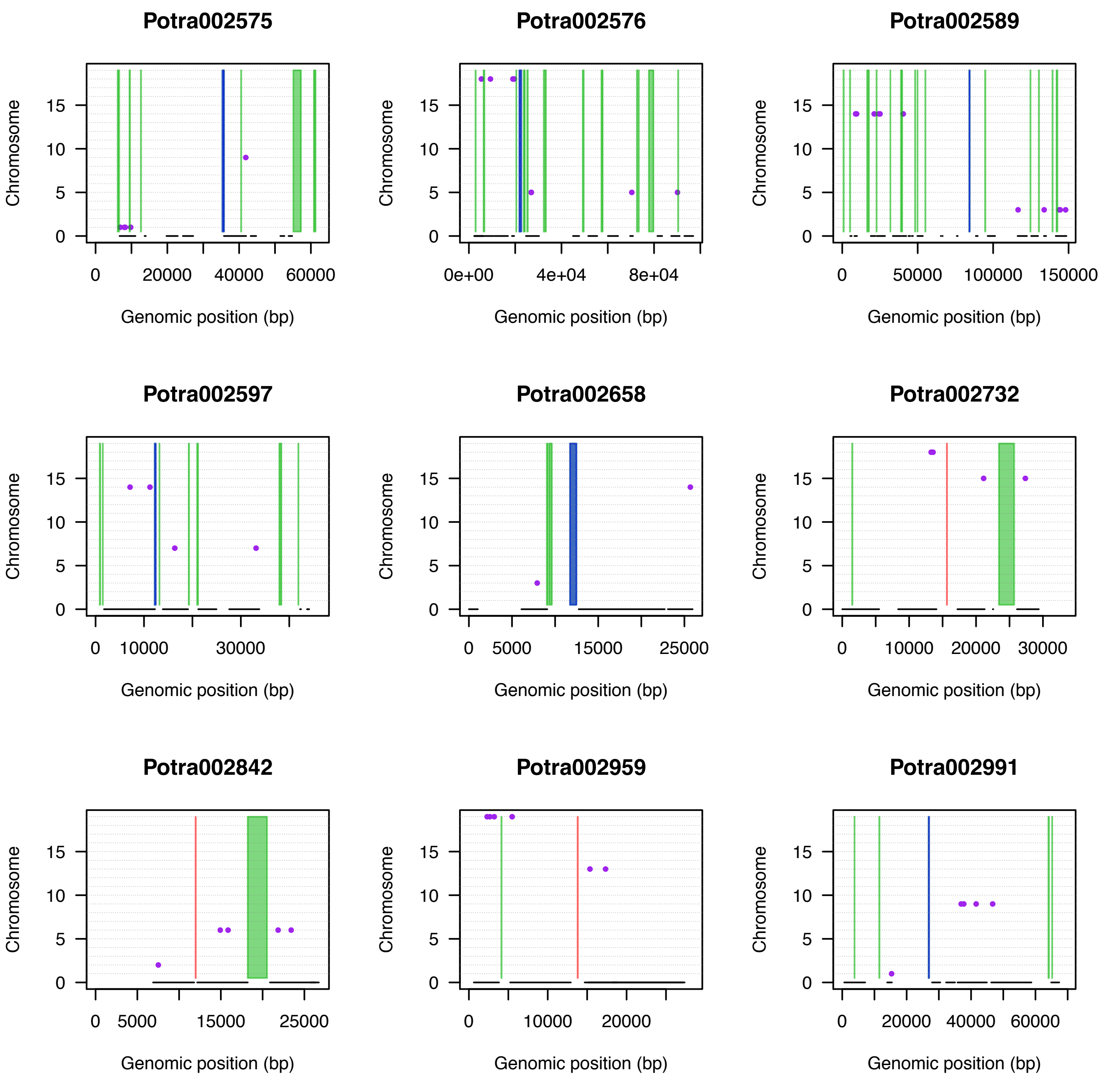
Potra002575
Potra002576
Potra002589
Potra002597
Potra002658
Potra002732
Potra002842
Potra002959
Potra002991
Chromosome
Genomic position (bp)
0
5
10
15
0
20000
40000
60000
0e+00
4e+04
8e+04
50000
100000
150000
10000
30000
5000
15000
25000
20000

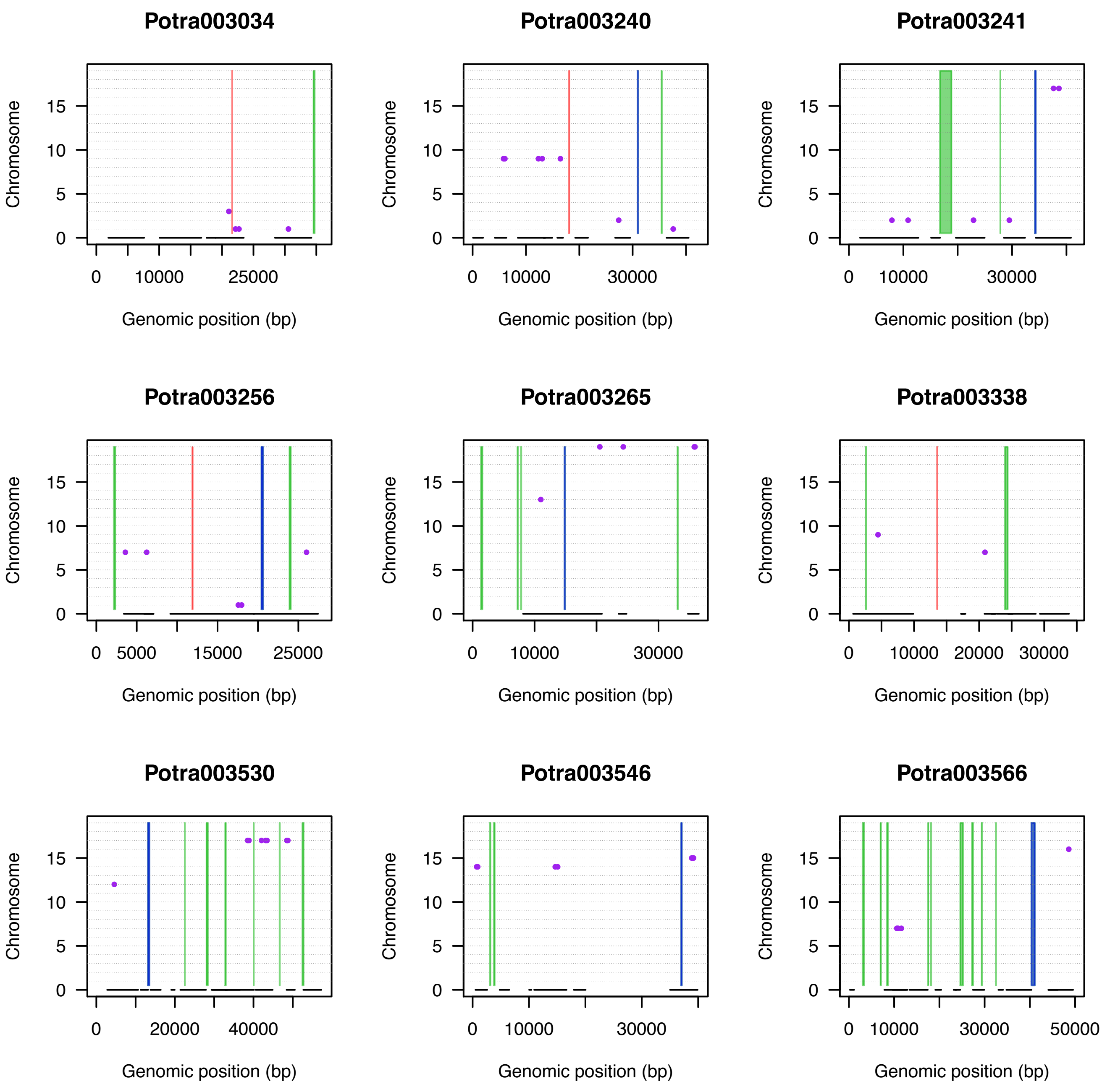
Potra003034
Potra003240
Potra003241
Potra003256
Potra003265
Potra003338
Potra003530
Potra003546
Potra003566
Chromosome
Genomic position (bp)
0
5
10
15
0
10000
25000
0
10000
30000
0
5000
15000
25000
0
10000
20000
30000
0
20000
40000
0
10000
30000
50000

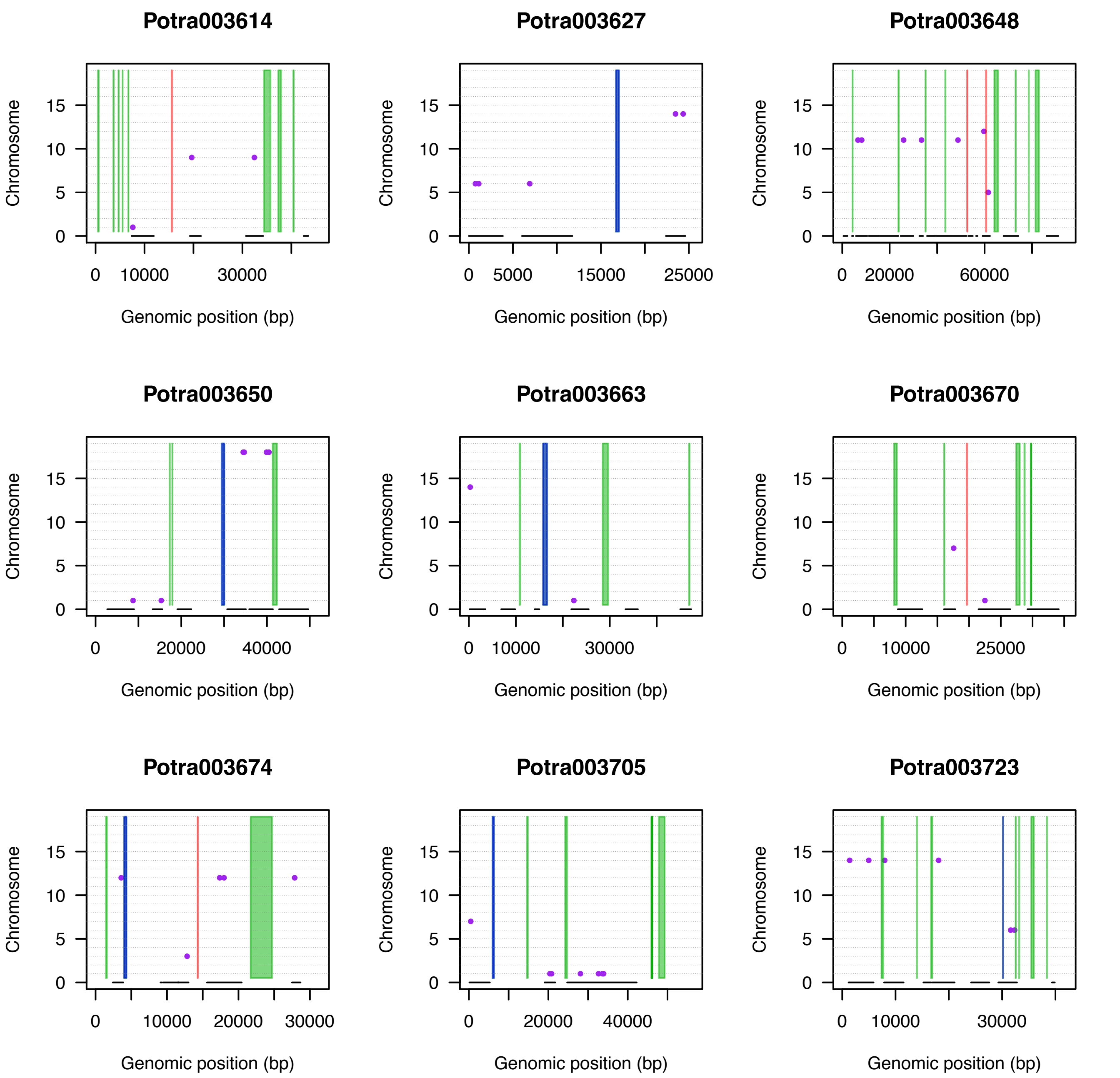

Potra003614
Potra003627
Potra003648
Potra003650
Potra003663
Potra003670
Potra003674
Potra003705
Potra003723
Chromosome
Genomic position (bp)
0
5
10
15
0
10000
30000
5000
15000
25000
20000
60000
40000

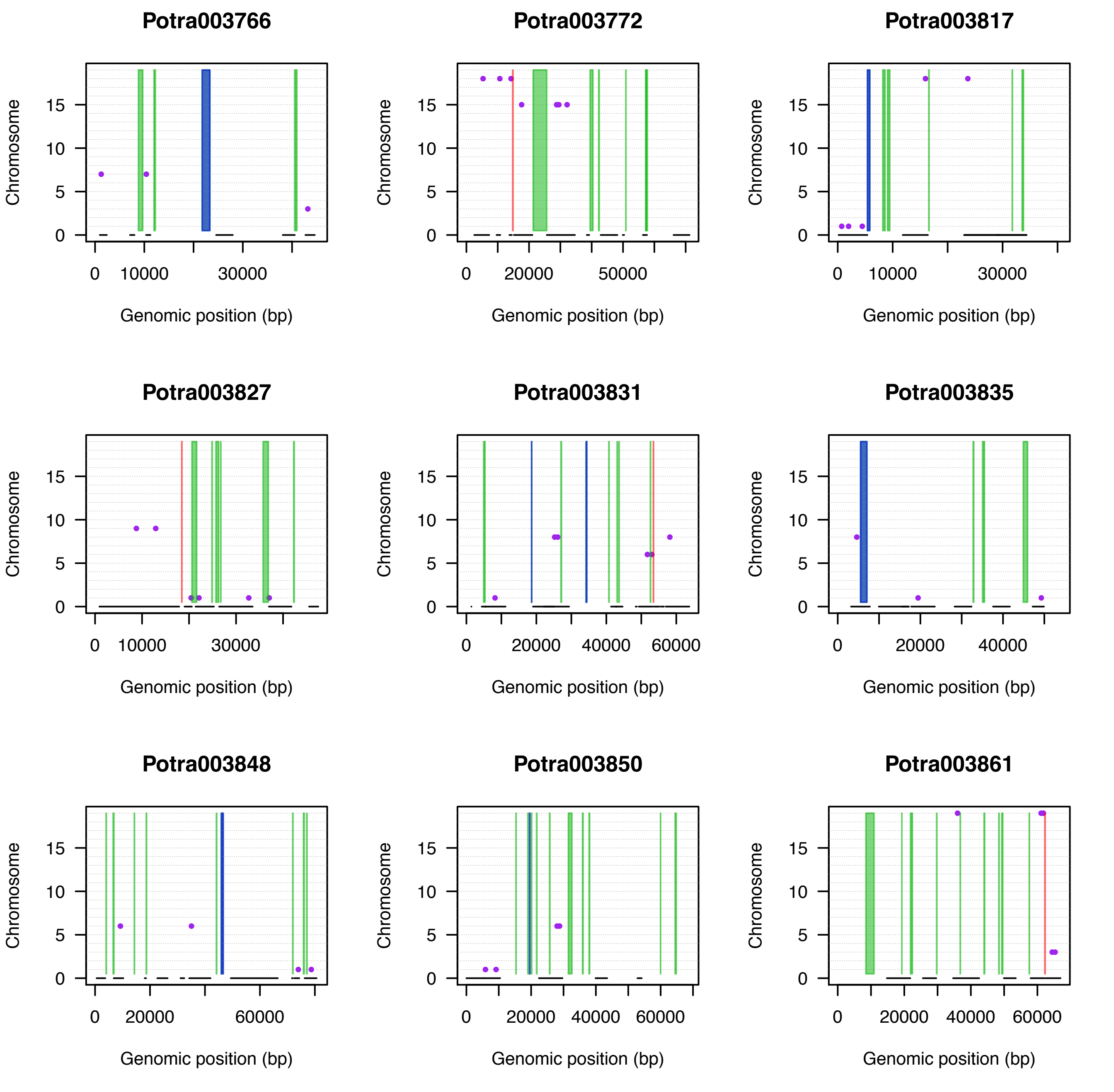
Potra003766
Potra003772
Potra003817
Potra003827
Potra003831
Potra003835
Potra003848
Potra003850
Potra003861
Chromosome
0
5
10
15
Genomic position (bp)
0
10000
20000
30000
40000
50000
60000

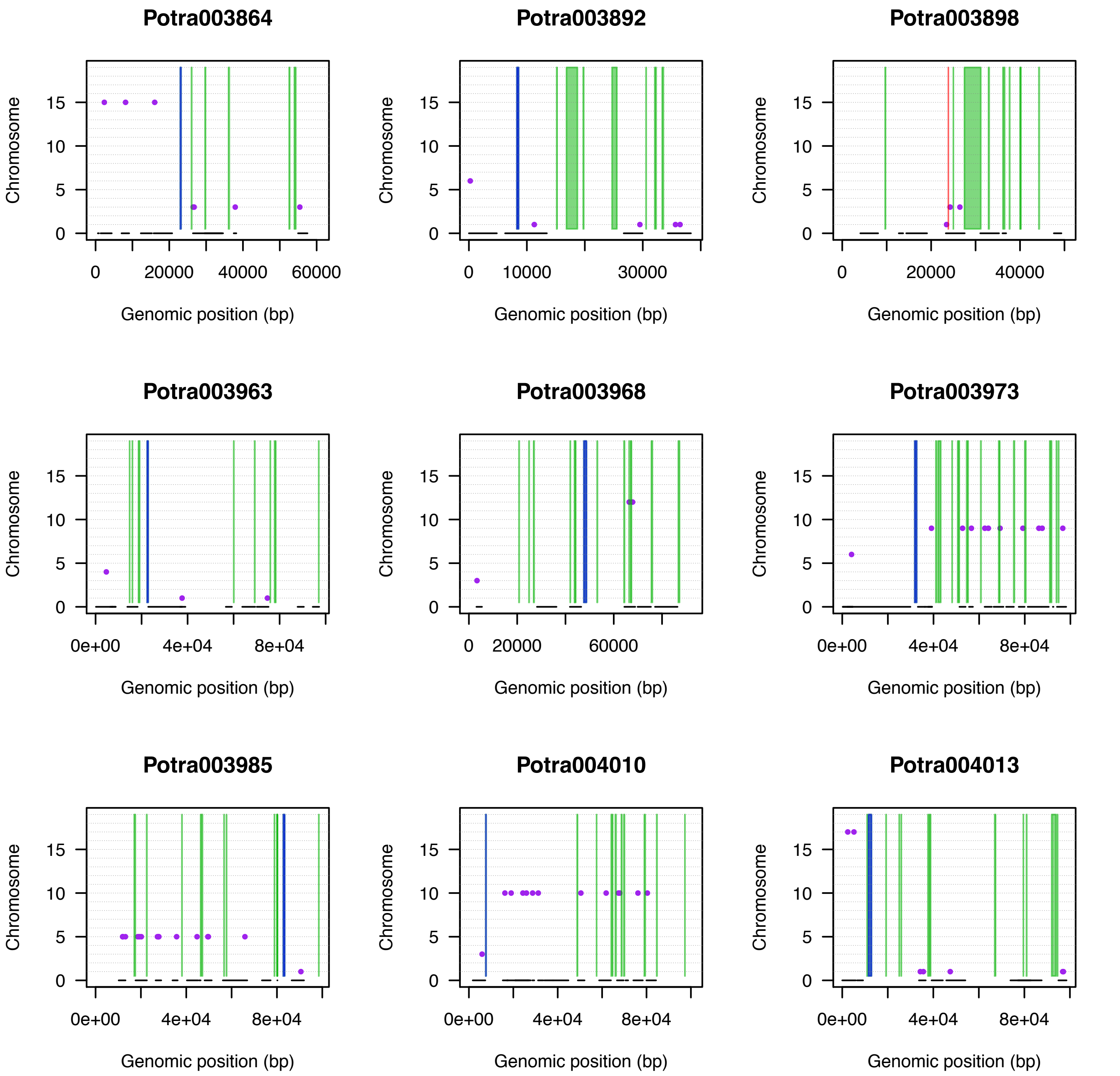
Potra003864
Potra003892
Potra003898
Potra003963
Potra003968
Potra003973
Potra003985
Potra004010
Potra004013
Chromosome
0
5
10
15
Genomic position (bp)
0
20000
40000
60000
10000
30000
0e+00
4e+04
8e+04

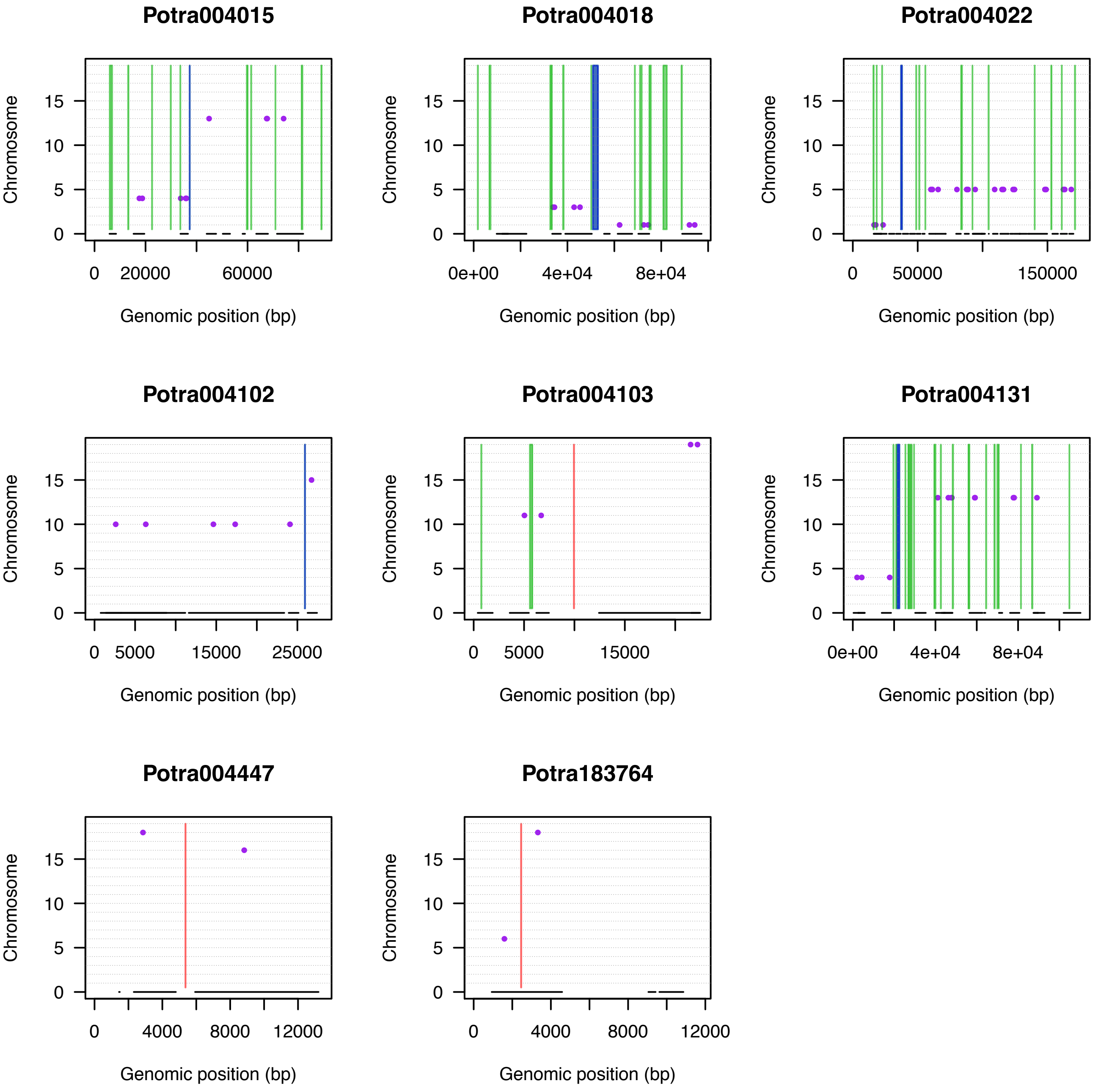
Potra004015
Potra004018
Potra004022
Potra004102
Potra004103
Potra004131
Potra004447
Potra183764
Chromosome
Genomic position (bp)
0
5
10
15
0
20000
60000
0e+00
4e+04
8e+04
50000
150000
5000
15000
25000
4000
8000
12000

Supplementary Figure 5

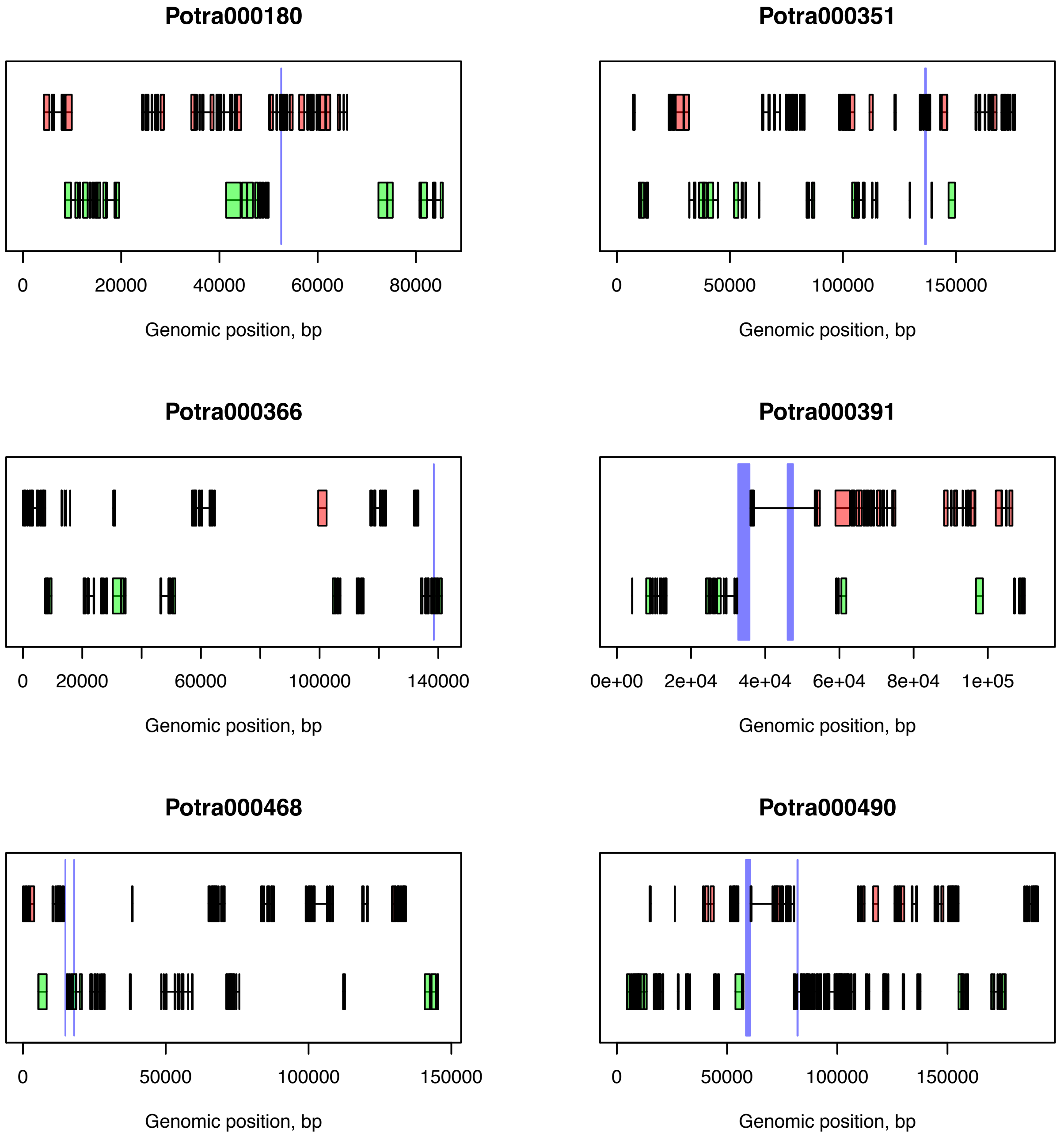

Potra000180
0
20000
40000
60000
80000
Genomic position, bp
Potra000351
0
50000
100000
150000
Genomic position, bp
Potra000366
0
20000
60000
100000
140000
Genomic position, bp
Potra000391
0e+00
2e+04
4e+04
6e+04
8e+04
1e+05
Genomic position, bp
Potra000468
0
50000
100000
150000
Genomic position, bp
Potra000490
0
50000
100000
150000
Genomic position, bp

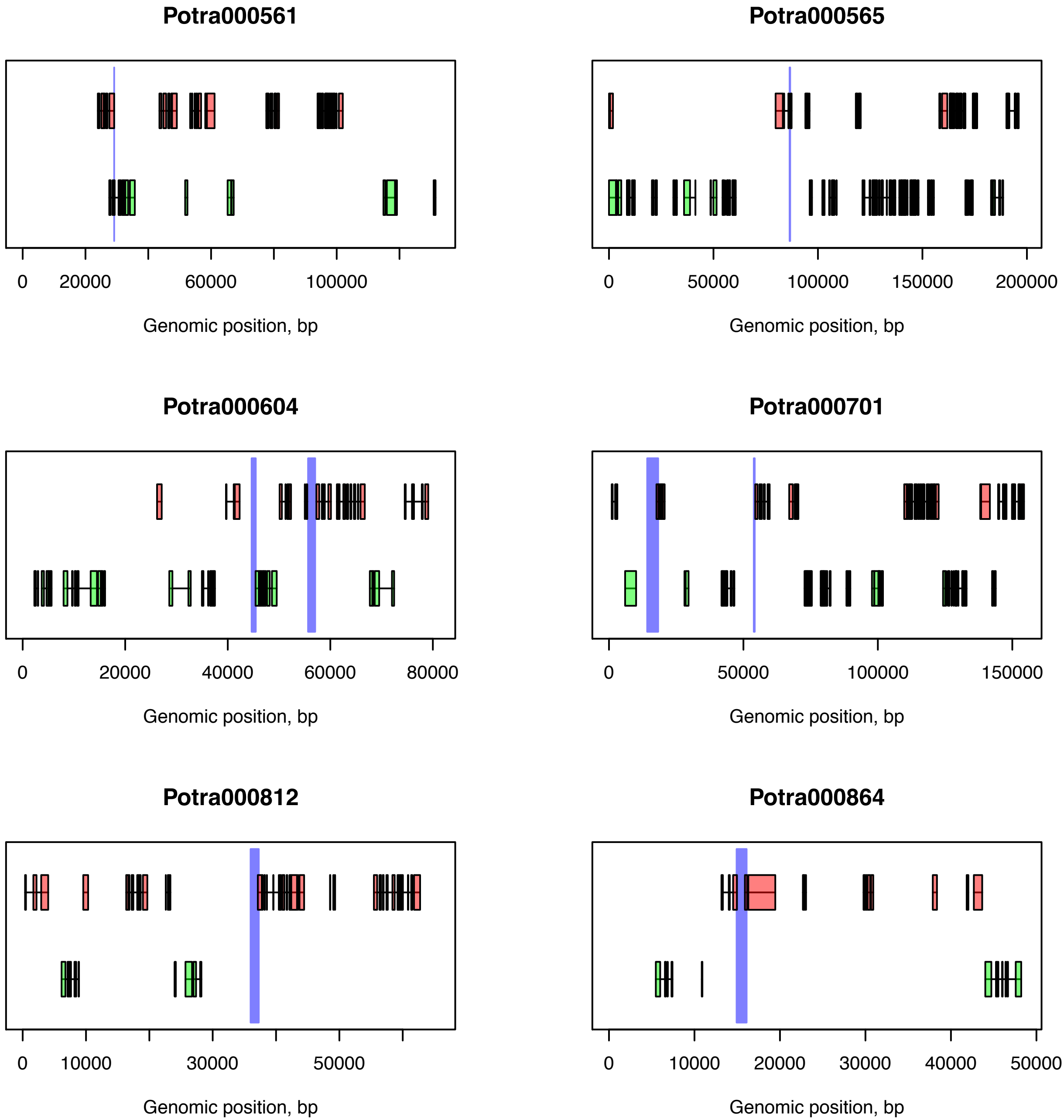
Potra000561
0
20000
60000
100000
Genomic position, bp
Potra000565
0
50000
100000
150000
200000
Genomic position, bp
Potra000604
0
20000
40000
60000
80000
Genomic position, bp
Potra000701
0
50000
100000
150000
Genomic position, bp
Potra000812
0
10000
30000
50000
Genomic position, bp
Potra000864
0
10000
20000
30000
40000
50000
Genomic position, bp

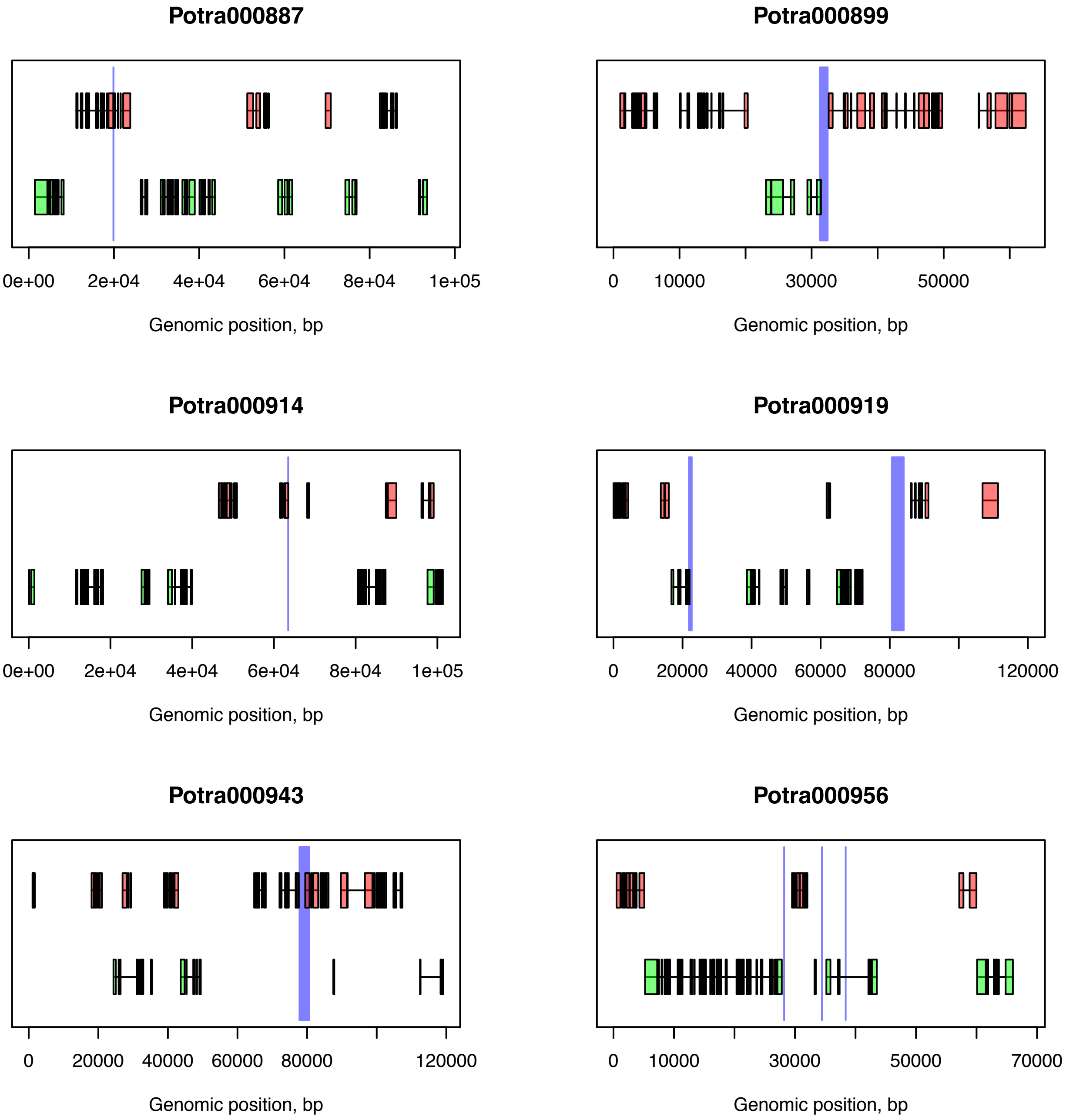

Potra000887
0e+00
2e+04
4e+04
6e+04
8e+04
1e+05
Genomic position, bp
Potra000899
0
10000
30000
50000
Genomic position, bp
Potra000914
0e+00
2e+04
4e+04
6e+04
8e+04
1e+05
Genomic position, bp
Potra000919
0
20000
40000
60000
80000
120000
Genomic position, bp
Potra000943
0
20000
40000
60000
80000
120000
Genomic position, bp
Potra000956
0
10000
30000
50000
70000
Genomic position, bp

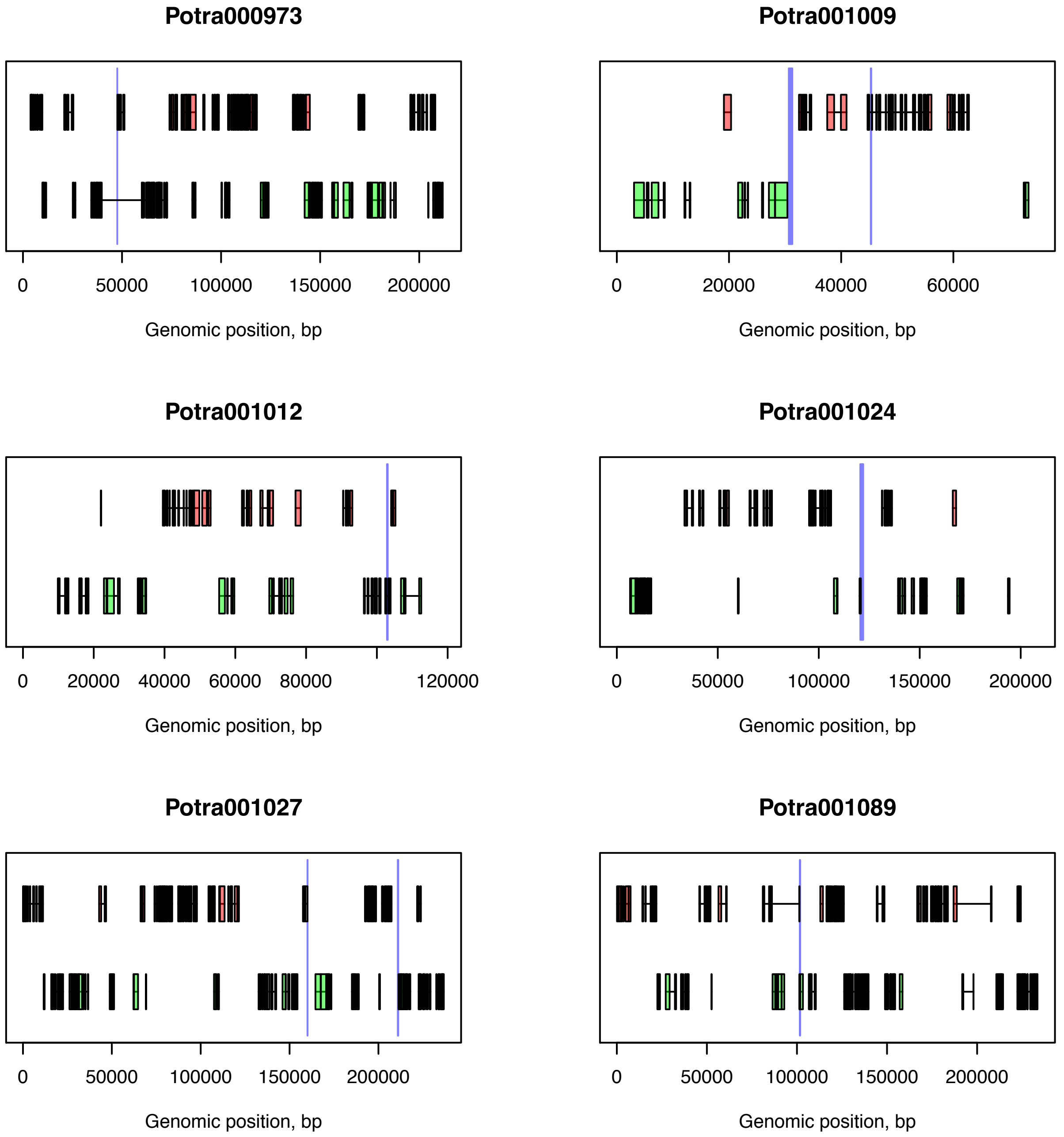
Potra000973
0
50000
100000
150000
200000
Genomic position, bp
Potra001009
0
20000
40000
60000
Genomic position, bp
Potra001012
0
20000
40000
60000
80000
120000
Genomic position, bp
Potra001024
0
50000
100000
150000
200000
Genomic position, bp
Potra001027
0
50000
100000
150000
200000
Genomic position, bp
Potra001089
0
50000
100000
150000
200000
Genomic position, bp

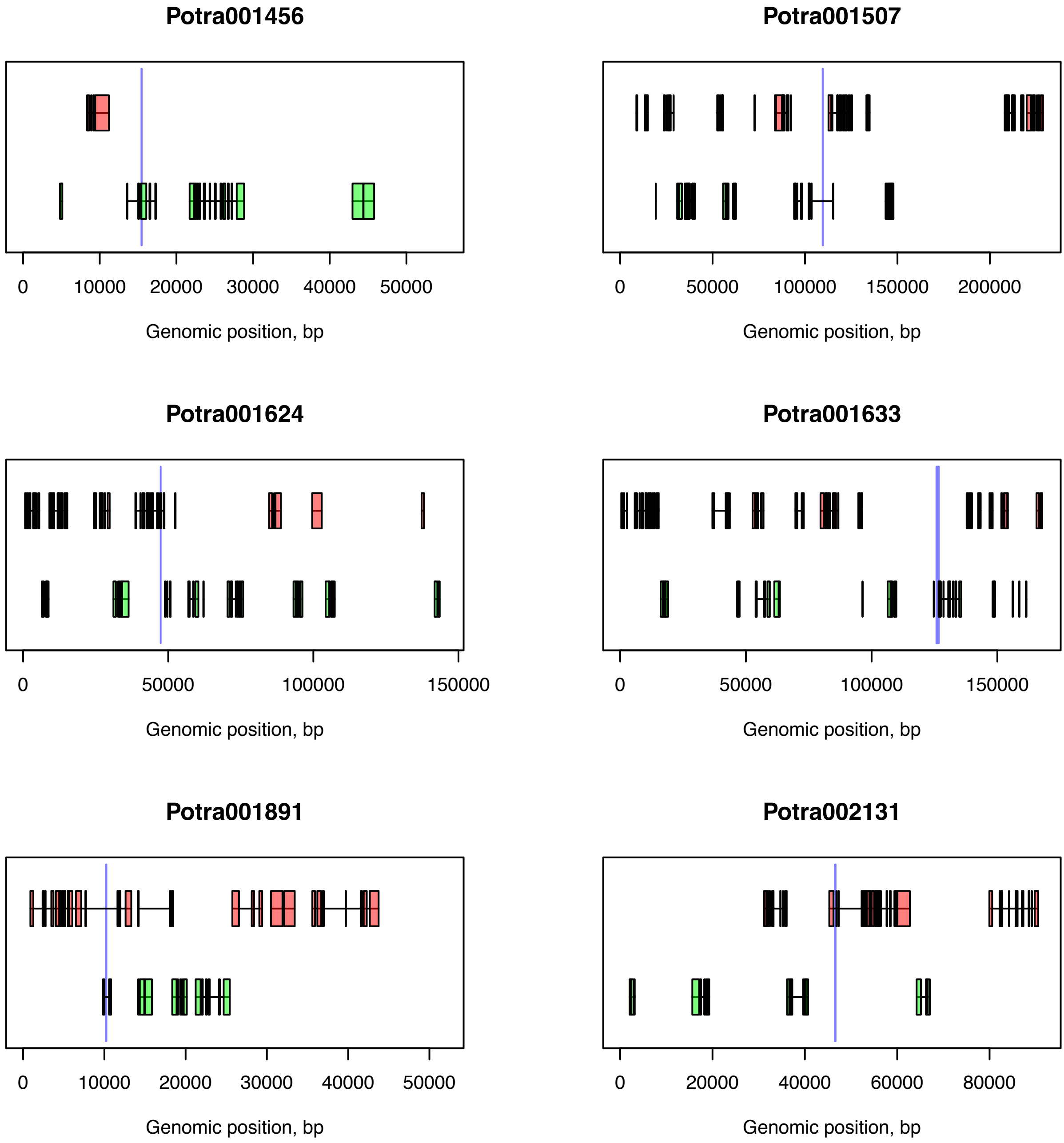

Potra001456
0
10000
20000
30000
40000
50000
Genomic position, bp
Potra001507
0
50000
100000
150000
200000
Genomic position, bp
Potra001624
0
50000
100000
150000
Genomic position, bp
Potra001633
0
50000
100000
150000
Genomic position, bp
Potra001891
0
10000
20000
30000
40000
50000
Genomic position, bp
Potra002131
0
20000
40000
60000
80000
Genomic position, bp

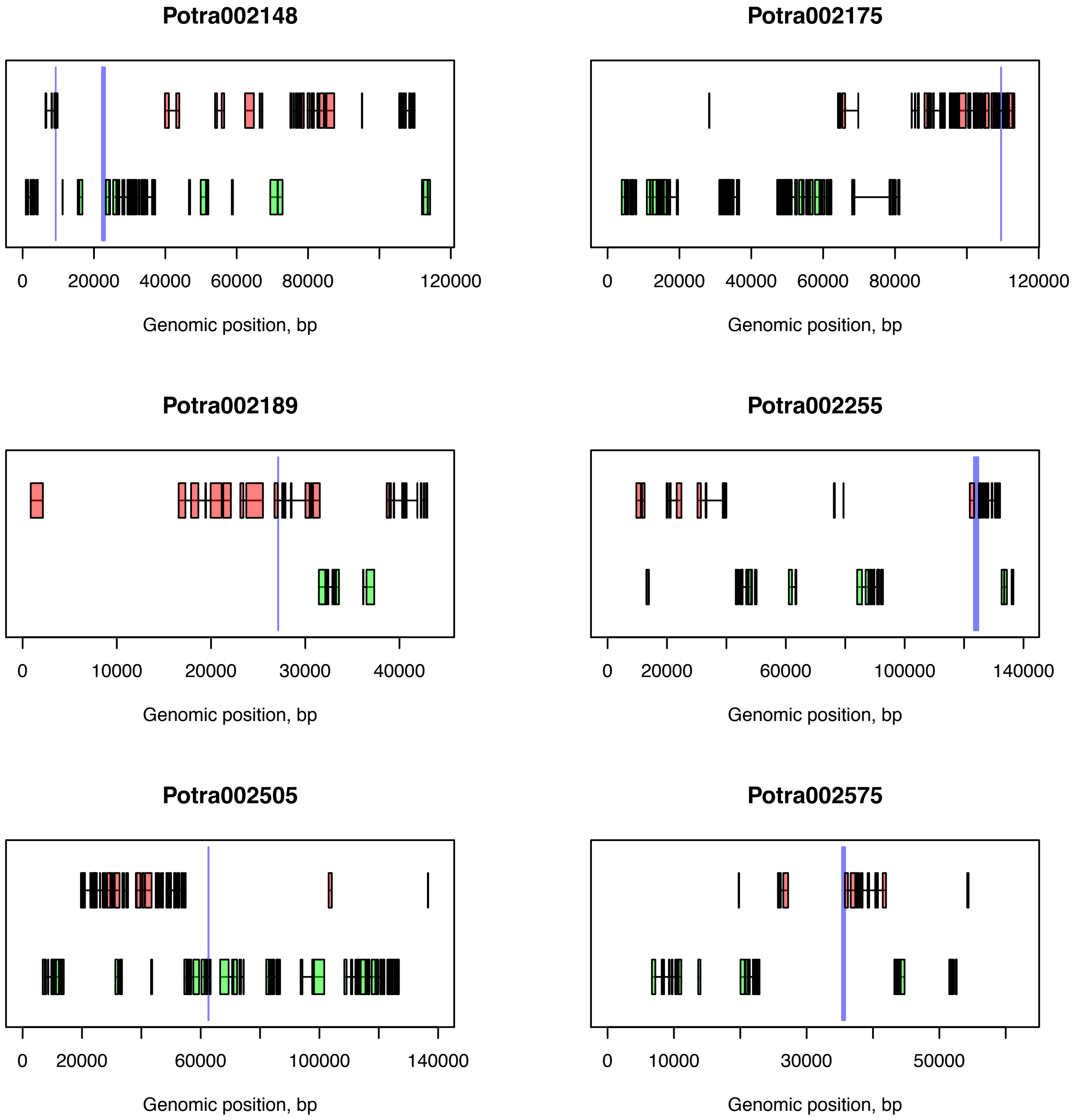
Potra002148
0
20000
40000
60000
80000
120000
Genomic position, bp
Potra002175
0
20000
40000
60000
80000
120000
Genomic position, bp
Potra002189
0
10000
20000
30000
40000
Genomic position, bp
Potra002255
0
20000
60000
100000
140000
Genomic position, bp
Potra002505
0
20000
60000
100000
140000
Genomic position, bp
Potra002575
0
10000
30000
50000
Genomic position, bp

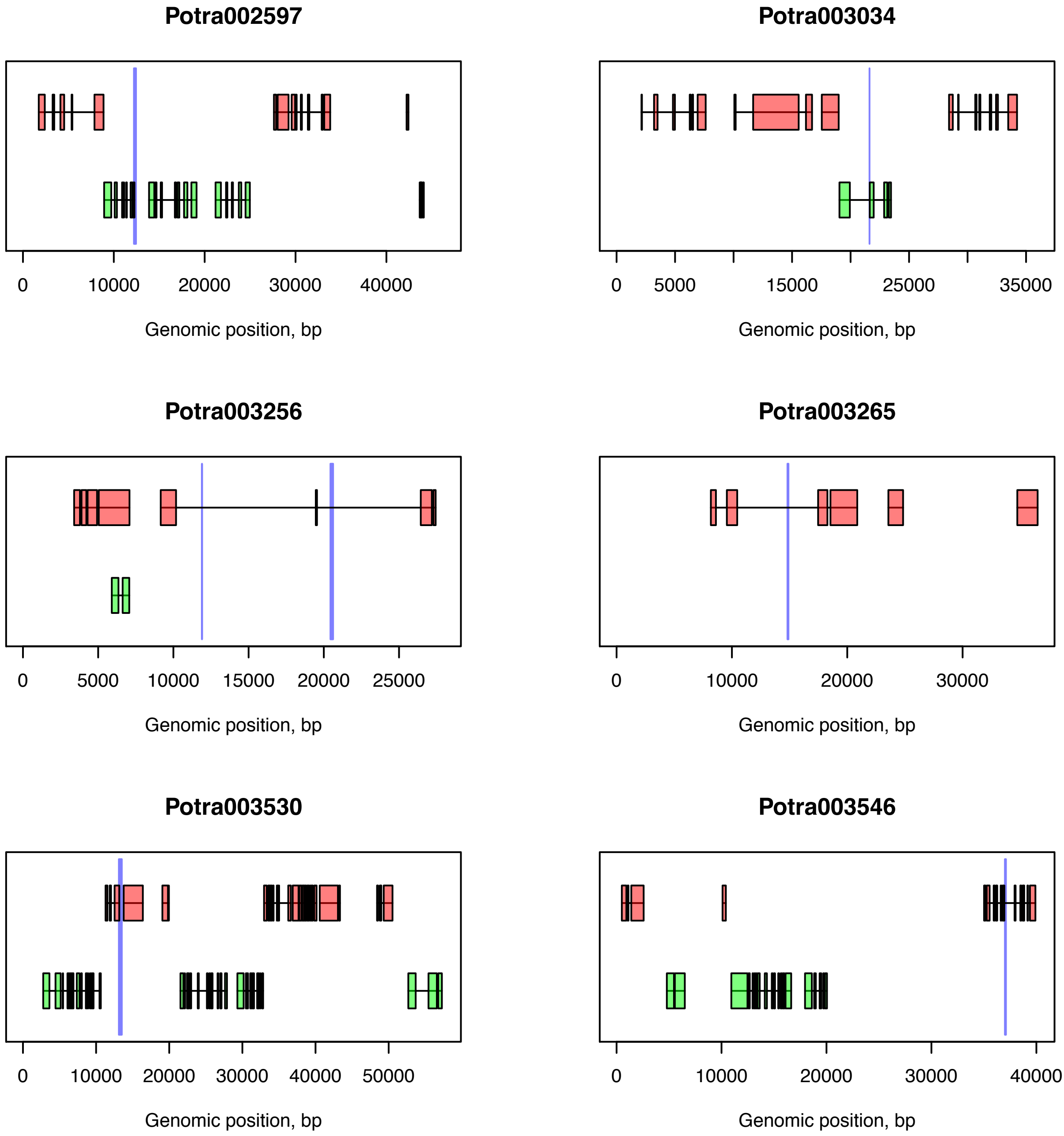
Potra002597
0
10000
20000
30000
40000
Genomic position, bp
Potra003034
0
5000
15000
25000
35000
Genomic position, bp
Potra003256
0
5000
10000
15000
20000
25000
Genomic position, bp
Potra003265
0
10000
20000
30000
Genomic position, bp
Potra003530
0
10000
20000
30000
40000
50000
Genomic position, bp
Potra003546
0
10000
20000
30000
40000
Genomic position, bp

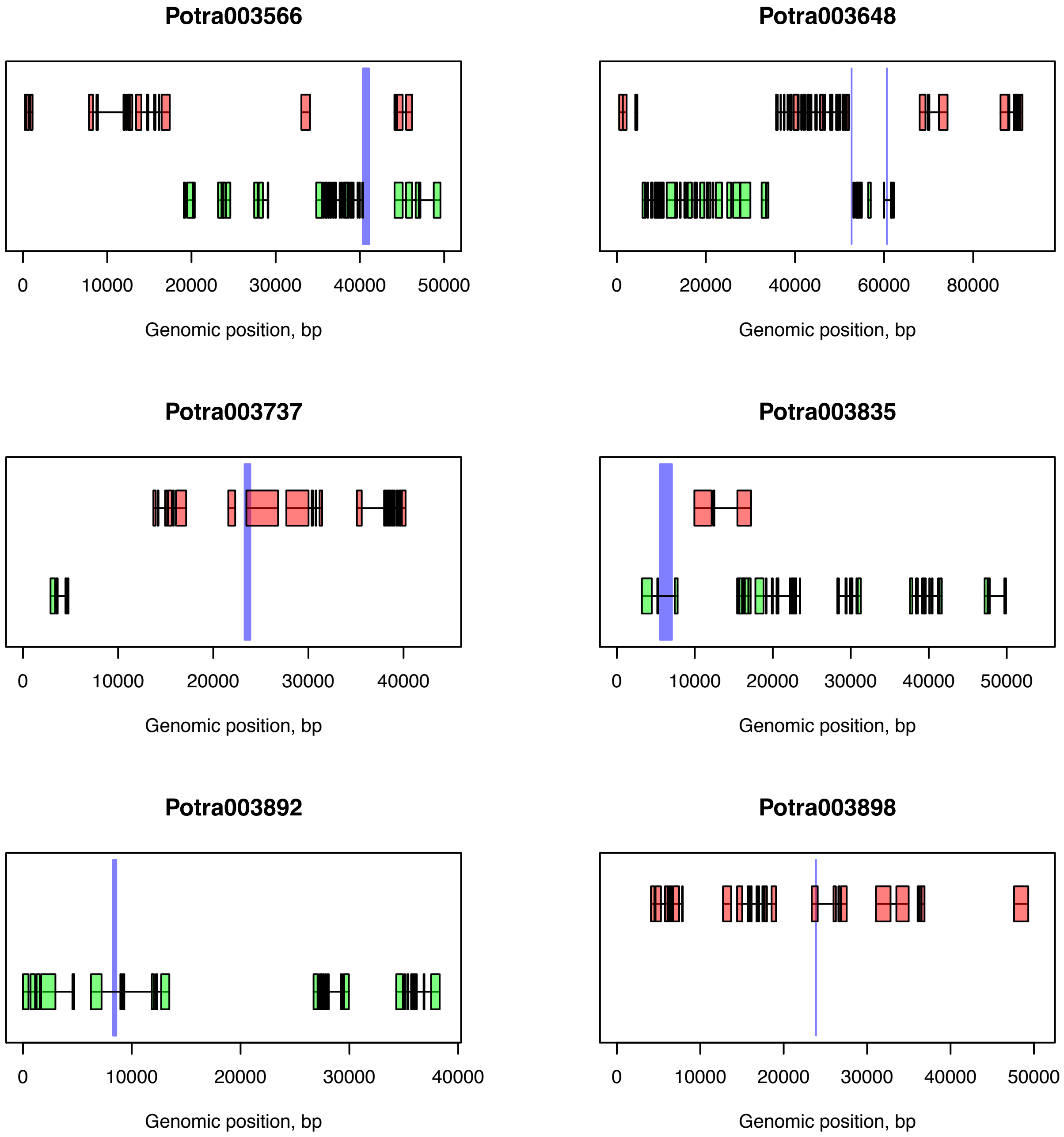

Potra003566
0
10000
20000
30000
40000
50000
Genomic position, bp
Potra003648
0
20000
40000
60000
80000
Genomic position, bp
Potra003737
0
10000
20000
30000
40000
Genomic position, bp
Potra003835
0
10000
20000
30000
40000
50000
Genomic position, bp
Potra003892
0
10000
20000
30000
40000
Genomic position, bp
Potra003898
0
10000
20000
30000
40000
50000
Genomic position, bp

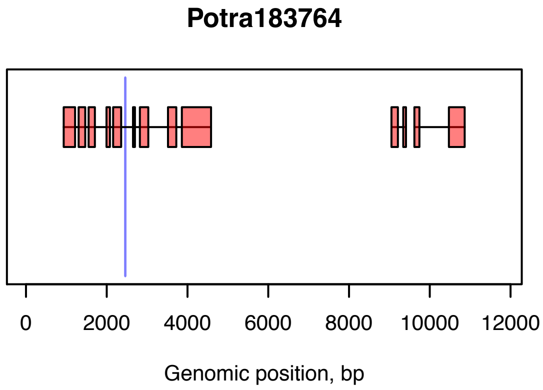
Potra183764
0
2000
4000
6000
8000
10000
12000
Genomic position, bp

Supplementary Figure 6

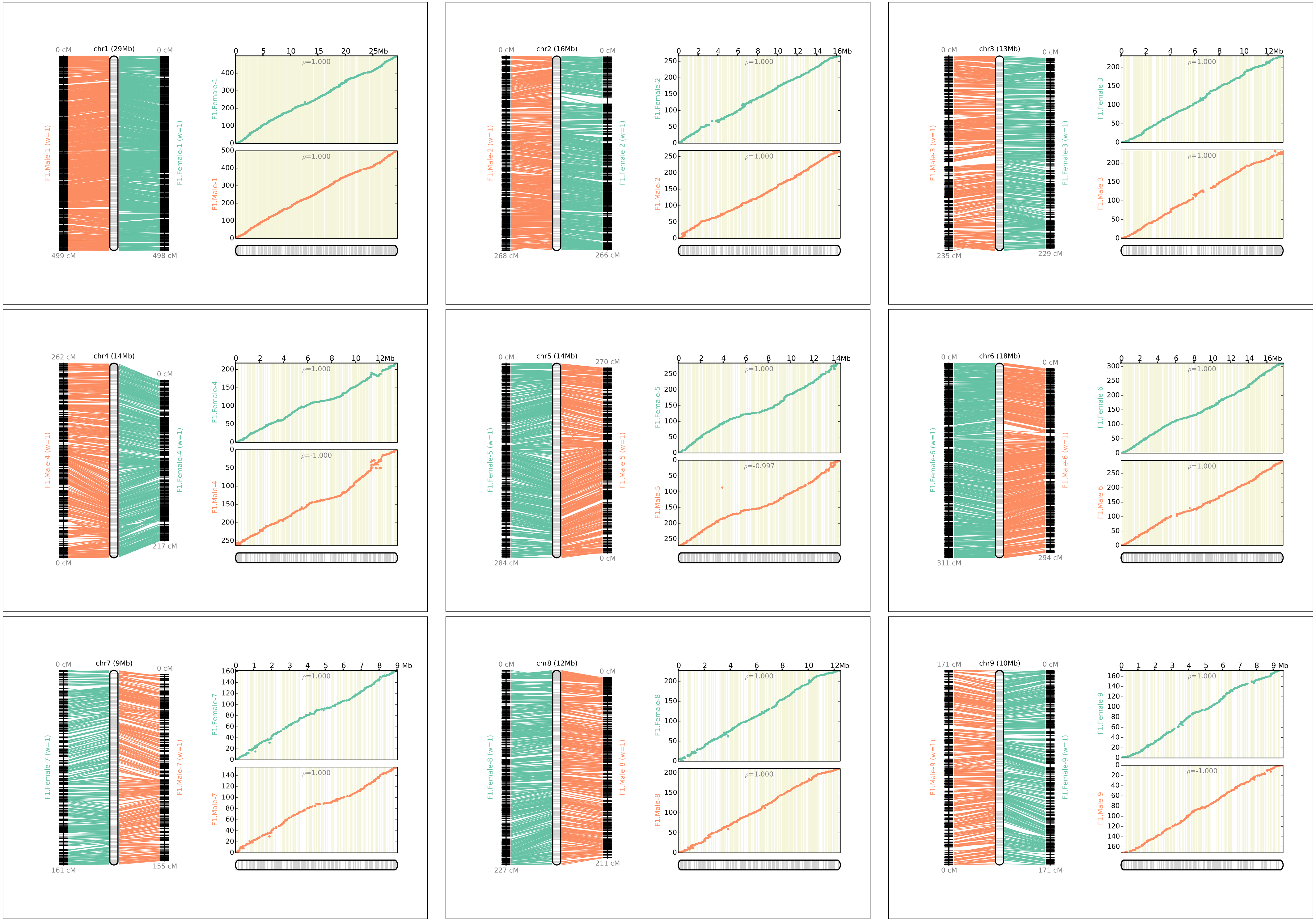
chr1 (29Mb)
0 cM
0 cM
499 cM
498 cM
F1.Male-1 (w=1)
F1.Female-1 (w=1)
0
5
10
15
20
25Mb
F1.Female-1
ρ=1.000
F1.Male-1
ρ=1.000
chr2 (16Mb)
0 cM
0 cM
268 cM
266 cM
F1.Male-2 (w=1)
F1.Female-2 (w=1)
0
2
4
6
8
10
12
14
16Mb
F1.Female-2
ρ=1.000
F1.Male-2
ρ=1.000
chr3 (13Mb)
0 cM
0 cM
235 cM
229 cM
F1.Male-3 (w=1)
F1.Female-3 (w=1)
0
2
4
6
8
10
12Mb
F1.Female-3
ρ=1.000
F1.Male-3
ρ=1.000
chr4 (14Mb)
262 cM
0 cM
0 cM
217 cM
F1.Male-4 (w=1)
F1.Female-4 (w=1)
0
2
4
6
8
10
12Mb
F1.Female-4
ρ=1.000
F1.Male-4
ρ=-1.000
chr5 (14Mb)
0 cM
270 cM
284 cM
0 cM
F1.Female-5 (w=1)
F1.Male-5 (w=1)
0
2
4
6
8
10
12
14Mb
F1.Female-5
ρ=1.000
F1.Male-5
ρ=-0.997
chr6 (18Mb)
0 cM
0 cM
311 cM
294 cM
F1.Female-6 (w=1)
F1.Male-6 (w=1)
0
2
4
6
8
10
12
14
16Mb
F1.Female-6
ρ=1.000
F1.Male-6
ρ=1.000
chr7 (9Mb)
0 cM
0 cM
161 cM
155 cM
F1.Female-7 (w=1)
F1.Male-7 (w=1)
0
1
2
3
4
5
6
7
8
9 Mb
F1.Female-7
ρ=1.000
F1.Male-7
ρ=1.000
chr8 (12Mb)
0 cM
0 cM
227 cM
211 cM
F1.Female-8 (w=1)
F1.Male-8 (w=1)
0
2
4
6
8
10
12Mb
F1.Female-8
ρ=1.000
F1.Male-8
ρ=1.000
chr9 (10Mb)
171 cM
0 cM
0 cM
171 cM
F1.Male-9 (w=1)
F1.Female-9 (w=1)
0
1
2
3
4
5
6
7
8
9 Mb
F1.Female-9
ρ=1.000
F1.Male-9
ρ=-1.000

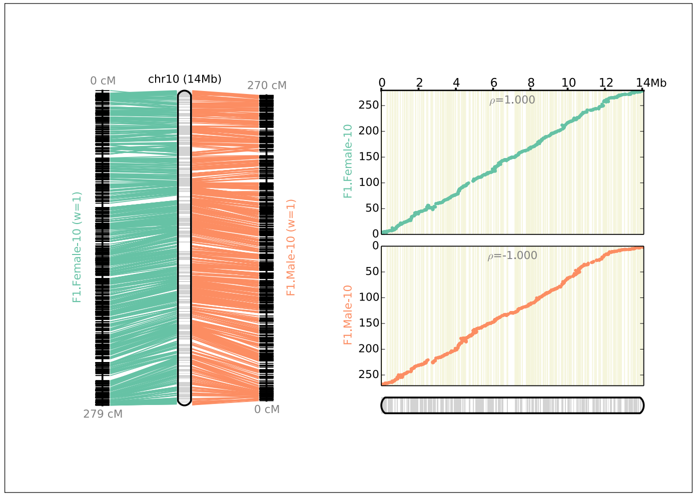
chr10 (14Mb)
0 cM
279 cM
270 cM
0 cM
F1.Female-10 (w=1)
F1.Male-10 (w=1)
F1.Female-10
F1.Male-10
ρ=1.000
ρ=-1.000
14Mb

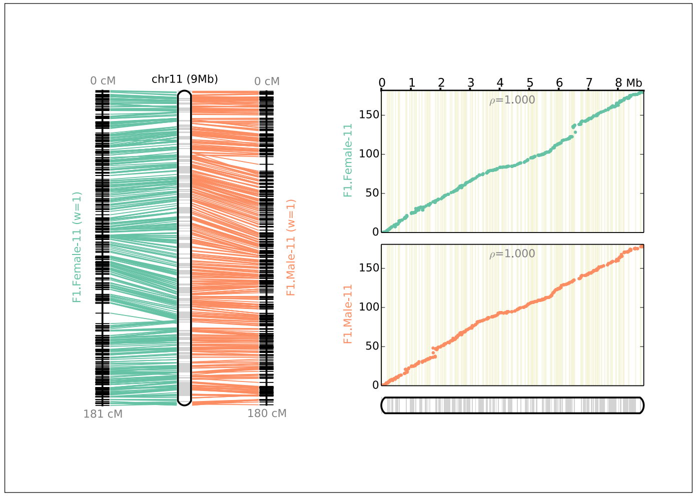
chr11 (9Mb)
0 cM
181 cM
0 cM
180 cM
F1.Female-11 (w=1)
F1.Male-11 (w=1)
F1.Female-11
F1.Male-11
ρ=1.000
ρ=1.000
8 Mb

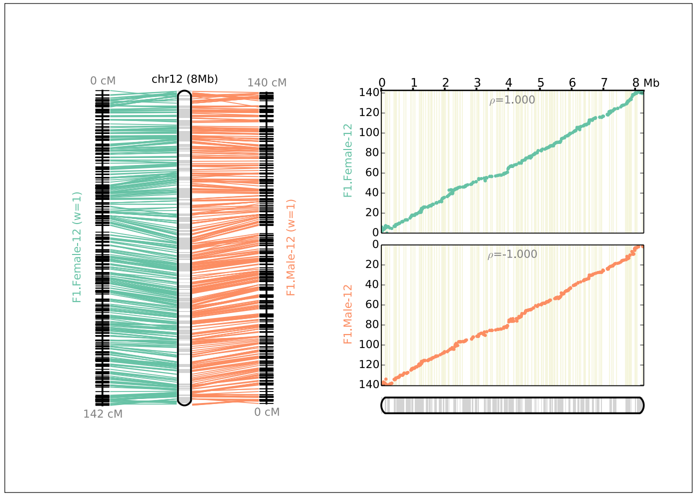
chr12 (8Mb)
0 cM
142 cM
140 cM
0 cM
F1.Female-12 (w=1)
F1.Male-12 (w=1)
F1.Female-12
F1.Male-12
ρ=1.000
ρ=-1.000
8 Mb

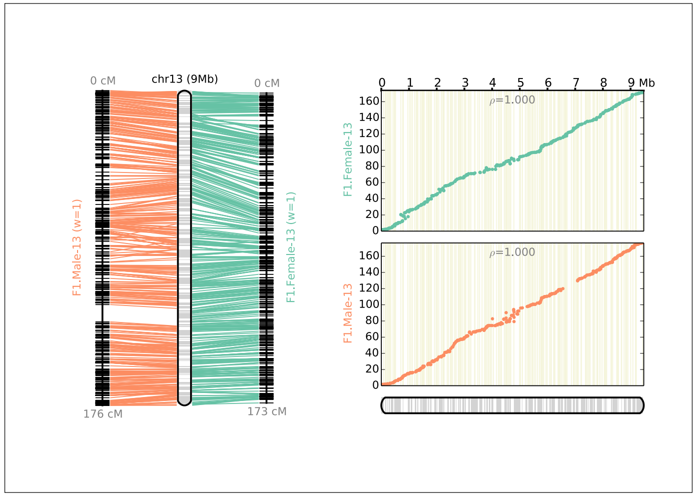
chr13 (9Mb)
0 cM
176 cM
0 cM
173 cM
F1.Male-13 (w=1)
F1.Female-13 (w=1)
F1.Female-13
F1.Male-13
ρ=1.000
ρ=1.000
9 Mb

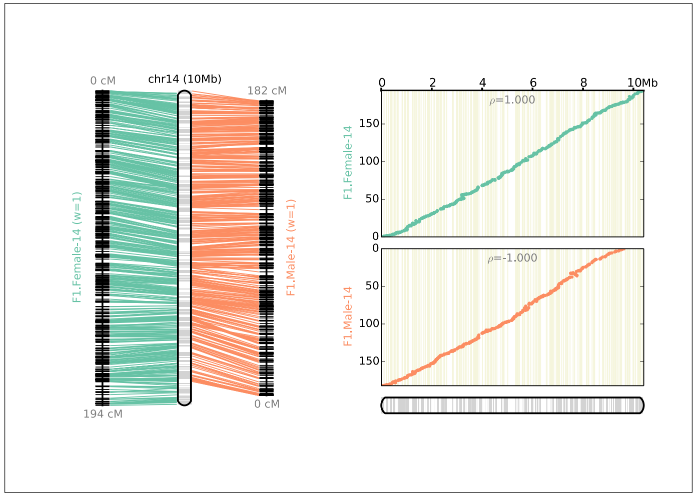
chr14 (10Mb)
0 cM
194 cM
182 cM
0 cM
F1.Female-14 (w=1)
F1.Male-14 (w=1)
F1.Female-14
F1.Male-14
ρ=1.000
ρ=-1.000
10Mb

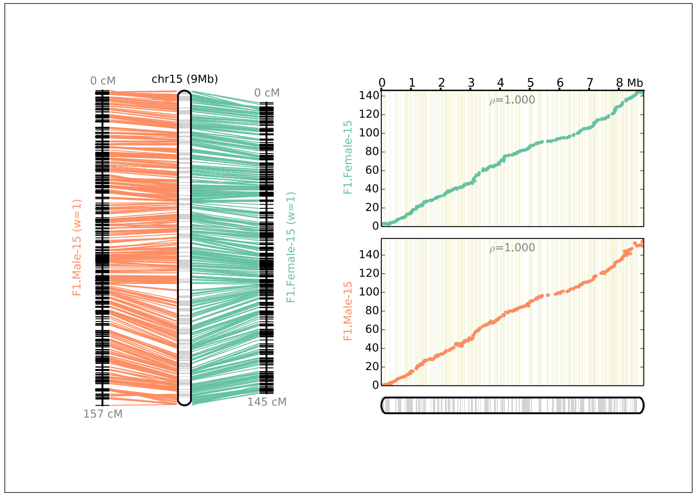
chr15 (9Mb)
0 cM
157 cM
0 cM
145 cM
F1.Male-15 (w=1)
F1.Female-15 (w=1)
F1.Female-15
F1.Male-15
ρ=1.000
ρ=1.000
8 Mb

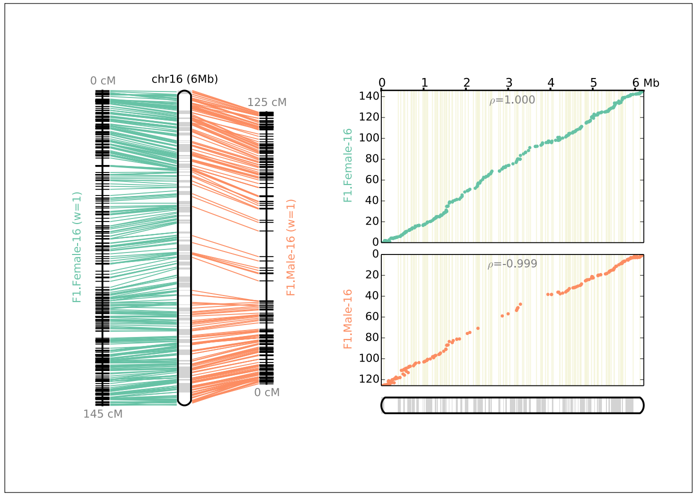
chr16 (6Mb)
0 cM
145 cM
125 cM
0 cM
F1.Female-16 (w=1)
F1.Male-16 (w=1)
F1.Female-16
F1.Male-16
ρ=1.000
ρ=-0.999
6 Mb

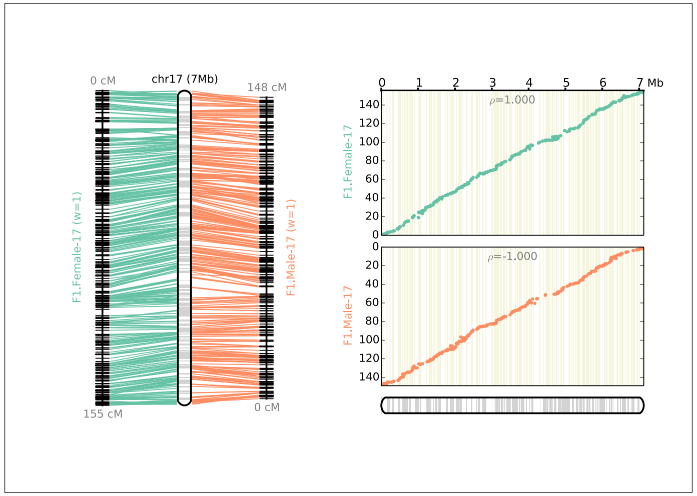
chr17 (7Mb)
0 cM
155 cM
148 cM
0 cM
F1.Female-17 (w=1)
F1.Male-17 (w=1)
F1.Female-17
F1.Male-17
ρ=1.000
ρ=-1.000
7 Mb

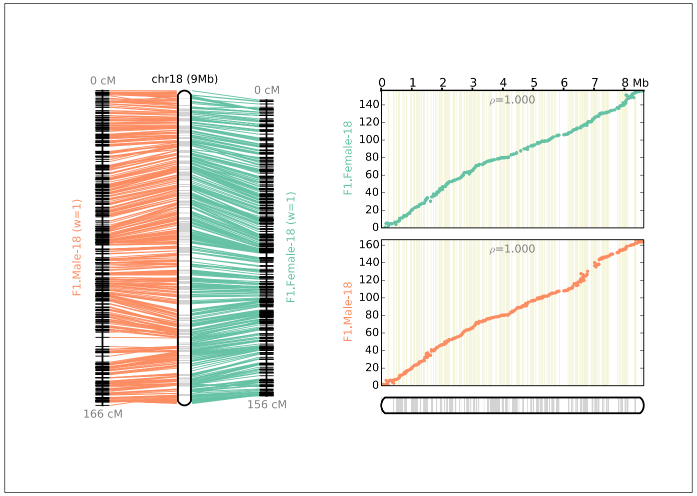
chr18 (9Mb)
0 cM
166 cM
0 cM
156 cM
F1.Male-18 (w=1)
F1.Female-18 (w=1)
F1.Female-18
F1.Male-18
ρ=1.000
ρ=1.000
8 Mb

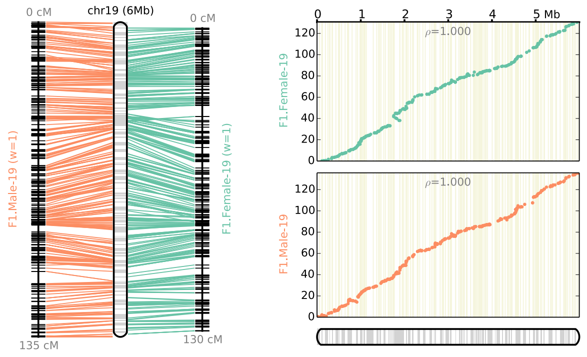

0 cM
chr19 (6Mb)
0 cM
F1.Male-19 (w=1)
F1.Female-19 (w=1)
135 cM
130 cM
0
1
2
3
4
5 Mb
ρ=1.000
F1.Female-19
F1.Male-19
0
20
40
60
80
100
120

Supplementary Figure 7

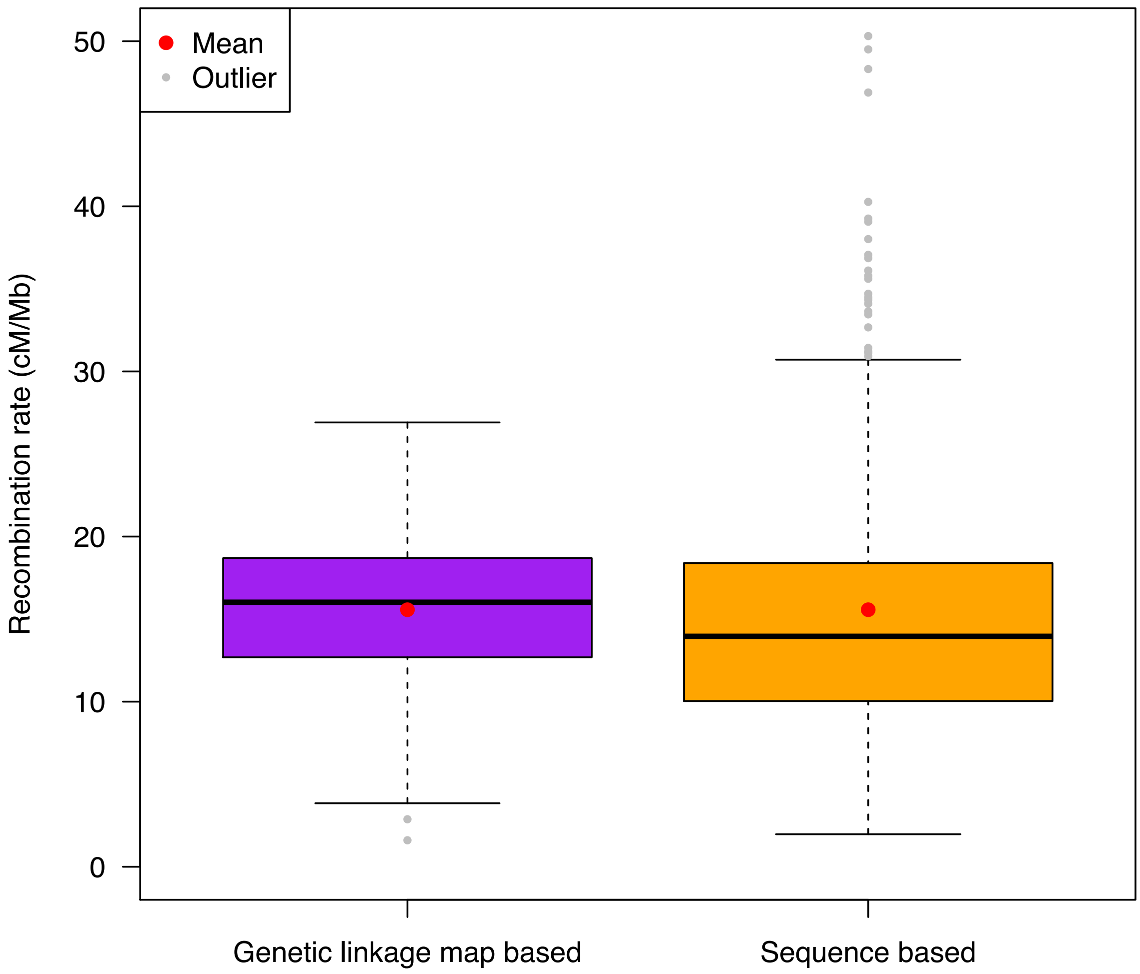

Supplementary Figure 8

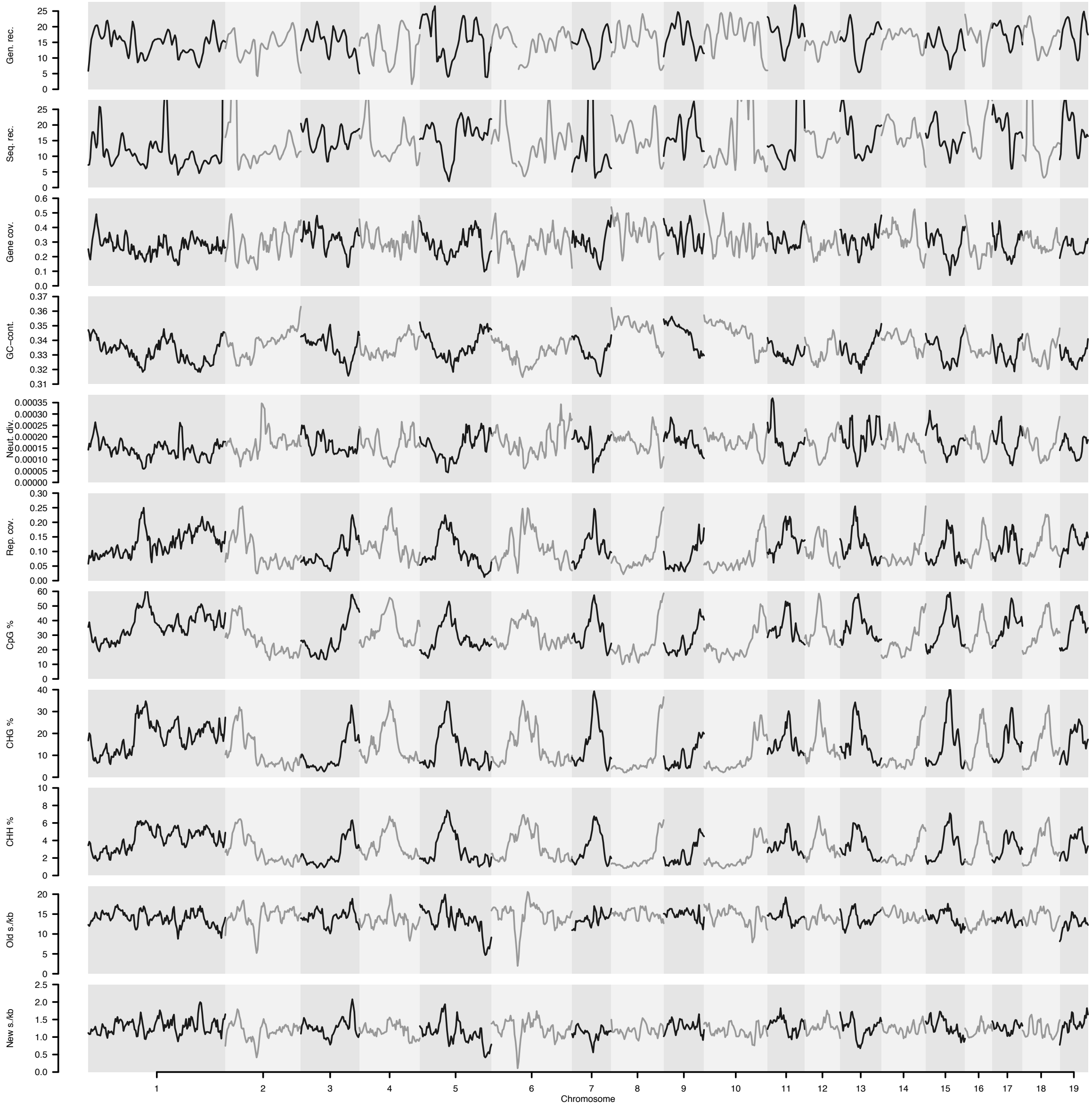

Supplementary Figure 9

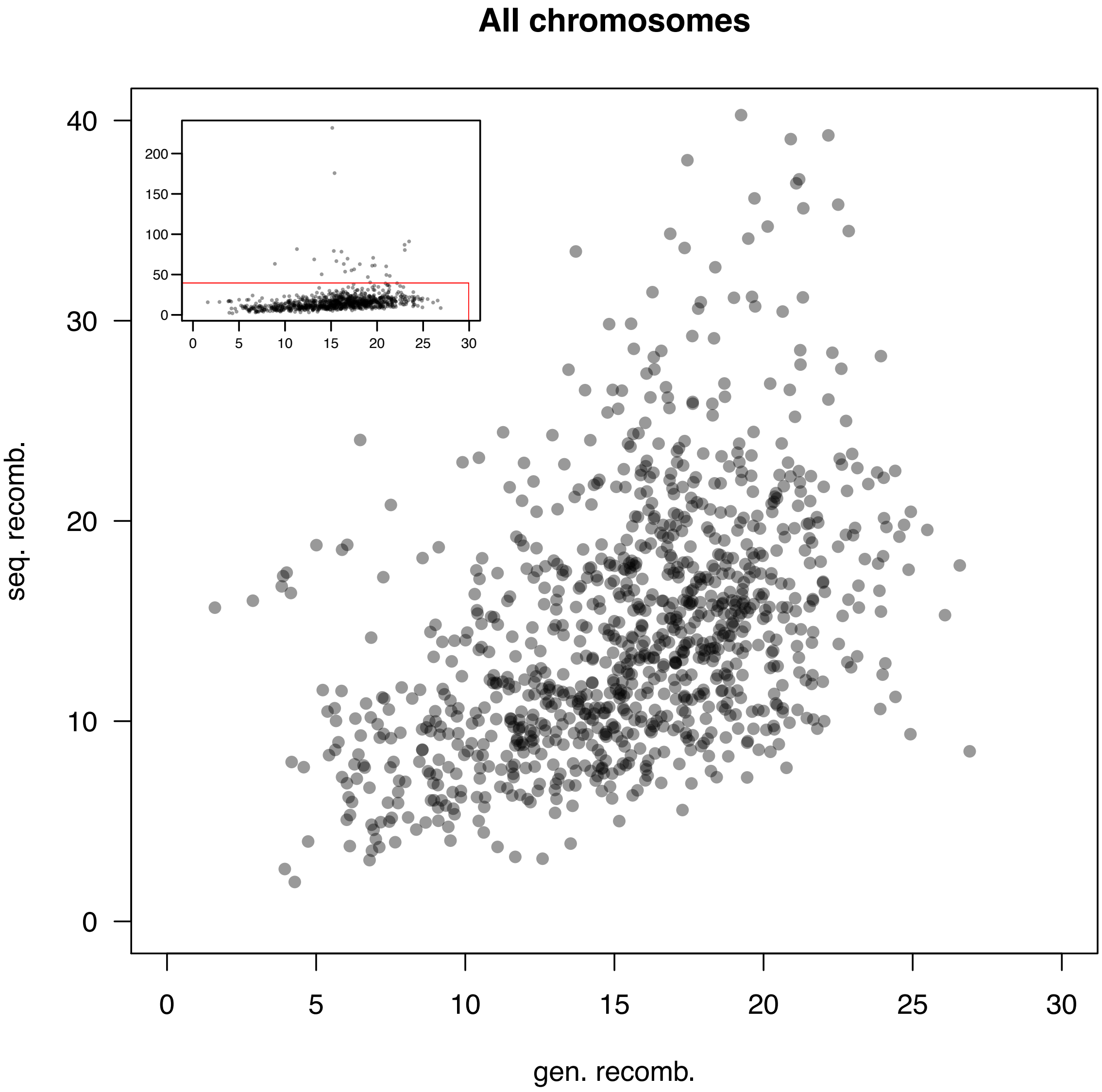
