## Supplementary Tables for "Constructing a high-density linkage map to infer the genomic landscape of recombination rate variation in European Aspen *(Populus tremula)*"

Supplementary Table 1

| | Female map | Male map |
|---|---|---|
| Linkage Groups | 19 | 19 |
| Markers (unique) | 14596 | 13996 |
| Markers per Mb | 69.3 | 68.3 |
| N50 Scaffolds | 1961 | 1911 |
| Scaffolds | 4184 | 4011 |
| Scaffolds with 1 marker | 1320 | 1294 |
| Scaffolds with 2 markers | 969 | 855 |
| Scaffolds with 3 markers | 502 | 525 |
| Scaffolds with >=4 markers | 1393 | 1337 |
| Total bases | 210,660,926 (54,6%) | 205,055,562 (53,1%) |

**Supplementary Table 2**

| | Map based | LD based |
|---|---|---|
| Min. | 1.605 | 1.969 |
| 1st Quantile | 12.685 | 10.039 |
| Median | 16.022 | 13.958 |
| Mean | 15.570 | 16.100 |
| 3rd Quantile | 18.692 | 18.375 |
| Max. | 26.911 | 231.801 |
